# Temporal N-terminomics analysis reveals proteome remodeling following brensocatib-mediated cathepsin C inhibition in promyeloblast cells

**DOI:** 10.64898/2026.09.03.748759

**Authors:** Yimin Zhu, Stephanie Luedtke, Bangyan Xu, Alexander R. Ziegler, Kai Qi Yek, Nichollas E. Scott, Elena K. Schneider-Futschik, Laura E. Edgington-Mitchell

## Abstract

Cathepsin C (CatC) is known to activate neutrophil serine proteases (NSPs) involved in innate immune function, although its broader impact on cellular proteolytic networks remains poorly defined. Here, we characterized the proteolytic landscape of HL-60 neutrophil progenitor cells following treatment with the CatC inhibitor brensocatib. Activity-based probes confirmed sustained inhibition of CatC by brensocatib, accompanied by progressive suppression of downstream elastase-like protease activation over 16 hours, 72 hours, and 7 days. An enrichment-free N-terminomics workflow was used to compare control and brensocatib-treated HL-60 cells. Following prolonged CatC inhibition, NSPs were reduced in abundance, while lysosomal cathepsins and endogenous protease inhibitors increased. These findings are consistent with remodeling of the protease-antiprotease network. Cleavage site analysis identified a pronounced, time-dependent, NSP-associated P1 cleavage signature enriched for Val, Thr, Ala, Ile, and Cys in control cells that was progressively lost following brensocatib treatment. We also identified established and candidate CatC-dependent cleavage events, together with proteolytic adaptations that emerged in the absence of CatC activity. Collectively, these findings demonstrate that CatC inhibition extends beyond suppression of canonical NSP activation to progressive remodeling of the broader protease-antiprotease network, providing new mechanistic insight into the cellular consequences of therapeutic CatC inhibition.

**Graphical Abstract:** 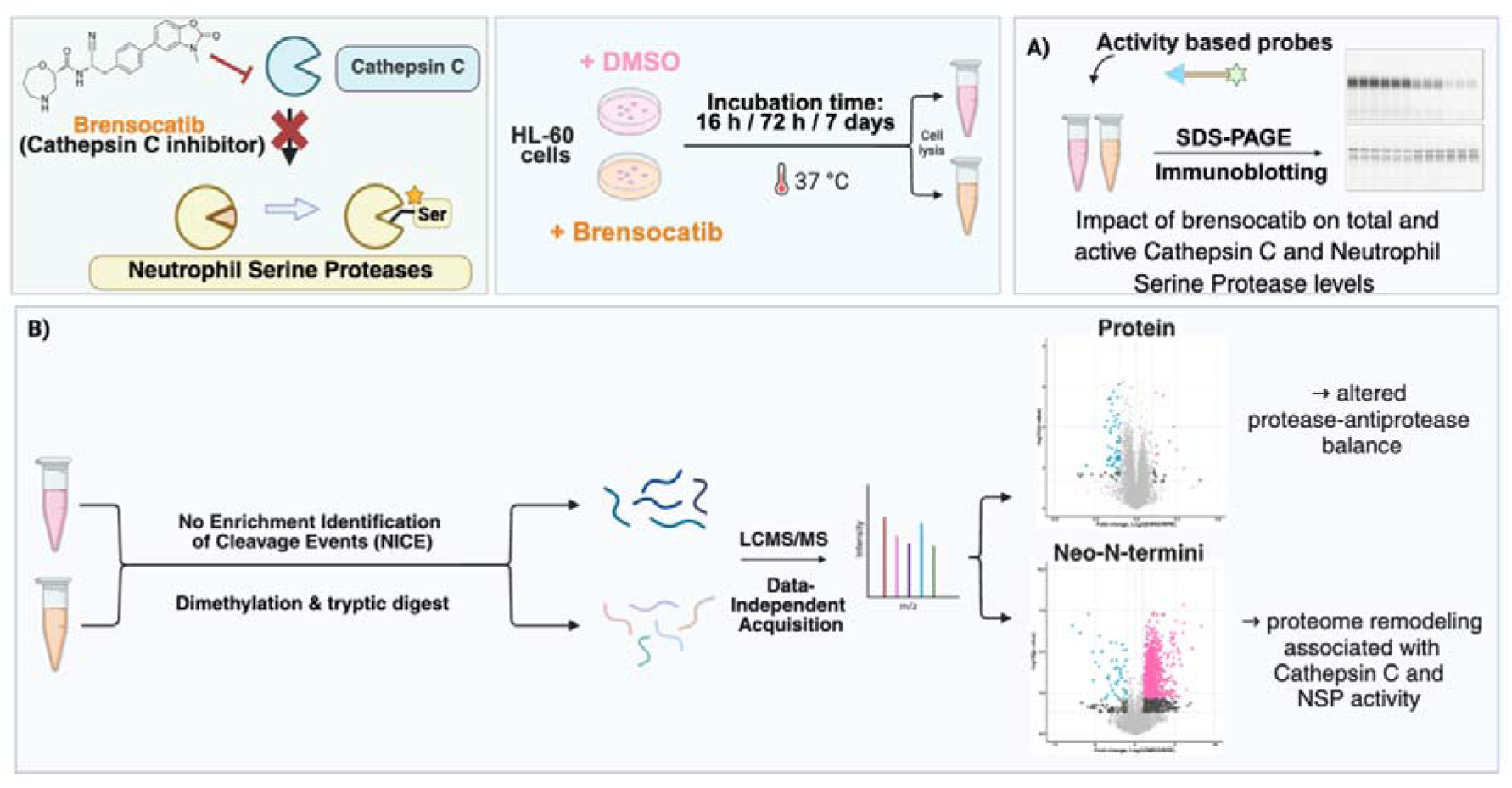

## Introduction

Cathepsin C (CatC), also known as dipeptidyl peptidase 1 (DPP-1), is a ubiquitously expressed lysosomal cysteine protease that functions as the primary activator of neutrophil serine proteases (NSPs) (1–3). CatC is synthesized as an inactive zymogen and trafficked to the endolysosomes primarily via the mannose-6-phosphate pathway, where proteolytic processing by other cathepsins produces the mature, tetrameric enzyme (4, 5). During promyelocytic stages of neutrophil maturation, CatC functions as a dipeptidyl aminopeptidase to remove two residues from the proform of the NSPs, including neutrophil elastase (NE), proteinase 3 (PR3), cathepsin G (CatG), and neutrophil serine protease 4 (NSP4) (6). This cleavage step is required for their enzymatic activation and downstream contribution to host defense (7–9).

While NSP activity is essential for innate immunity, elevated activation contributes to tissue damage and chronic inflammation (10, 11). Excessive NSP activity is associated with the severity of neutrophil-dominated lung diseases, including bronchiectasis, cystic fibrosis, and chronic obstructive pulmonary disease (12–15), where proteases degrade extracellular matrix proteins (16), inactivate endogenous protease inhibitors (17), and impair mucociliary function (18, 19). CatC has consequently emerged as an attractive therapeutic target whereby inhibiting a single upstream protease prevents activation of multiple downstream NSPs.

This strategy resulted in the 2025 FDA approval of the CatC inhibitor brensocatib as the first targeted therapeutic treatment for bronchiectasis patients (20), with several other inhibitors in the clinical pipeline (21–23). Prior to brensocatib, treatment of bronchiectasis primarily focused on symptom management, including the use of mucolytics and antibiotics (24, 25), without targeting the persistent neutrophilic inflammation that drives disease progression. By preventing NSP activation, CatC inhibition provides an approach to restore the protease-antiprotease balance and limit severe exacerbations associated with tissue damage. Phase 3 clinical trials demonstrated that brensocatib significantly reduced the annualized rate of pulmonary exacerbations compared with placebo (20).

Despite this therapeutic success, the broader molecular consequences of CatC inhibition remain poorly understood. Its effects on global protein abundance, proteolytic processing, and downstream protease networks have not been systematically studied. Defining these changes would improve our understanding of its functions in both normal physiology and the broader consequences of therapeutic CatC targeting. Herein, we have applied activity-based probes in combination with quantitative proteomics and degradomics methods to characterize the impact of pharmacological inhibition of CatC in human promyeloblast HL-60 cells. By integrating measurements of protease activity, total protein abundance, and proteolytic processing, we define the temporal effects of brensocatib on the intracellular proteolytic landscape, identify candidate CatC-dependent cleavage events, and reveal proteolytic adaptations that emerge following sustained CatC inhibition.

## Methods

### Cell culture

HL-60 cells (ATCC, CCL-240) were cultured in Iscove’s modified Dulbecco’s medium (IMDM, ATCC, 30-2005) supplemented with 20% (v/v) fetal bovine serum (FBS, CellSera) and 1% antibiotics (100 U/ml penicillin-streptomycin, Thermo Fisher Scientific). The cells were maintained in a humidified atmosphere at 37°C with 5% CO_2_ and sub-cultured to ensure cell density remained below 1.5 × 10^6^ cells/mL.

### Concentration-response assay of Cathepsin C in HL-60 cells for 16 h and 72 h

HL-60 cells were seeded at 1.5 × 10^6^ or 1 × 10^6^ cells/mL for 16 h or 72 h cultures, respectively. Brensocatib (MedChemExpress, HY-101056) was dissolved in DMSO and serially diluted to 1000x stocks (20 mM, 10 mM, 5 mM, 1 mM and 0.2 mM) before being added directly to cell culture media at a final DMSO concentration of 0.1% (v/v). Cells were incubated with brensocatib or DMSO for the indicated time before harvesting.

### Time course of brensocatib treatment in HL-60 cells

HL-60 cells (n = 3/group) were treated with brensocatib (20 μM) or an equivalent volume of DMSO (0.1% v/v) for 16 h or 72 h before harvesting. For 7-day continuous inhibition of CatC, HL-60 cells were seeded at 1 × 10^5^ cells/mL and treated with brensocatib (20 μM), replenshing drug-containing media every three days.

### Monitoring the impact of brensocatib on protease activity

After incubation with brensocatib or DMSO, cells were harvested at the indicated timepoints and lysed in either citrate buffer (50 mM citrate [Thermo Fisher Scientific, pH 5.5], 0.5% CHAPS [Sigma], 0.1% Triton X-100, 4 mM DTT [Sigma]) or phosphate-buffered saline (PBS) with 0.1% Triton X-100 buffer. The lysates were clarified by centrifugation for 7 min at max speed at 4°C. Protein concentration was measured by BCA assay (Pierce), and 40-80 μg total protein was aliquoted into a final volume of 20 μL lysis buffer. Lysates were incubated at 37°C for 20 min with the activity-based probes FY01 (26), PK105b (27) or BMV109 (28) at a final concentration of 1 μM (1% DMSO). Sample buffer (5x: 50% glycerol, 250 mM Tris-Cl, pH 6.8, 10% SDS, 6.25% beta-mercaptoethanol, 0.04% bromophenol blue) was added at 1x to quench the reaction. Samples were boiled at 95°C for 5 min, and proteins were resolved on a freshly prepared 15% SDS-PAGE gel. Probe labeling was visualized by scanning for Cy5 fluorescence on a Typhoon 5 flatbed laser scanner (GE Healthcare). Proteins were transferred onto nitrocellulose membranes using a Trans-Blot Turbo Transfer System (BioRad) for immunoblotting as below.

### Immunoblotting

Primary antibodies were diluted in blocking buffer (1:1 PBS containing 0.1% Tween-20 [Sigma; PBS-T] and Intercept blocking buffer [Li-Cor]) and incubated with nitrocellulose membranes on an orbital shaker overnight at 4°C, followed by 3 washes with PBS-T for 5 min. Secondary antibodies were incubated in blocking buffer at room temperature for 1 h and washed 3 times with PBS-T for 5 min before scanning. IRDye-800-conjugated antibodies were scanned for fluorescence via Typhoon 5. Horseradish peroxidase (HRP)-conjugated antibodies were detected by chemiluminescence using Pierce ECL Western blotting substrate (Thermo Fisher Scientific, PI32106) on a Chemidoc MP imager (Bio-Rad). Antibodies used in the study include CatC (1:1000, R&D, AF1034), NE (1:1000, R&D, AF4517), PR3 (1:1000, R&D, AF6134), Cathepsin X (1:1000, R&D, AF1033), Cathepsin B (1:1000, R&D, AF965), beta-actin (1:10000, Sigma A5060), donkey anti-goat IRDye-800 (1:10000, Li-Cor; 926-32214) and goat anti-rabbit-HRP (1:10000, Invitrogen, A16035).

### Immunoprecipitation

Probe-labeled lysates (60-80 μg total protein) were aliquoted into input and pulldown samples. 500 μL of immunoprecipitation (IP) buffer (PBS, pH 7.4, 0.5% Nonidet P-40 Substitute [Sigma], 1 mM EDTA) was added to the pulldown samples followed by pre-washed protein A/G agarose beads (40 µl slurry; Santa Cruz Biotechnology). Antibodies used for the pulldowns (CatC [R&D, AF1034], NE [R&D, AF4517], Cathepsin S (CatS) [R&D, AF1183], Cathepsin L (CatL) [R&D, AF1515]and PR3 [Thermo Fisher Scientific, PA5-85928]) were added (10 μL per pulldown) and the tubes were rotated overnight at 4°C. Pulldown samples were washed four times in IP buffer followed by a final wash in 0.9% NaCl. Proteins were eluted from beads by boiling in 2x sample buffer (20 µl) for 5 min at 95°C and loaded onto a 15% SDS-PAGE gel alongside input samples. Probe labeling was visualized by scanning for Cy5 fluorescence on a Typhoon 5.

### No-enrichment Identification of Cleavage Events (NICE) analysis

HL-60 cells (n = 4/group) were plated with either 0.1% DMSO or 20 μM brensocatib and incubated for 16-h, 72-h, or 7 days as above. Cells were harvested and then lysed by sonication in lysis buffer containing 4% SDS, 50 mM HEPES (pH 7.5, Sigma), and Roche cOmplete, EDTA-free protease inhibitor (Sigma). Total protein concentration in the samples was determined by BCA, and 100 µg protein was aliquoted and stored at −20°C prior to proteomics sample preparation.

Sample preparation for NICE was performed according to the previously reported method (29, 30). Proteins were first reduced with dithiothreitol (DTT; 20 mM final, 10 min, 80°C, 500 rpm) and alkylated using iodoacetamide (50 mM, 30 min, 37°C, 500 rpm). The reaction was quenched by addition of excess DTT (50mM final, 20 min, 37°C, 500 rpm). Pre-washed Sera-Mag™ magnetic Speedbeads (Cytiva, 45152105050250 and 65152105050250 in 1:1 ratio, with 1:20 protein-to-bead ratio) were added to the sample and incubated (20 min, 25°C,1000 rpm) prior to 3 washes using 80% ethanol and then resuspended in 200 mM HEPES. Dimethylation was initiated by addition of formaldehyde (30 mM) and sodium cyanoborohydride (30 mM) for 1 h at 37°C, 1000 rpm. The labeling step was repeated once more to ensure complete labeling and then quenched by the addition of 25 µl Tris (4M, pH 6.8, 1 h, 37°C, 1000 rpm). Labeled samples were cleaned up by the addition of SP3 beads (1:10 protein-to-bead ratio) and three washes with 80% ethanol. Proteins were digested overnight at 37°C with trypsin (Merck) in 200 mM HEPES with the resulting peptides collected and cleaned up using C18 StageTips (Empore, 3M) with the addition of Oligo R3 reversed-phase resin material (Thermo Fisher Scientific) according to the previously described method (31). Peptides were mixed with Buffer A* (2% acetonitrile, 0.1% trifluoroacetic acid), loaded onto conditioned StageTip columns, washed with Buffer A* before being eluted with Buffer B (80% acetonitrile, 0.1% formic acid) and dried with a SpeedVac.

### LC-MS/MS analysis

Samples were reconstituted in 25 µL Buffer A* prior to LC-MS/MS analysis. Peptides were loaded onto a two-column configuration consisting of the Acclaim Pepmap nanotrap enrichment column (C18, 100□Å, 75□µm × 2□cm) and the high-throughput μPAC™ Neo HPLC Column (C18, 100□300 Å, 75□µm × 5.5□cm) on the Ultimate 3000 RSLC nano-flow reversed-phase HPLC system (Dionex). Samples were loaded onto the nanotrap enrichment column at a flow rate of 5 µL/min for 6 min in Buffer A (0.1% formic acid, 2% DMSO), before altering to 0.3 µL/min at 3% buffer B (0.1% formic acid, 77.9% acetonitrile, 2% DMSO) for 1 min, then increasing to 4% Buffer over 1 min, and from 4% to 80% for 75 min. The composition was held at 80% B for 3 min before dropping to 3% B within 0.1 min and maintained at 3% for 4.9 min. Each run was 95 min total. Full MS1 spectra were acquired using Orbitrap Exploris 480 followed by data-independent acquisition of the precursors on the Orbitrap MS/MS scans. MS1 event (120k resolution, normalized automated gain control (AGC) target at 250%, scan range 350-1400 m/z) and 50 MS2 events (HCD collision energy 30%, 30k resolution, normalized AGC at 2000%, 200-2000 m/z) of 13.7 m/z from 360.5 m/z to 1046.5 m/z were collected. The 7-day treated cell samples were run on a separate setup using the VanquishNeo2 UHPLC coupled to the Orbitrap Astral mass spectrometer, with the same column configuration. Each sample was analyzed using a 30 min analytical run, with the Orbitrap MS scan (resolution of 120k, normalized AGC target at 500%, 380-980 m/z) followed by 300 Astral MS2 events (precursor isolation window of 2 m/z, normalized AGC target at 500%, scan range 150-2000 m/z, HCD collision energies of 27%) collected from 380.422 m/z to 980.695 m/z. Each 30 min run was initiated at 3% Buffer B at 0.75 µL/min (0.1% formic acid, 77.9% acetonitrile, 2% DMSO). The Buffer B gradient was increased to 6% by 3 min, 23.5% by 23 min, 40% by 26.7 min, and further increased to 50% for 2 min before reaching 99% with a. 2 µL/min flow rate after 0.1 min and maintaining this until 30 min.

### Quantitative proteomics and N-terminomics analyses

Data files were processed and searched using FragPipe (v22.0) (32) against the human proteome (UniProt Accession: UP000005640), with a 50% reverse decoy database. For N-terminomics analysis, cleavage specificity was set to “TrypsinR” (Arg-C) and “SEMI-N_TERM”, with a maximum of one missed cleavage. Variable modifications were added including methionine oxidation (+15.9949 Da), acetylation at N-terminus (+42.0106 Da), N-terminal cyclization of Q/E (−18.0106 /-17.0625 Da), dimethylation at N-terminus (+28.0313 Da), lysine dimethylation (+28.0313 Da), and N-terminal lysine double dimethylation (+56.0626 Da). Cysteine carbamidomethylation (+57.02146 Da) was also set as a fixed modification. All other parameters were set to default. Quantitative analysis of the output was performed in Perseus (v1.6.0.7) (33). Data was log_2_ transformed, filtered for valid values by keeping those with at least three values in at least one group and missing values were imputed at random using a downshifted normal distribution (σ-width = 0.3, σ-downshift = −1.8) based the observed ion intensities. Peptide-level data was further processed in R by first normalizing to the protein-level abundance. Peptides with unlabeled lysine side chains were excluded from the analysis to minimize false positives associated with mislocalization of the dimethylation to the N-terminus. C-terminomic analysis of the 7-day dataset was performed using the same search parameters as above, except with the cleavage specificity set to “SEMI” to search for non-tryptic C-terminal peptides. Downstream filtering of C-terminal peptides was performed in R to exclude tryptic peptides, which end in arginine or unmodified lysine. Consensus sequence analysis was performed using pLogo, with all dimethylated N-terminal peptides detected in the corresponding experiment used as the background (34). Gene ontology (GO) pathway analysis was performed in R using enrichGO with the background defined as either the total proteins or total N-terminal peptides detected in each respective experiment (35).

### Statistical analysis

For probe and immunoblotting experiments, statistical analysis was performed using Prism (GraphPad Software, Boston, MA). Comparison between multiple experiment groups was performed using one-way ANOVA unless specified. All data are presented as mean ± SEM. For proteomics experiments, the mean differences of protein and peptide abundances between DMSO- and brensocatib-treated groups were compared using Student’s *t*-test. Volcano plots were generated using a threshold of –log_10_ (raw *p*-value) ≥ 1.3 (*p* < 0.05) and |log_2_(DMSO/Bre)| > 1. Multiple hypothesis testing correction was performed in Perseus using permutation-based false discovery rate (FDR) analysis, with resulting q-values reported in the Supplementary Information. Analyses and data visualization were performed in R (version 2025.5.0.496).

## Results & Discussion

### Time-dependent suppression of NSP activation after brensocatib-mediated cathepsin C inhibition in HL-60 cells

Although CatC is well recognized as the activator of NSPs, its broader substrate repertoire and the cellular consequences of its inhibition on the proteome remain largely unexplored. To establish a cellular model of sustained CatC inhibition for proteomic and degradomics analysis, we first characterized the time- and concentration-dependent effects of brensocatib (**Supplementary Fig. 1a**) on CatC and downstream NSP activity in HL-60 promyeloblasts. HL-60 cells express active CatC and have been previously used to assess its processing (5, 36). We first determined the brensocatib concentration required to achieve sustained CatC inhibition and suppression of downstream NSP activation.

HL-60 cells were treated with 0, 0.2, 1, 5, 10, or 20 μM of brensocatib for 16 h, and residual CatC activity was measured using the fluorescent activity-based probe FY01 (**Supplementary Fig. 1b**) (26). FY01 is a covalent vinyl sulfone-based probe known to target CatC, although non-specifically (37). At 16 h, labeling of a ∼23 kDa species, corresponding to the active CatC heavy chain, was reduced by brensocatib in a concentration-dependent manner (**Fig. 1a,b**). Immunoprecipitation (IP) of the FY01-labeled lysates with a CatC antibody confirmed that the 23 kDa band contains active CatC, and low levels of cathepsin S and L (**Fig 1d**). No FY01-labeled CatC was immunopurified after brensocatib treatment, confirming suppression of its activity (**Fig 1e**). By contrast, total CatC levels, as assessed by immunoblot, trended towards an increase with increasing drug concentrations, although this did not reach statistical significance (**Fig. 1a,c**).

**Figure 1.**
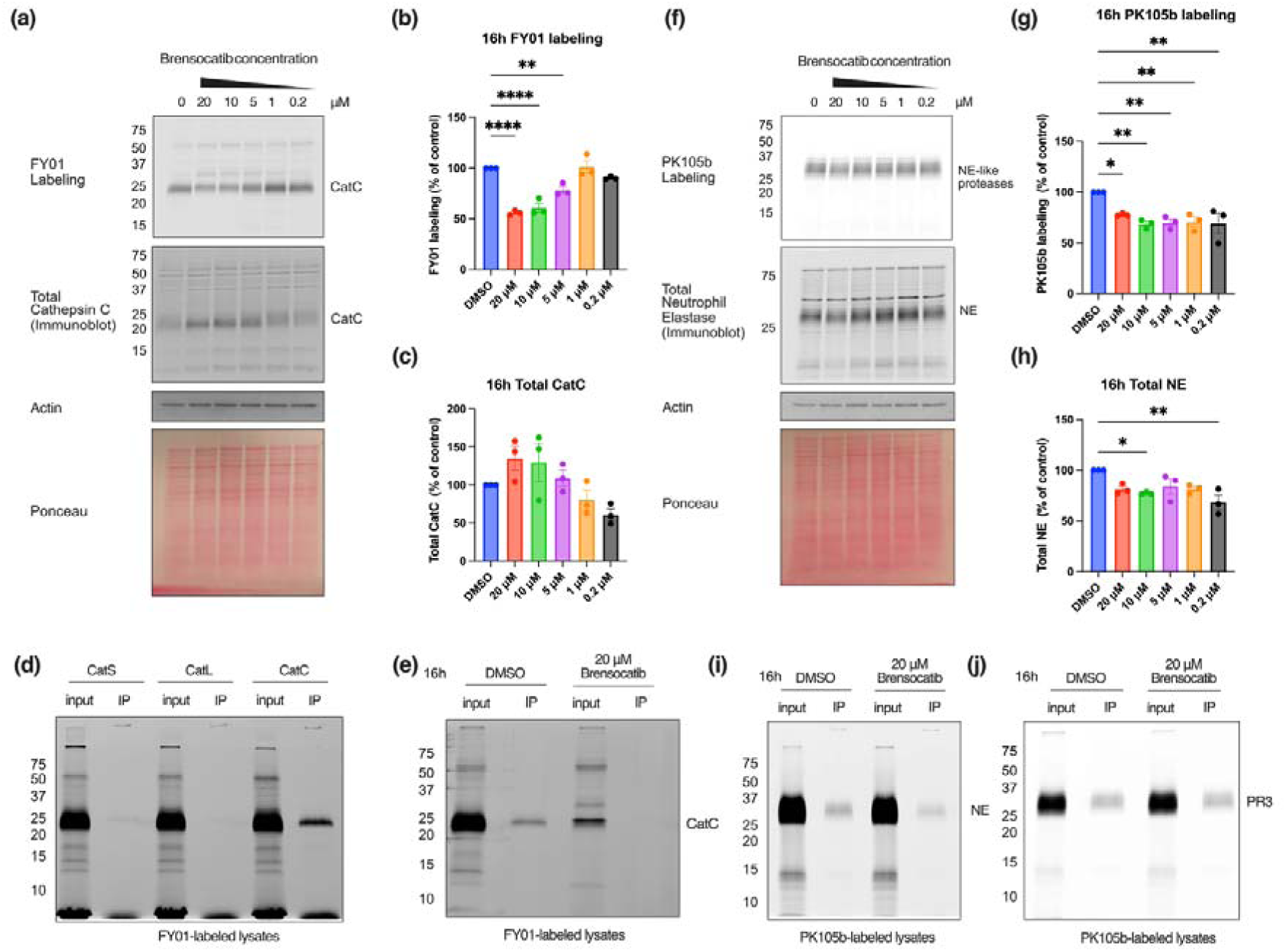
Concentration-dependent impact of 16-h brensocatib treatment on cathepsin C and elastase-like protease activity in HL-60 cells. HL-60 cells were treated with DMSO (vehicle) or brensocatib (0.2, 1, 5, 10, 20 μM), and cells were harvested after 16 h. **(a)** The FY01 activity-based probe was used to detect active CatC and is visualized by in-gel fluorescence, followed by immunoblotting for total CatC levels. Ponceau S stain and actin were used as loading and housekeeping controls, respectively. Densitometry of **(b)** FY01 labeling and **(c)** total CatC are graphed as a percentage of the DMSO control. **(d)** FY01 labeling of lysates from naïve cells before (input) and after immunoprecipitation (IP) antibodies for CatS, CatL, or CatC. **(e)** FY01 labeling of lysates before (input) and after immunoprecipitation (IP) with a CatC antibody before and after brensocatib treatment. **(f)** The PK105b activity-based probe was used to detect elastase-like activity and visualized by in-gel fluorescence, followed by immunoblotting to measure total NE levels. Densitometry of **(g)** active elastase-like proteases and **(h)** total NE are graphed as a percentage of the DMSO control. PK105b labeling of lysates before (input) and after IP with antibodies for **(i)** NE or **(j)** PR3 is shown for each condition. Data are represented as the mean ± SEM from three separate experiments (n=3), and ordinary one-way ANOVA with Dunnett’s multiple comparisons test comparing the mean with each column with the DMSO group mean was used for statistical analysis. \**p* < 0.05, \*\**p* < 0.01, \*\*\*\**p* < 0.0001.

In addition to measuring CatC activity, we also used an activity-based probe to measure the downstream impact on NSP activation after brensocatib treatment. PK105b is a fluorescent diphenyl phosphonate-based probe targeting elastase-like proteases including NE and PR3 (38, 39). At the 16-h timepoint, PK105b labeling was significantly suppressed (∼25%) across all brensocatib concentrations (**Fig 1f,g**), while the levels of total NE and PR3, as shown by immunoblot, did not change (**Fig. 1h, Supplementary Fig. 2a**). Immunoprecipitation of PK105b-labeled lysates with antibodies for NE and PR3 revealed that both proteases were labeled by this probe under these conditions, and that both were still active after 16-h brensocatib treatment (**Fig. 1i,j**).

Given the minimal impact of brensocatib on NSP activation observed after 16 h, we reasoned that sustained CatC inhibition may be required for more complete suppression. We therefore treated HL-60 cells with brensocatib for 72 h prior to assessment of CatC and NSP activity. In agreement with the 16-h treatment, CatC activity, as measured by FY01 labeling, was suppressed after treatment with 10-20 µM brensocatib for 72 h (p<0.001 and p<0.01, respectively; **Fig. 2a,b**). Immunoprecipitation with a CatC antibody revealed that CatC labeling was abolished by 20 µM brensocatib (**Fig. 2d**). Total CatC levels were slightly reduced at lower concentrations of 0.2 μM and 1 μM (**Fig. 2a,c**). At higher concentrations, the total amount was unchanged but a qualitative change in the CatC species was observed, possibly due to alterations in its processing as the result of perturbations in the proteolytic network (**Fig. 2a-c**). In contrast to the 16-h timepoint, PK105b labeling indicated that NSP activation was reduced by ∼70% at 20 μM upon 72-h treatment (**Fig. 2e,f**). Total NE protein abundance was minimally reduced at the lower drug concentrations (0.2 and 1 μM) but were unchanged at the higher doses (**Fig. 2e, g**). A qualitative shift in the total NE band was observed, with its accumulation in a higher molecular weight form observed at the higher concentrations of brensocatib (**Fig. 2e**). This likely represents immature NE that had not been cleaved by CatC. Immunoprecipitation of PK105b-labeled lysates revealed that both NE and PR3 activity were suppressed after 72 h of brensocatib treatment (**Fig 2h,i**), and total PR3 levels were reduced in the presence of brensocatib (**Supplementary Fig. 2b**). Collectively, these experiments reveal that CatC activity is rapidly inhibited by brensocatib (e.g., 16 h), but that longer incubation is required (e.g., 72 h) to observe significant suppression of CatC-mediated NSP activation.

**Figure 2.**
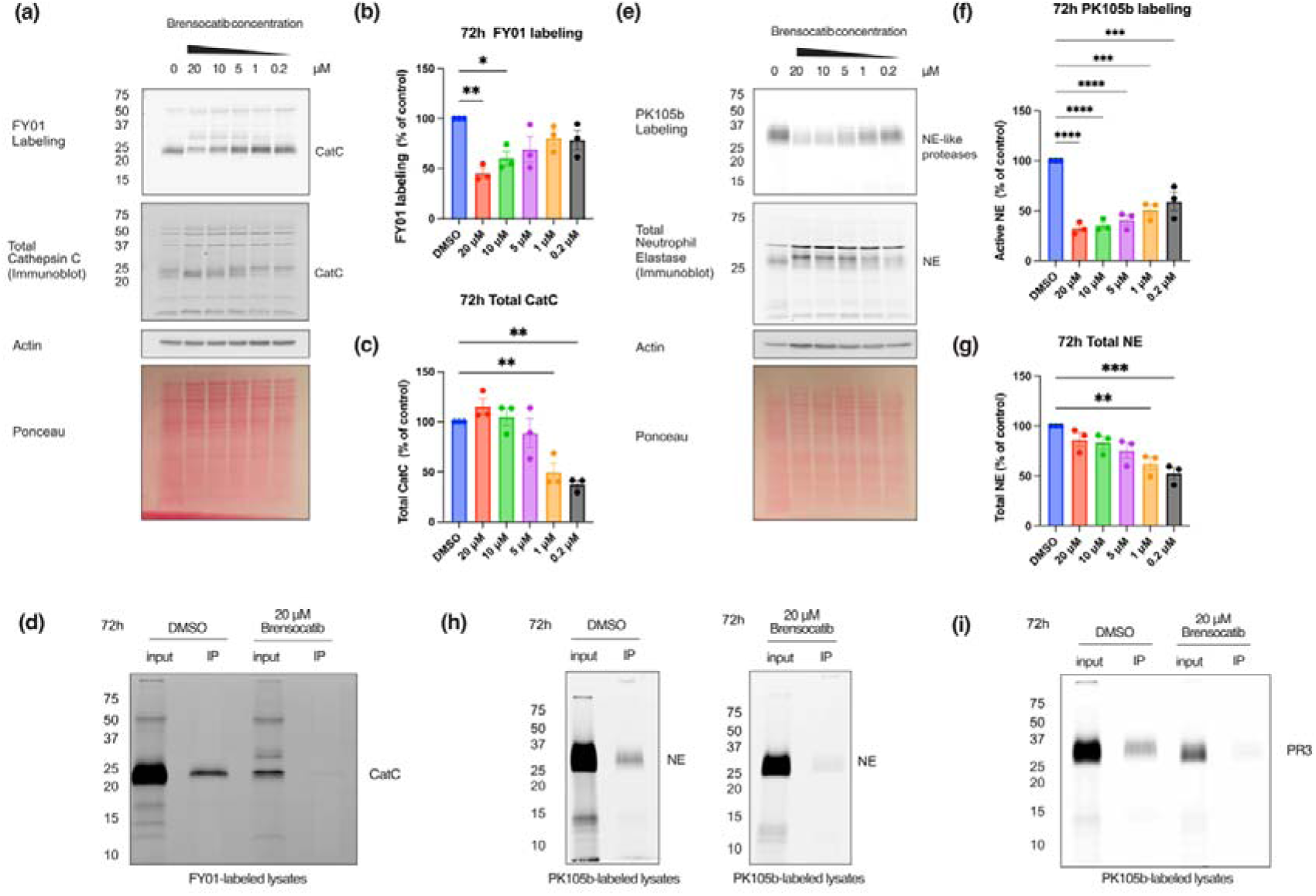
Concentration-dependent impact of 72-h brensocatib treatment on inhibition of cathepsin C and elastase-like protease activity in HL-60 cells. HL-60 cells were treated with DMSO (vehicle control) or brensocatib (0.2, 1, 5, 10, 20 μM), and cells were harvested after 72 h. **(a)** The FY01 activity-based probe was used to detect active CatC and visualized by in-gel fluorescence, followed by immunoblotting to measure total CatC levels. Ponceau S stain and actin were used as loading and housekeeping controls, respectively. Densitometry of **(b)** FY01 labeling and **(c)** total CatC are graphed as a percentage of the DMSO control. **(d)** FY01 labeling of lysates before (input) and immunoprecipitation (IP) with CatC antibody is shown for each condition. **(e)** The PK105b activity-based probe was used to detect elastase-like activity and visualized by in-gel fluorescence, followed by immunoblotting to measure total NE levels. Densitometry of **(e)** PK105b labeling and **(f)** total NE are graphed as a percentage of the DMSO control. PK105b labeling of lysates before (input) and after IP with antibodies for **(h)** NE or **(i)** PR3 is shown for each condition. Samples from the DMSO- and brensocatib-treated conditions for the NE pulldown were resolved on separate gels and scanned at same intensity. Data are represented as the mean ± SEM from three separate experiments (n=3), and ordinary one-way ANOVA with Dunnett’s multiple comparisons test comparing the mean with each column with the DMSO group mean was used for statistical analysis. \**p* < 0.05, \*\**p* < 0.01, \*\*\**p* < 0.001, \*\*\*\**p* < 0.0001.

Based on the initial concentration-response studies, 20 μM brensocatib was selected as the concentration that achieved maximal inhibition of both CatC and the downstream NSPs, enabling assessment of time-dependent consequences of CatC inhibition. We further tested this concentration at three timepoints in a side-by-side comparison: 16 h, 72 h and 7 days. CatC activity was inhibited by brensocatib at all timepoints relative to vehicle control, with similar magnitude of inhibition across the time course (FY01 labeling reduced by 45%, 41% and 30% at 16 h, 72 h, 7 days respectively) (**Fig. 3a,b**). While total CatC levels did not differ significantly between control and treatment groups, the qualitative change in CatC species was again apparent across the time course (**Fig. 3a,c**). PK105b labeling revealed a pronounced, time-dependent decrease in NSP activity, from 84% residual activity at 16-h to 35% and 11% at 72 h and 7 days, respectively (**Fig. 3d,e**). This reflects cumulative loss of NSP activity following sustained inhibition of their activation by brensocatib, consistent with previous studies (2, 36). Total levels of NE were not significantly different to the control across the time course (**Fig. 3d,f**), while PR3 levels were significantly reduced at 72 h and 7 days (**Supplementary Fig. 2c**). This is consistent with previous studies reporting reduced PR3 levels following both direct and indirect CatC inhibition and may suggest differential regulation of individual NSPs following CatC inhibition (5, 36).

**Figure 3.**
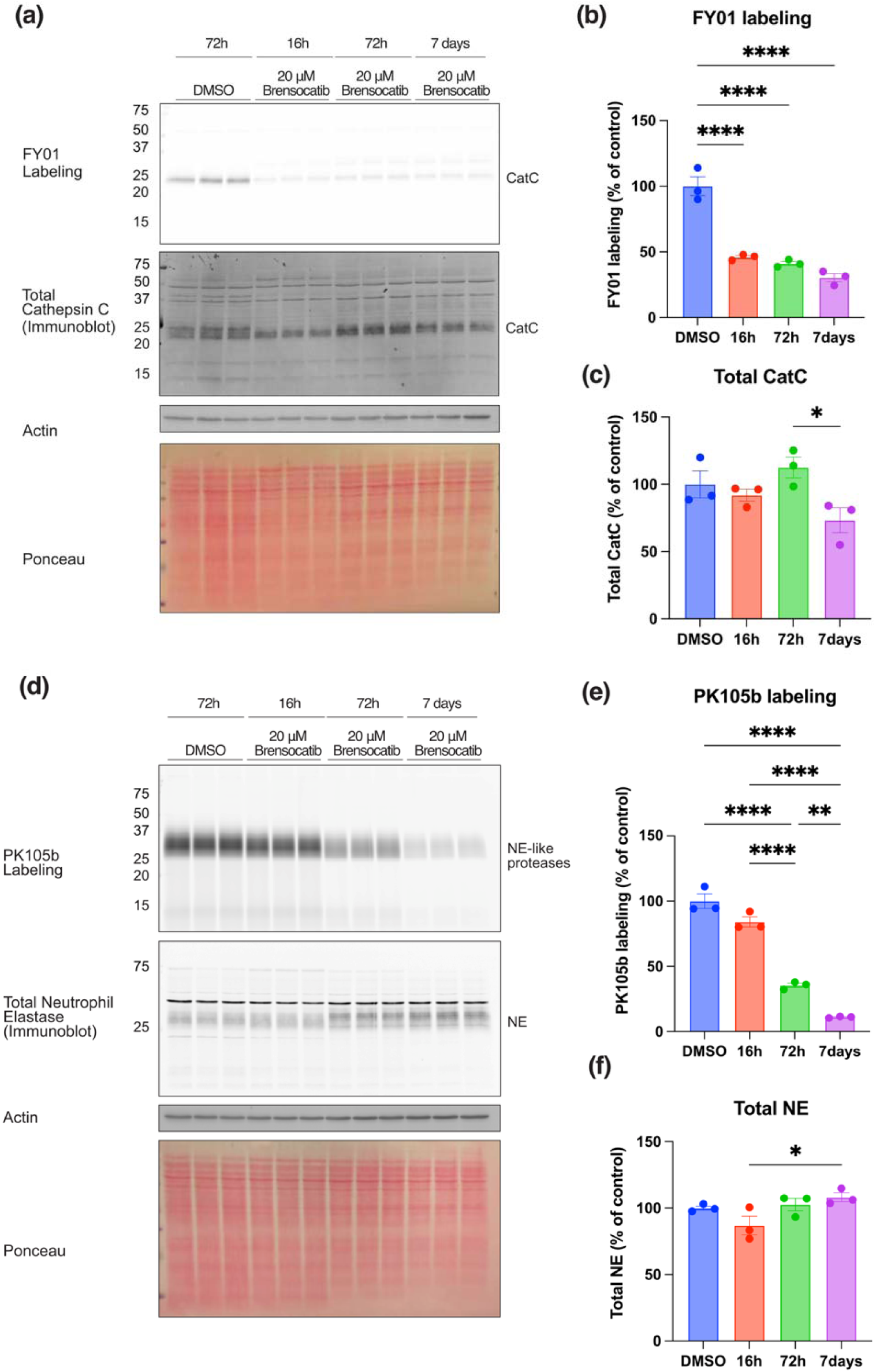
Time-dependent inhibition of CatC and NE by brensocatib in HL-60 cells. HL-60 cells were treated with DMSO (vehicle control) or brensocatib (20 μM), and cells were harvested after 16 h, 72 h, or 7 days (n=3/group). **(a)** The FY01 activity-based probe was used to detect active CatC and visualized by in-gel fluorescence, followed by immunoblotting to measure total CatC levels. Ponceau S stain and actin were used as loading and housekeeping controls, respectively. Densitometry of (b) FY01 labeling and (c) total CatC are graphed relative to the DMSO control following actin normalization. **(d)** The PK105b activity-based probe was used to detect elastase-like activity and visualized by in-gel fluorescence, followed by immunoblotting to measure total NE levels. Densitometry of **(e)** PK105b labeling and **(f)** total NE are graphed relative to the DMSO control. Data are represented as the mean ± SEM, and ordinary one-way ANOVA with Tukey’s multiple comparisons test. \****p*** < 0.05, \*\****p*** < 0.01, \*\*\*\****p*** < 0.0001.

Together, these findings highlight that continuous inhibition of CatC is required to observe suppression of NSP activation, likely owing to the requirement for previously activated NSPs to be turned over. As additional alternative, yet unidentified, NSP-activating proteases, have been proposed, along with the reversible nature of brensocatib, full suppression of NSP activation is unlikely to be achieved with this treatment alone (36, 40, 41); thus, we proceeded to investigate the impact of CatC-mediated activity on the proteome and degradome within these parameters.

### Longitudinal impact of brensocatib treatment on the global HL-60 proteome

To determine the impact of CatC inhibition on the HL-60 proteome, we applied No-Enrichment Identification of Cleavage Events (NICE), a dimethylation-based N-terminomics pipeline that enables simultaneous quantification of protein abundance and proteolytic cleavages events without the need for enrichment steps (**Supplementary Fig. 1c**) (29, 30, 42, 43). HL-60 cells were treated with 20 µM brensocatib or DMSO vehicle for 16 h, 72 h, or 7 days prior to harvesting for NICE analysis. Native and neo-N-termini of denatured proteins, along with lysine side chains, were dimethylated prior to trypsin digestion. To achieve comprehensive proteome coverage and quantitative reproducibility, data-independent acquisition (DIA) mass spectrometry was employed. Internal peptides with a free amine (generated by trypsin digest) and peptides with N-terminal dimethylation (native and neo-N-termini) were analyzed together to quantify protein abundance, while the latter was filtered bioinformatically to quantify cleavage events. Principal component analysis (PCA) indicated clear clustering of the DMSO- and brensocatib-treated samples at both protein and peptide levels, with progressively improved separation observed over time, particularly at the peptide level (**Supplementary Fig. 3**).

To investigate the impact of cathepsin C inhibition on protein abundance over time, all proteins detected in at least three of the four replicates in at least one group were included for analysis (**Supplementary Table 1**). Of 5,467 proteins detected at the 16-h timepoint, only seven were altered by >2-fold with p<0.05, including two enriched in brensocatib-treated cells (ANXA8 and UBQLN2) and five enriched in DMSO-treated cells (MYC, CARNMT1, POC1A, STMN4, GINS2) (**Fig. 4a**). Of these 7, only MYC was significant following FDR correction (q<0.05). MYC is an oncogene involved in the cell cycle and expressed in HL-60 cells, but to a lesser extent when they are differentiated into mature granulocytes (45). The observed downregulation of MYC after brensocatib treatment (q value < 0.05) may reflect an early shift towards granulocytic differentiation or reduced proliferative signalling.

**Figure 4.**
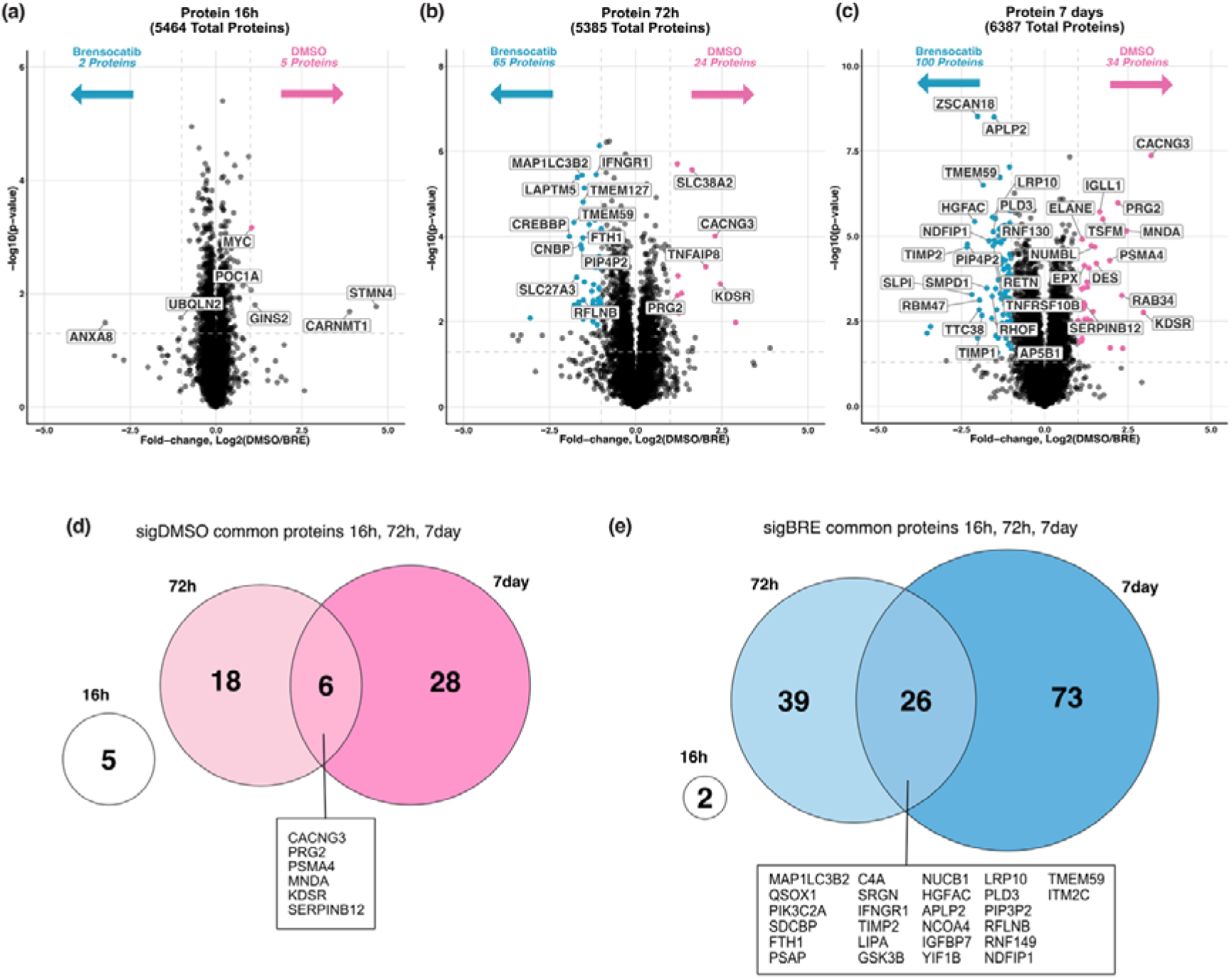
Time-dependent impact of brensocatib (20μM) treatment on the global proteome of HL-60 cells. HL-60 cell lysates were subject to NICE analysis. Proteins identified in ≥3 of 4 biological replicates in at least one group (n = 4/group) were analyzed by Student’s t-test and visualized by volcano plot. A threshold of –log_10_(*p*-value) ≥ 1.3 (corresponding to *p* < 0.05; horizontal lines) and |log_2_(DMSO/Bre)| > 1 (vertical dotted lines) was applied. DSMO- and brensocatib-enriched proteins that passed q value < 0.05 are highlighted in pink and blue, respectively at **(a)** 16 h, **(b)** 72 h, and **(c)** 7 days. Venn diagram of overlapping proteins enriched in **(d)** DMSO- and **(e)** brensocatib-treated cells at 16 h, 72 h, and 7 days. \****p*** ≤ 0.05, \*\****p*** ≤ 0.01, \*\*\****p*** ≤ 0.001, \*\*\*\****p*** ≤ 0.0001.

As expected, longer incubation times resulted in increased protein level changes. Of 5,388 proteins detected at the 72-h timepoint, 65 proteins were enriched in the brensocatib-treated cells, while 24 proteins were enriched in the DMSO-control group (fold change >2; p<0.05; **Fig. 4b**). After 7 days of drug treatment, 6,393 proteins were identified, with 100 proteins enriched following brensocatib treatment and 34 enriched following DMSO treatment (**Fig. 4c**).

Comparison of protein level changes across the three timepoints revealed that changes at the 16-h timepoint were unique, and 32 proteins were consistently altered between 72-h and 7-day treatments (**Fig. 4d,e**). Six proteins were commonly increased in the DMSO group (**Fig. 4d, Supplementary Table 2)**, including proteoglycan 2 (PRG2) and serine protease inhibitor SERPINB12 (44). By contrast, 26 proteins were commonly enriched in brensocatib-treated cells after 72 h and 7 days, including proteins involved in immune responses, granule secretory pathway regulation, cellular signaling, and homeostasis, suggesting long-term cellular adaptations (**Fig. 4e, Supplementary Table 3)**. Increased abundance of serglycin (SRGN) and ubiquitination-mediated turnover proteins (E3 ubiquitin-protein ligase RNF149, Nedd4 family-interacting protein 1 NDFIP1) were found in both groups which may reflect cellular adaptation to CatC inhibition (45–47).

Importantly, we observed reduced abundance of NSPs in brensocatib-treated cells across the two later timepoints (log_2_(DMSO/Bre)> 0.5) (**Supplementary Table 1**). Particularly at the 7-day timepoint, NE (log_2_(DMSO/Bre)=1.1; q=0.001), PR3 (log_2_(DMSO/Bre)=0.6; q=0.01), and CTSG (log_2_(DMSO/Bre)=0.5; q=0.01) were significantly downregulated after brensocatib-treatment, consistent with the impaired maturation by CatC and potentially reflecting degradation of their zymogens at the 7-day timepoint (**Fig. 3d**).

Gene Ontology (GO) analysis of the brensocatib-enriched proteins revealed an enrichment in biological processes associated with negative regulation of proteolysis (**Fig. 5b,d**). Of particular note, we observed increased abundance of other cathepsin proteases, including CatD (log_2_(DMSO/Bre)=-0.8; q=0.0005), CTSB (log_2_(DMSO/Bre)=-0.8; q=0.001), CatZ (log_2_(DMSO/Bre)=-1.0; q=0.0006), CTSA (log_2_(DMSO/Bre)=-0.5; q=0.0006) and CatS (log_2_(DMSO/Bre)=-0.4; q=0.004) (**Supplementary Table 1**). We confirmed increased expression of CatB and CatX by immunoblot (**Supplementary Fig. 4**). In addition, a number of protease inhibitors were upregulated following CatC inhibition, including kunitz-type serine peptidase inhibitor 1 (SPINT1, log_2_(DMSO/Bre)=-1.4; q=0.001), tissue inhibitor of metalloproteinases 1 (TIMP1, log_2_(DMSO/Bre)=-1.8; q=0.0008), tissue inhibitor of metalloproteinases 2 (TIMP2, log_2_(DMSO/Bre)=-1.9; q=0.006), secretory leukocyte protease inhibitor (SLPI, log_2_(DMSO/Bre)= −2.2; q=0.002), and calpastatin (CAST, log_2_(DMSO/Bre)=-1.3; q=0.0006) (**Supplementary Table 1**). In agreement with our observations, SLPI expression was elevated in bronchiectasis patients receiving brensocatib and shows an inverse correlation with NE abundance (48, 49). As NE cleaves and inactivates SLPI, this may reflect reduced SLPI degradation (50, 51). Collectively, these findings demonstrate widespread remodeling of the protease-anti-protease network following CatC inhibition. We next investigated how these changes influenced protein processing across the whole degradome.

**Figure 5.**
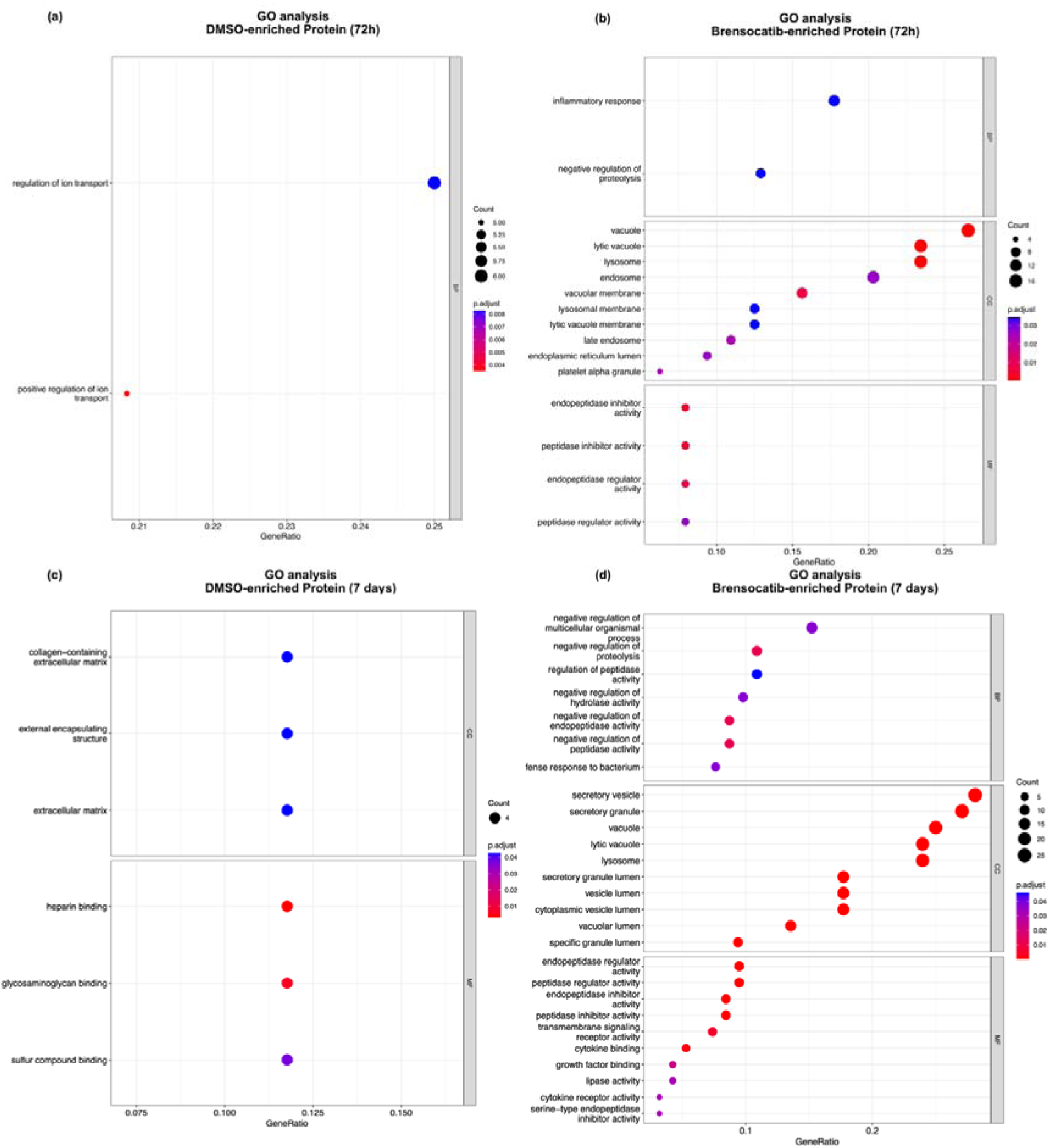
Protein-level analysis of HL-60 cells treated with DMSO or brensocatib (20 μM) for 72 h or 7 days (n=4/group). Gene ontology biological processes enrichment dot plot of differentially expressed proteins detected in the DMSO-treated group and brensocatib-treated cells after 72 h **(a-b)**, and 7 days **(c-d)** respectively. All proteins identified in the corresponding datasets were used as the background universe for each enrichment analysis.

### Longitudinal impact of brensocatib treatment on proteolytic cleavage events

To identify proteolytic cleavage events that were altered upon CatC inhibition with brensocatib, either directly or through altered downstream protease networks, we filtered the NICE datasets for peptides with high-confidence N-terminal dimethylation. Sample dimethylation efficiency was assessed by examining the amino acid distribution at residues preceding the cleavage sites and at the end of the N-terminal peptides. Consistent with the parameters of a semi-tryptic search, most of the peptides ended in arginine (∼98%; **Supplementary Fig. 5a,c,e)**. Residues preceding the identified peptide N-termini, corresponding to the P1 position of the cleavage sites, displayed a broader amino acid distribution, indicating cleavages at non-tryptic sites **(Supplementary Fig. 5b,d,f)**. We also examined the proportion of N-terminal peptides bearing fully or partially dimethylated lysines (**Supplementary Fig. 6a,c,e**). As the assignment of localization in DIA datasets requires additional scrutiny (52, 53), only peptides containing fully dimethylated internal lysine residues were considered for analysis to avoid potential false positives. Only peptides identified in at least three of the four samples in at least one of the groups were included. Following this filtering, 2,661, 2,868, and 2,444 peptides in the three timepoints, respectively, were classified as dimethylated native or neo-N-termini (**Supplementary Fig. 6b,d,f**).

Of the 2,661 N-termini detected in the 16-h samples, 33 and 47 peptides were enriched in the brensocatib- and DMSO-treated groups, respectively (**Fig. 6a**). After 72 h and 7 days, the number of DMSO-enriched neo-N-termini increased substantially (1008 and 1156 among 2,868 and 2,444 total N-termini, respectively), as did the brensocatib-enriched cleavage sites (110 and 109, respectively) (**Fig. 6b,c**).

**Figure 6.**
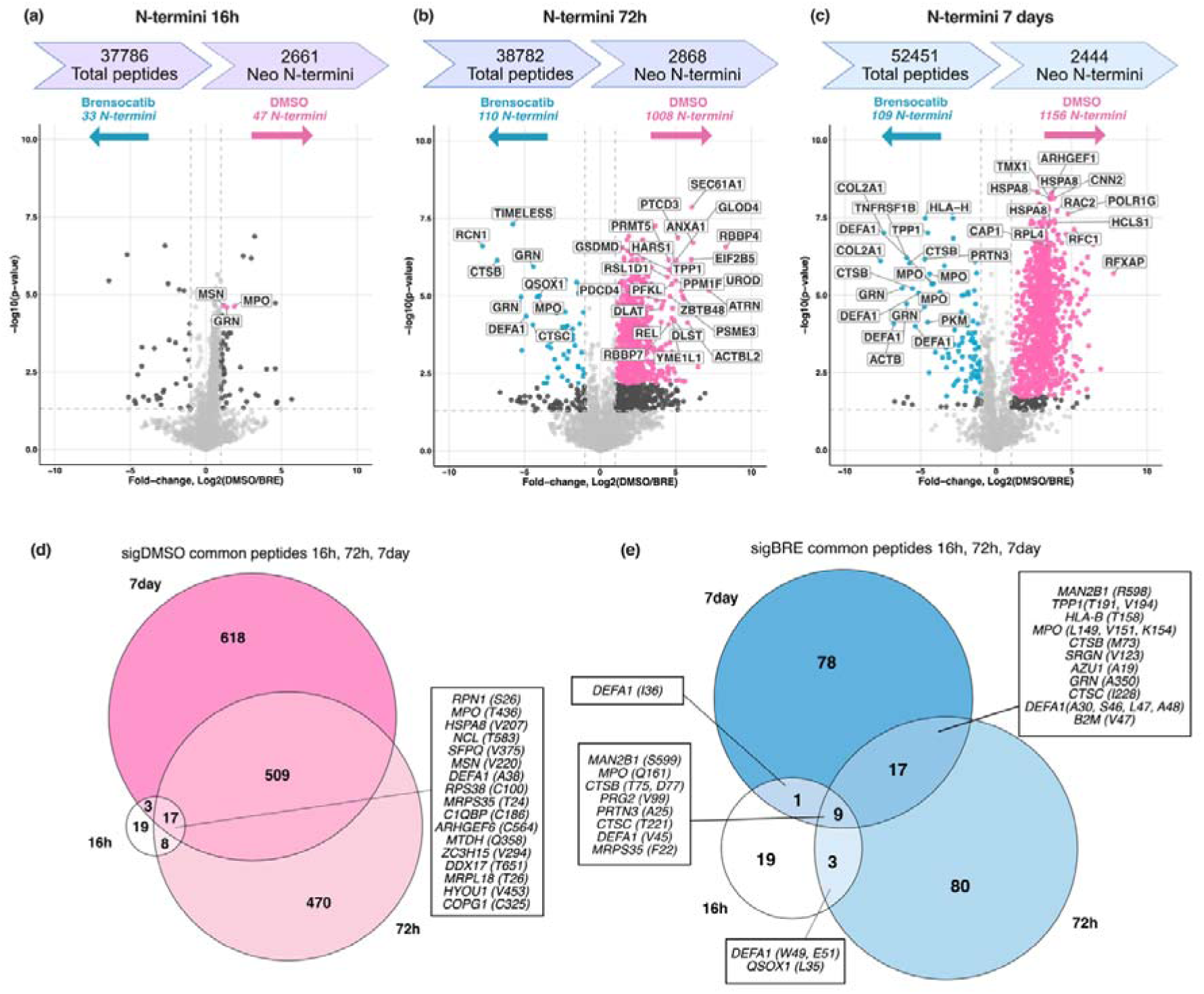
Time-dependent impact of brensocatib (20 μM) treatment on the degradome of HL-60 cells. HL-60 cell lysates were subject to NICE analysis. Native and protease-generated peptides (neo-N-termini) that were detected in at least three of four replicates in at least one group (n=4/group) were plotted using Student’s t-test with a threshold of –log_10_(p-value) ≥ 1.3 (corresponding to p < 0.05; horizontal dotted lines) and |log_2_(DMSO/Bre)| > 1 (vertical dotted lines). N-termini meeting these criteria but not passing multiple testing correction are shown in dark grey. N-termini significantly enriched after multiple testing correction (FDR q < 0.05) are highlighted in pink (DMSO-enriched) or blue (brensocatib-enriched) at **(a)** 16 h, **(b)** 72 h, and **(c)** 7 days. Number of total peptides and total N-termini are shown above each volcano plot, along with number of significantly enriched N-termini in each group. **(d-e)** Venn diagrams showing overlapping N-termini and the corresponding P1 site that were enriched at each timepoint in DMSO- and brensocatib-treated cells, respectively.

The DMSO-enriched N-termini represent cleavage events mediated by CatC-dependent proteolytic networks, including the proteases that it regulates (e.g., NSPs). Among these, we observed substantial overlap across the 3 datasets, including 17 N-termini detected at all three timepoints and 509 detected at both 72 h and 7 days (**Fig. 6d**). We also observed cleavage events that were uniquely detected at each timepoint (19, 470, 618 unique cleavage events at 16 h, 72 h, and 7 days, respectively). At 72-h and 7-day timepoints, GO analysis suggested DMSO-enriched N-termini were associated with metabolic processes **(Supplementary Fig. 7a,c)**, whereas brensocatib-enriched cleavage events mapped to proteins involved in cell death pathways, including cathepsins (CatB, CatD, CatC) and granular proteins (MPO, ELANE, SRGN, AZU1) **(Supplementary Fig. 7b,d)**.

### Time-dependent emergence of a neutrophil serine protease cleavage signature

To identify motifs among the cleavage sites altered by brensocatib, we generated pLogos of enriched sites (P5-P5’) within each group at each timepoint, relative to a background of P5-P5’ residues of all detected N-termini. At 16 h, no clear motifs were observed (**Fig. 7a**). After 72 h and 7 days, however, the DMSO-enriched sites exhibited a significant enrichment of valine (Val), threonine (Thr), cysteine (Cys), isoleucine (Ile), and alanine (Ala) residues at the P1 position (ordered based on significance) (**Fig. 7b, c**). While the brensocatib-enriched cleavage sites did not contain significant motifs at any timepoint, indicating a more heterogeneous proteolytic landscape (**Fig. 7d-f**). This was further evident in the volcano plots depicting time-dependent shifts in the abundance of cleavage events at the P1 position (**Supplementary Fig. 9**).

**Figure 7.**
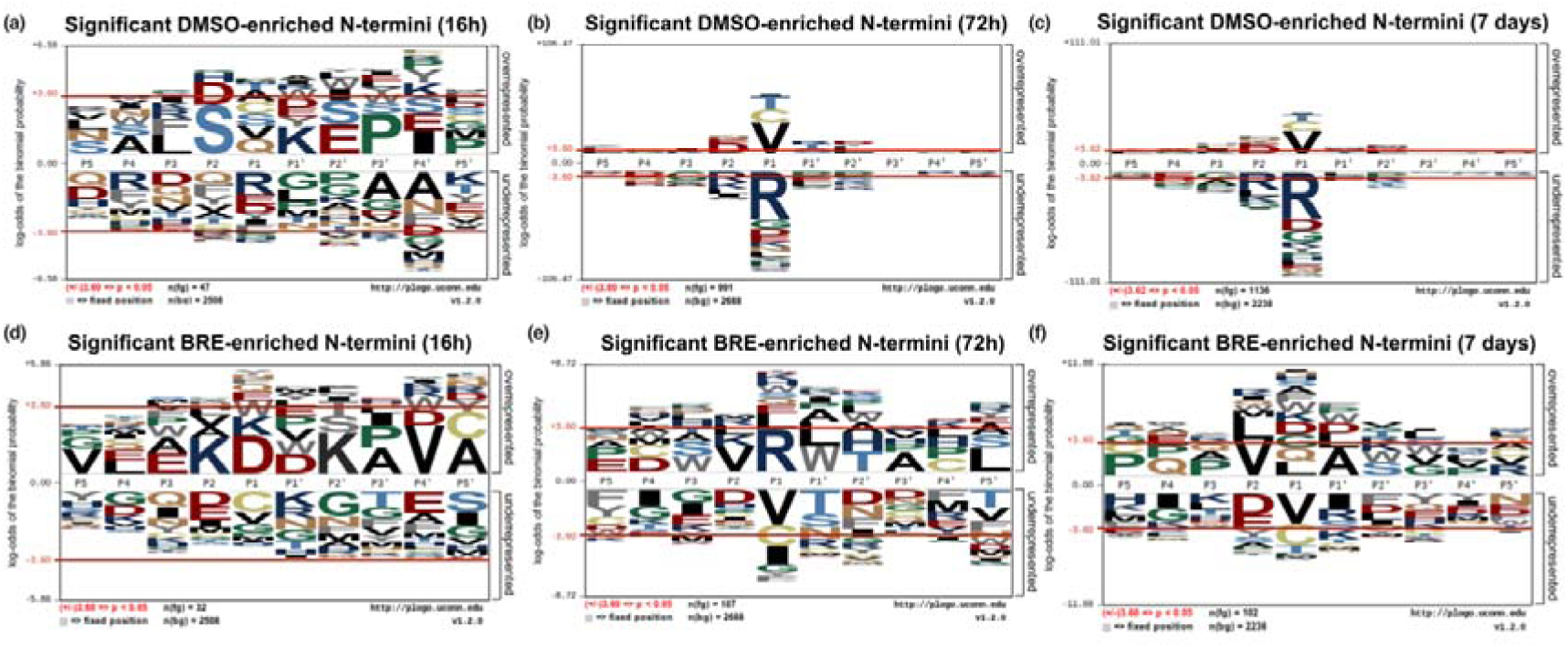
Consensus motifs of cleavage sites enriched over time in DMSO- or brensocatib-treated HL-60 cells. P5-P5’ residues for each cleavage site enriched in DMSO- or brensocatib-treated cells at **(a,d)** 16 h, **(b,e)** 72 h and **(c,f)** 7 days were subject to pLogo analysis against the background of all cleavage events identified within each dataset. Over-represented and underrepresented amino acids are determined by the cutoff of p<0.05 (log-odds of the binomial distribution ±3; red lines).

The time-dependent shift in the PLogo motif supports our previous activity measurements (**Fig. 3**), where residual NSP activity was 84%, 35% and 11% at 16 h, 72 h and 7 days, respectively. The enriched P1 residues may reflect the P1 preferences of NE and PR3 for small aliphatic residues (Val, Ala, Ile), and their tolerance for Thr (54–56). P1 Cys is likely compatible with the S1 binding pocket of NE and PR3. Although it is not typically reported, likely due to the omission of Cys from positional scanning libraries, it has been highlighted in elastase-related cleavages events in human cancer-specific proteolytic signatures (54). P2 aspartic acid (Asp) and glutamic acid (Glu) preferences potentially reflect PR3 activity, as it is known to prefer negatively charged amino acid at this position (57). CatG is known to prefer aromatic or aliphatic P1 residues phenylalanine (Phe), tyrosine (Tyr), tryptophan (Trp), and leucine (Leu) (58). While there was some evidence of these sites enriched in the 7-day DMSO treatment (Leu 11; Phe 10; Tyr 2), this was not the dominant signature (**Supplementary Table 1**). In brensocatib-treated cells, this dominant NE/PR3 signature was collapsed both in proportion and absolute number, with increased representation of basic and acidic residues (**Fig. 7, Supplementary Fig. 8**). These observations collectively support that prolonged CatC inhibition gradually leads to inhibition of NSP activation, which in turn limits cleavage of their substrates.

### Dipeptidyl peptidase signatures

CatC primarily functions as a dipeptidyl aminopeptidase that removes two amino acids from the N-terminus of proteins. To identify candidate CatC substrates, we searched our NICE dataset for cleavage sites separated by two residues between treatment groups. This identified 10 candidate proteins (**Table 1, Supplementary Table 4**), including the established CatC substrate PR3 (59). Consistent with known CatC-dependent maturation, PR3 was predominantly cleaved at E27↓I28 in DMSO-treated cells, while upstream cleavage at A25↓A26 accumulated in brensocatib-treated cells (7-days) (**Table 1; Supplementary Fig. 10a**), reflecting retention of the two-residue propeptide. Although NE is a known substrate of CatC, we did not observe altered processing of the propeptide S28-E29 in our analysis. We did, however, observe increased cleavage at G23↓T24 after the 7-day brensocatib treatment, which may correlate to the increased higher molecular weight NE species observed by immunoblot (**Supplementary Fig. 10b, Fig. 3d**). In the DMSO control group, we also consistently identified NE peptides arising from cleavage at A131↓N132 across the 72-h and 7-day timepoints. Enrichment of NE peptides cleaved at N130↓A131 and N132↓V133 was previously observed in sputum of chronic obstructive pulmonary disease patients with exacerbation compared to those with clinically stable conditions (60). Given cleavage within this region would disrupt the active site of NE by removing the catalytic residue S202, it is not expected to promote its activity but may instead reflect altered turnover of NE in response to drug treatment.

**Table 1.**
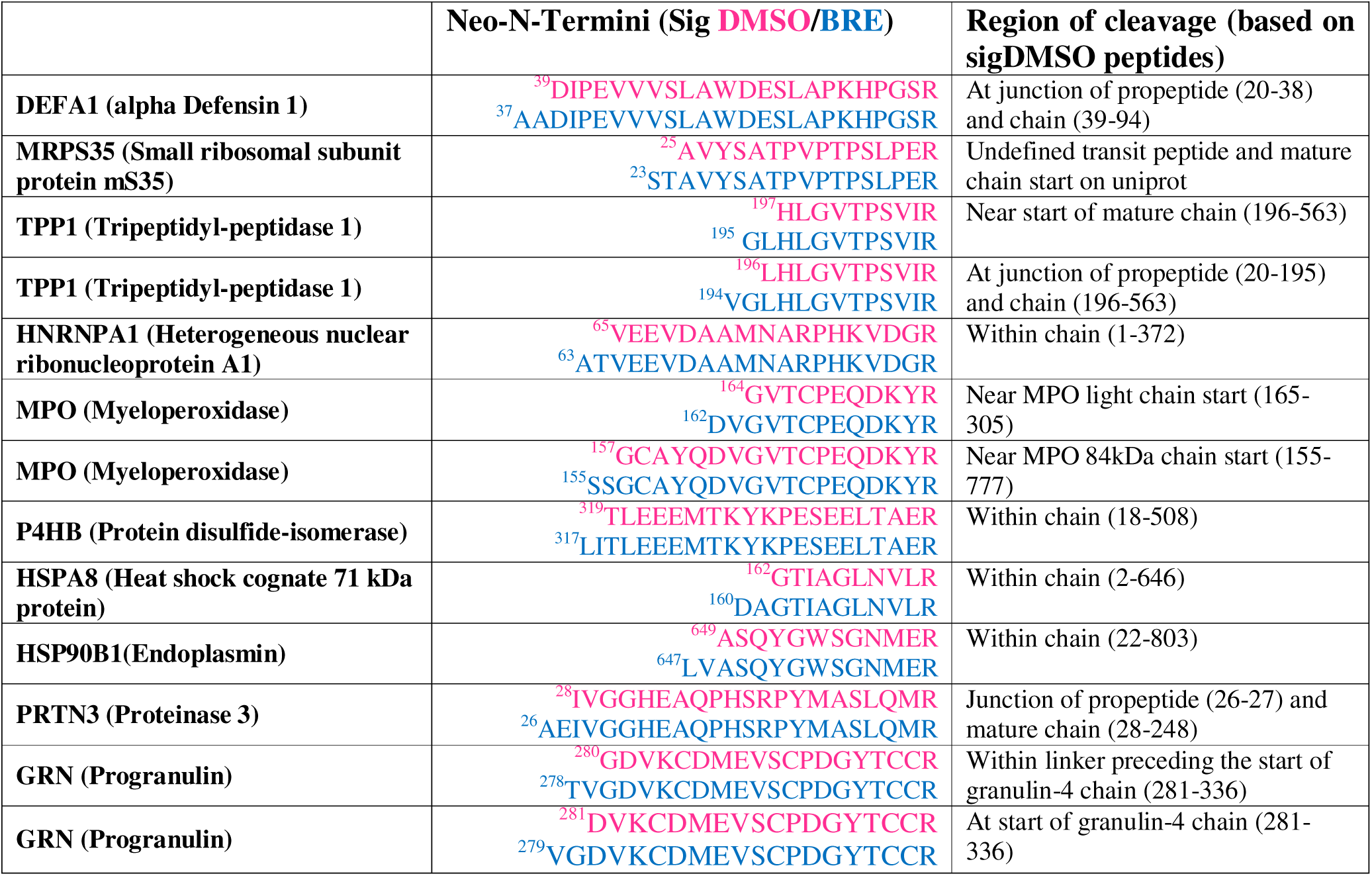
Summary of potential cathepsin C mediated cleavage.

We identified a further 9 proteins exhibiting a 2-residue shift that may represent potential CatC substrates (**Table 1**). Tripeptidyl peptidase 1 (TPP-1) was differentially cleaved at T193↓V194 and G195↓L196 in brensocatib- and DMSO-treated cells, respectively (72 h). The two-residue shift within the linker between the propeptide and mature enzyme is consistent with suppressed N-terminal dipeptide cleavage following CatC inhibition (**Supplementary Fig. 10c**). TPP1 is a serine protease localized within lysosomes that functions both as an exopeptidase at pH 5, as well as an endopeptidase under more acidic conditions (61). TPP1 has been reported to autoactivate *in vitro* through cleavage at L196 at pH 3.5, or at S181↓L182, Q189↓V190, D168↓F169 in higher pH conditions (61, 62). *In vivo*, its activation is proposed to occur within lysosomes and be mediated by a serine protease sensitive to AEBSF inhibition (63). This prior observation may suggest that TPP1 cleavage is mediated by CatC-dependent NSPs rather than by CatC directly, although additional studies are required to delineate these hypotheses in HL-60 cells.

Progranulin (GRN) is a glycoprotein that undergoes proteolytic processing in lysosomes and extracellular environments to generate granulin peptides. We detected differential cleavage within the linker preceding granulin-4 (GRN-4) (**Supplementary Fig. 10d**). Cleavage at H277↓T278 and T278↓V279 was enriched in brensocatib-treated cells, while cleavage at V279↓G280 and G280↓D281 was enriched in DMSO-treated cells. These sites differ by two residues, a pattern consistent with reduced N-terminal dipeptide trimming following CatC inhibition. Although CatC is unable to digest full-length GRN (64), it may contribute to secondary processing following endopeptidase events by other proteases (e.g., cathepsin L, PR3 (65), matrix metalloproteinases (66), elastase (67, 68)).

Alpha defensin 1 (DEFA1) is an antimicrobial peptide stored within neutrophil azurophilic granules. Previous studies have identified processing steps of prodefensin in HL-60 cells, including removal of the signal peptide A19↓E20, cleavage at A38↓D39 that yields an intermediate form of defensin and at A65↓C66 or M64↓A65 to produce the smaller mature forms (**Supplementary Fig. 11a**) (69, 70). *In vitro,* NE and PR3 are able to release the smaller mature alpha defensin peptide from the prodefensin, while the protease responsible for generating the intermediate form has not been characterized definitively (71). We observed differential cleavage at I36↓A37 and 38A↓D39 in the brensocatib- and DMSO-treated cells, respectively. These observations may be consistent with proteolytic trimming of DEFA1 to its intermediate form for subsequent activation.

Collectively, these observations identify several candidate CatC-dependent processing events. Whether these proteins represent direct CatC substrates or are processed indirectly through altered NSP activity or secondary protease networks will require further biochemical validation.

### Proteolytic remodeling in the absence of CatC activity

Following CatC inhibition, a distinct set of peptides became enriched, where 9 N-termini were enriched in brensocatib-treated cells across all timepoints, 17 shared between 72 h and 7 days, and 19, 80, and 78 sites unique to 16 h, 72 h, and 7 days, respectively (**Fig. 6e**). These cleavage events likely reflect protease activities that predominate in the absence of both CatC and NSP activity.

Among the cleavage events enriched in brensocatib-treated cells across all timepoints, we observed CatB cleaved at T75↓E76 and D77↓L78, situated near the near the start of the mature chain (L80) (**Fig. 6e; Supplementary Fig 12a**) (72). This likely reflects the increase in total CatB levels following treatment (log_2_(DMSO/Bre)=-0.8; q=0.0009; **Supplementary Table 1**) (73). Brensocatib treatment also appeared to alter CatC cleavage patterns just proximal to the mature enzyme, but independent of protein abundance. At both 72 h and 7 days, cleavage of CatC at T221↓A222 and I228↓L229 were enriched in brensocatib-treated cells, while Q226↓K227, K227↓I228 and L229↓H230 were enriched in the DMSO group (**Supplementary Fig. 12b**).

At the two later timepoints, cleavage of azurocidin (AZU1) at A19↓G20, corresponding to the annotated mature chain, was enhanced after brensocatib treatment (log_2_(DMSO/Bre)=-2.8; q=0.004, 7-day timepoint) without associated changes in total protein levels (log_2_(DMSO/Bre)=-0.3; q=0.03) (**Fig. 6e, Supplementary Fig. 12c)**. As AZU1 is stored within azurophilic granules, this cleavage event may reflect altered maturation or processing of granule-associated immune proteins following CatC inhibition. AZU1 sputum concentration was significantly reduced following brensocatib treatment in the phase 2 WILLOW clinical trial, which may reflect differences in extracellular degranulation as opposed to the intracellular processing we observed in the HL-60 cells (48, 74, 75). Increased cleavage of the mitochondrial serine protease HTRA2 was detected at A133↓A134 (log_2_(DMSO/Bre)=-1.2, q=0.004), corresponding to the C-terminus of its propeptide (**Supplementary Fig. 12d**). As total HTRA2 protein abundance was unchanged (log_2_(DMSO/Bre) = 0.084, q = 0.045), altering its proteolytic processing may influence its role apoptotic signalling pathways (76).

As total CatZ and CatB levels were enriched in brensocatib-treated cells after 7 days (log_2_(DMSO/Bre)=-1.0; q=0.0006 and log_2_(DMSO/Bre)=-0.8; q=0.001, respectively; **Supplementary Fig. 4**), we investigated whether their activity could contribute to cleavage events enriched in the absence of CatC/NSP activity. Since these cysteine proteases are predominantly carboxypeptidases, N-terminomics analysis is insufficient to identify their substrates owing to the short N-termini generated (1-2 residues long). We therefore re-analyzed the 7-day dataset with SEMI search parameters to identify putative C-termini (non-tryptic peptides cut after residues other than arginine). Among 50 neo-C-termini enriched across 35 proteins in the brensocatib-treated cells, 8 events (16%) occurred in the last 10 residues (SRGN, AZU1, MAN2B1, PGAM1), potentially indicating carboxypeptidase activity (**Fig. 8a**, **Supplementary Table 5**). Among the 620 DMSO-enriched C-termini, only 5 (0.8%) occurred within the last 10 residues of the native N-termini. Across all C-termini, the P1 enrichment signature mirrored the N-termini (**Fig. 8b**), with 84 DMSO-enriched sites detected in both searches and 1162 and 580 uniquely detected as N-termini and C-termini (**Fig. 8c,d**).

**Figure 8.**
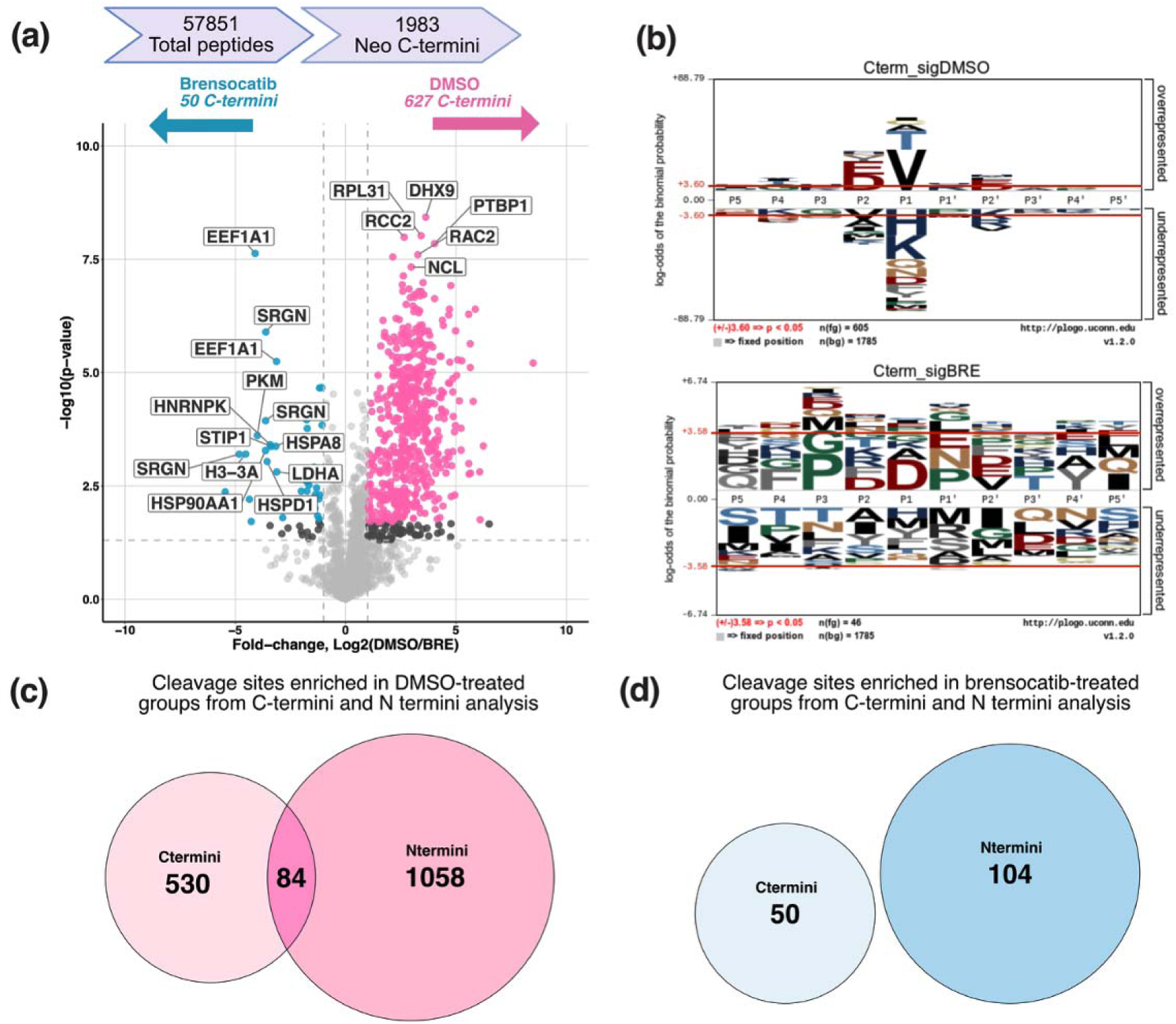
C-termini SEMI search of the 7-day brensocatib- and DMSO-treated samples data (n=4/group). **(a)** Volcano plot showing C-termini increased using a threshold of – log_10_(*p*-value) ≥ 1.3 (corresponding to *p* < 0.05; horizontal lines) and |log_2_(DMSO/Bre)| > 1 (vertical lines). **(b)** P5-P5’ residues for cleavage sites enriched in DMSO-treated or brensocatib-treated cells. Number of overlapping cleavage sites identified in both C-termini and N-termini analysis enriched in the **(c)** DMSO-treated group and **(d)** Bre-treated group.

## Conclusion

CatC is widely recognized for its role in activating NSPs, yet understanding of the broader impact of its inhibition on the proteome and degradome has remained incomplete. By integrating activity-based probes and an enrichment-free degradomics workflow, we demonstrate that pharmacological inhibition of CatC with brensocatib progressively suppresses NSP activation, resulting in remodeling of the proteolytic landscape. These analyses revealed a characteristic collapse of NSP cleavage signature and identified candidate CatC-dependent processing events and proteolytic adaptations that emerge in the absence of CatC activity.

Collectively, these findings extend the current understanding of CatC biology by providing a global view of how sustained CatC inhibition reshapes intracellular proteolytic networks. Beyond this, we reinforce the utility of unbiased, enrichment-free degradomics for characterizing protease function in living cells. The resulting resource of candidate substrates and cleavage events provides a foundation for future mechanistic studies of CatC biology and for understanding the molecular consequences of therapeutic CatC inhibition.

## Supporting information

Suppementary Figures

Original gels

## List of Supplementary Materials

### Supplementary Tables

**Supplementary Table 1.** Protein, N-terminal and C-terminal peptide quantification and statistics of DMSO- and brensocatib-treated cells at 16 h, 72 h and 7 days

**Supplementary Table 2.** Common proteins significantly upregulated in the DMSO-treated cells at 72 h and 7 days

**Supplementary Table 3.** Common proteins significantly upregulated in the brensocatib-treated cells at 72 h and 7 days

**Supplementary Table 4.** Dipeptidyl peptidase signatures based on N-termini analysis in DMSO- and brensocatib-treated cells at 16 h, 72 h and 7 days

**Supplementary Table 5.** Summary C-terminal peptides significantly upregulated in DMSO- and brensocatib-treated cells at 7 days.

### Supplementary Figures

**Supplementary Figure 1.** Schematic of cathepsin C interaction with brensocatib, FY01 probe and an overview of the downstream proteomics workflow.

**Supplementary Figure 2.** Activity-based probe labelling and immunoblot of PR3 following brensocatib treatment

**Supplementary Figure 3.** PCA analysis of protein and peptide data in DMSO- and brensocatib-treated HL-60 cells at 16 h, 72 h and 7 days

**Supplementary Figure 4.** BMV109 probe labelling and immunoblot of cathepsin X and B following brensocatib treatment

**Supplementary Figure 5.** Dimethylation efficiency of HL-60 cells treated with DMSO or brensocatib (20 μM) for 16 h, 72 h, or 7 days

**Supplementary Figure 6.** Sample dimethylation efficiency and N-terminal modifications of HL-60 cells treated with DMSO or brensocatib

**Supplementary Figure 7.** Gene ontology of N-termini enriched in DMSO- or brensocatib-treated HL-60 cells for 72 h and 7 days

**Supplementary Figure 8.** Summary of cleavage distributions.

**Supplementary Figure 9.** Volcano plots of cleavage events classified by P1 residue in HL-60 cells treated with DMSO or brensocatib for 16 h, 72 h, or 7 days

**Supplementary Figure 10.** Heatmap of all peptides identified for the selected proteins at the 7-day timepoint, subset 1.

**Supplementary Figure 11.** Heatmap of all peptides identified for the selected proteins at the 7-day timepoint, subset 2.

**Supplementary Figure 12.** Heatmap of all peptides identified for the selected proteins at the 7-day timepoint, subset 3.

## DECLARATIONS

### Animal Ethics Committee Approval

Not applicable.

### Human Ethics Committee Approval

Not applicable.

### Author Contributions

Conceptualization: LEE-M, YZ, EKS-F, NES. Resources: EKS-F, LEE-M, NES. Writing - Original Draft: YZ. Writing -Review & Editing: YZ, LEE-M, EKS-F. Data Acquisition and Analysis: YZ, BX, SL, ARZ, KQY. Supervision: LEE-M, EKS-F. Funding acquisition: EKS-F, LEE-M.

## Acknowledgements

We thank the Mass Spectrometry and Proteomics Facility at the Bio21 Institute for their outstanding support. We thank E. Deu for providing the FY01 and K. Huang, A. Dobric and C.W. Armstrong for editorial feedback and scientific discussion.

## Availability of data and materials

The mass spectrometry proteomics and N-terminomics data have been deposited in the ProteomeXchange Consortium via the PRIDE partner repository with the data set identifiers PXD083578. Other data are provided herein, and additional information is available from the corresponding authors on request.

## Abbreviations

CatC: Cathepsin C
DPP-1: Dipeptidyl peptidase I
NE: Neutrophil elastase
PR3: Proteinase 3
Catg: Cathepsin G
CatX: Cathepsin X
CatB: Cathespin B
CatS: Cathepsin S
DIA: data-independent acquisition
GO: Gene ontology
NICE: No-enrichment identification of Cleavage events
NSP: Neutrophil serine proteases

