## Supplementary material for "Temporal N-terminomics analysis reveals proteome remodeling following brensocatib-mediated cathepsin C inhibition in promyeloblast cells": Suppementary Figures

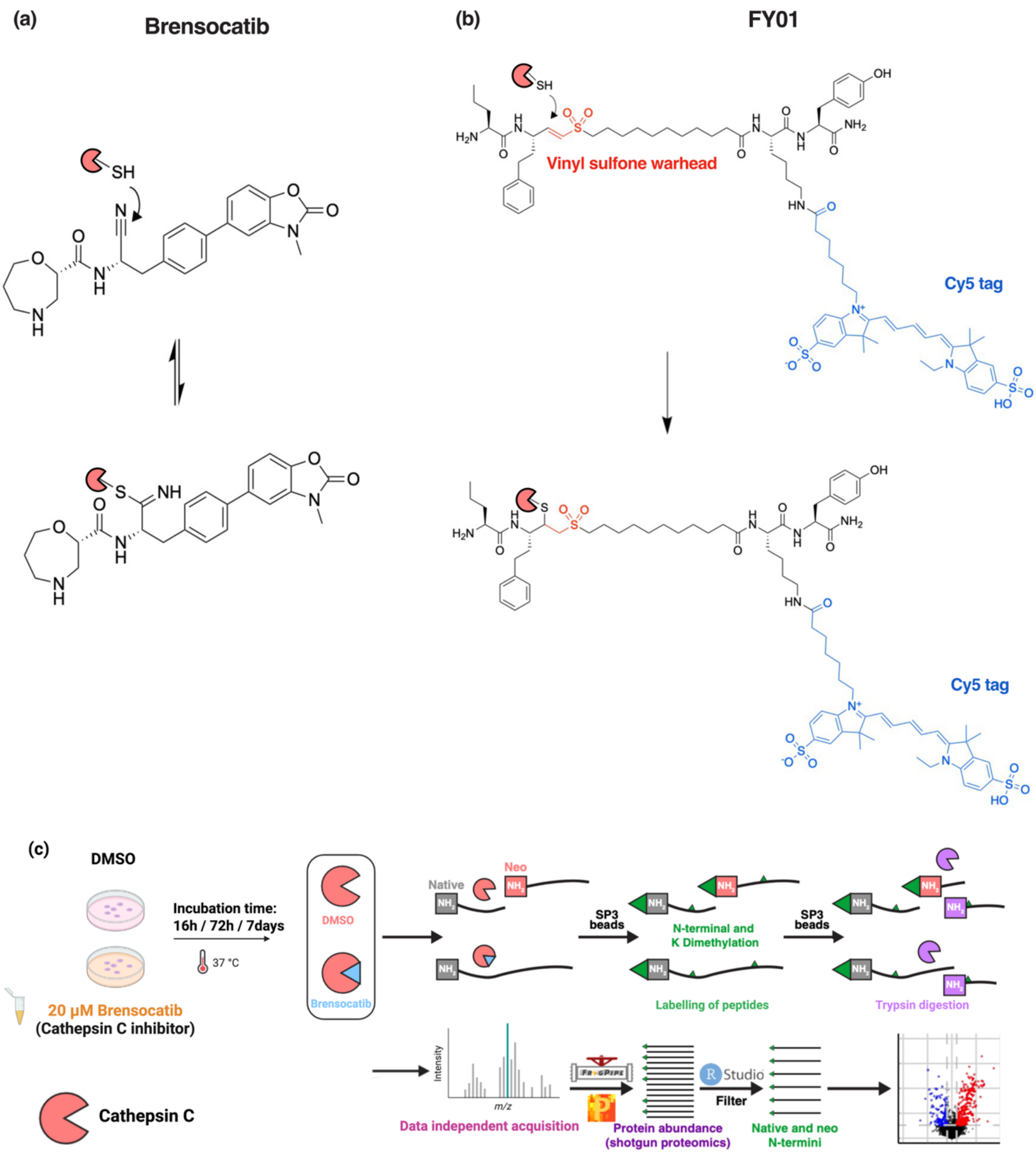


**Supplementary Figure 1. Schematic of cathepsin C interaction with brensocatib, FY01 probe and an overview of the downstream proteomics workflow.** (a) Interaction of the brensocatib nitrile warhead with the active-site cysteine residue of cathepsin C that forms a reversible thioimidate complex. (b) Structure of the activity-based probe FY01 and its covalent irreversible interaction cathepsin C through the vinyl sulfone warhead. FY01 probe has a Cy5 tag that allows subsequent quantification of active cathepsin C by fluorescence-based measurements. **(c)** Overview of the proteomics workflow used to characterize cathepsin C-dependent proteolytic changes, figure adapted from Ziegler *et al*. (2025).

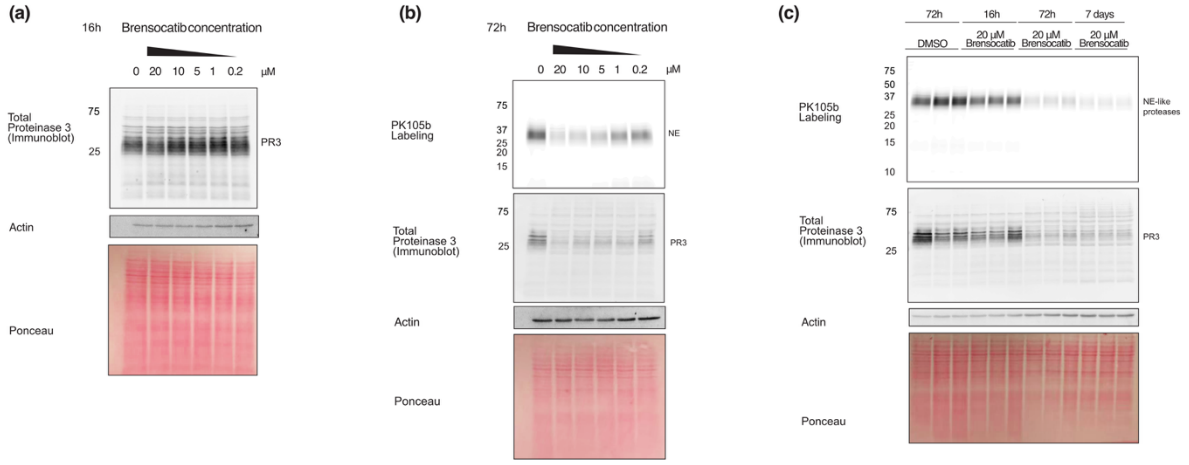


**Supplementary Figure 2**. **Total and active PR3 are reduced by brensocatib treatment in HL-60 cells.** PK105b labelling was used to identify elastase-like protease activity, and PR3 antibodies were used for immunoblot to detect total PR3 levels. HL-60 cells were treated with DMSO (vehicle control) or brensocatib (0.2, 1, 5, 10, 20 µM) for **(a)** 16 h or **(b)** 72 h. **(c)** HL-60 cells were treated with DMSO (vehicle control) or brensocatib (20 µM) for 16 h, 72 h, or 7 days followed by PK105b labelling and PR3 immunoblot (n=3/group).

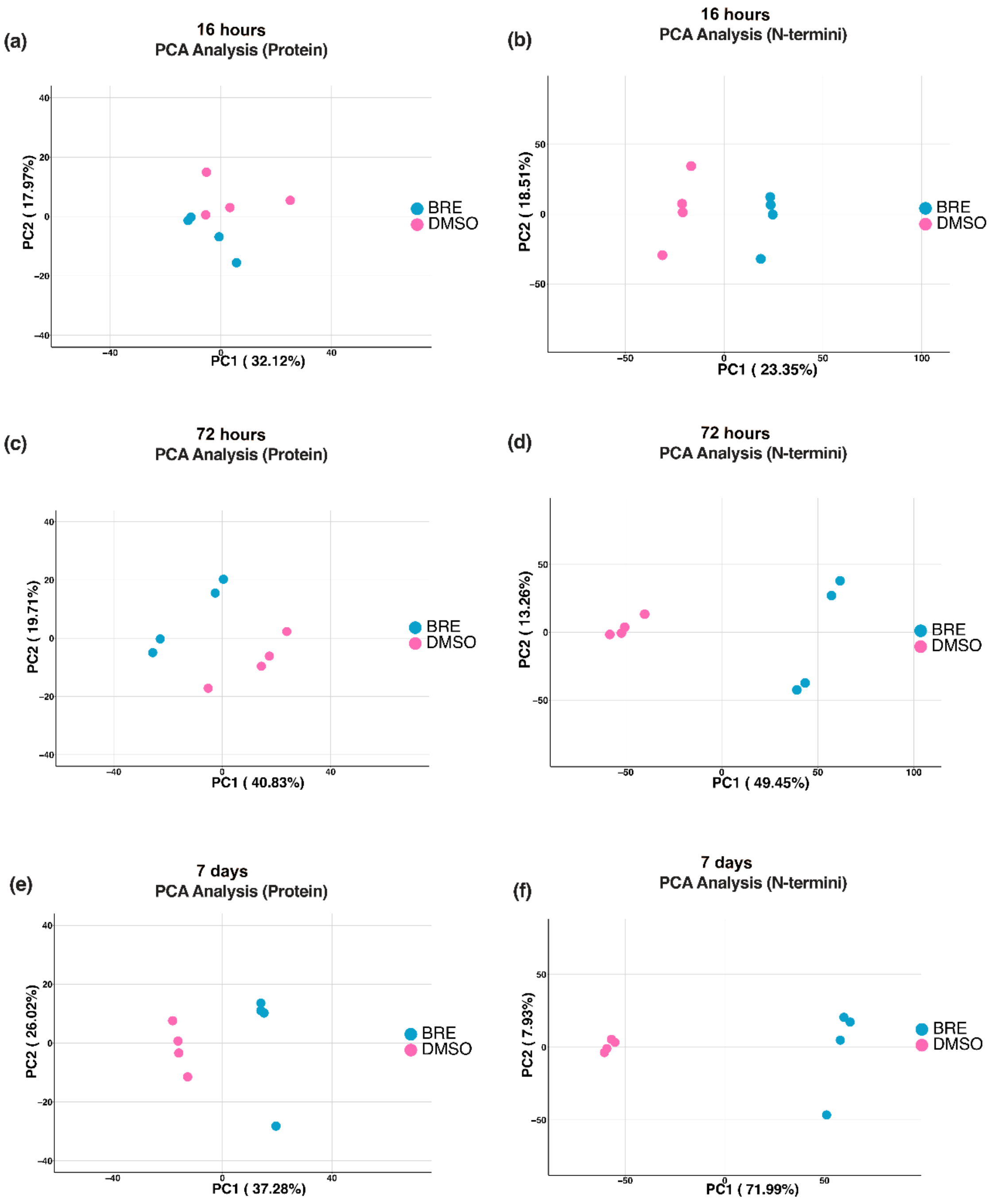


**Supplementary Figure 3. NICE analysis of DMSO- and brensocatib-treated HL-60 lysates reveals clear time-dependent clustering of replicate samples.** Principal component analysis (PCA) of global protein abundance and N-termini at 16 h (a, b), 72 h (c, d) and 7 days (e,f). DMSO data are shown in pink and brensocatib data in blue (n=4/group)

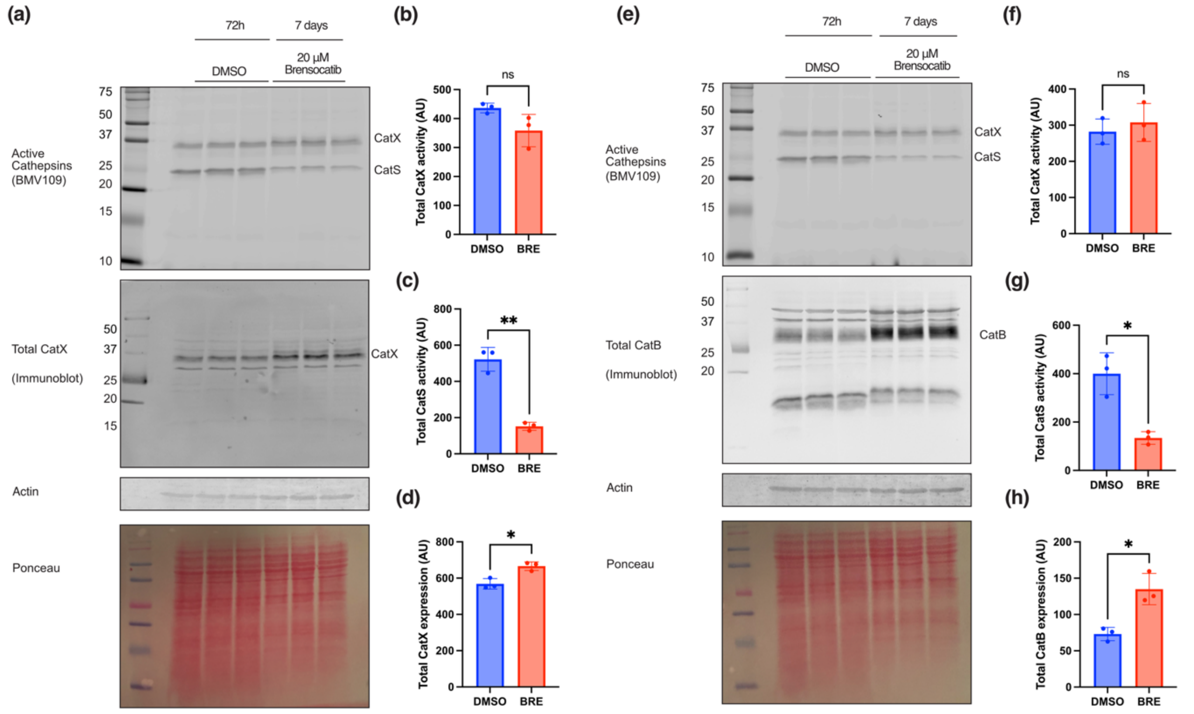


**Supplementary Figure 4. Total CatX and CatB expression are increased by brensocatib treatment in HL-60 cells.** HL-60 cells were treated with DMSO (vehicle control) or brensocatib (20 µM) for 72 h or 7 days (n=3/group). **(a,e)** The BMV109 activity-based probe was used to detect active cathepsins and visualized by in-gel fluorescence, followed by immunoblotting to measure total protease levels. Densitometry of active CatX **(b,f),** active CatS **(c,g),** total CatX **(d),** and total CatB **(h)** are graphed. Ns $p > 0.05$, *$p < 0.05$, **$p < 0.01$

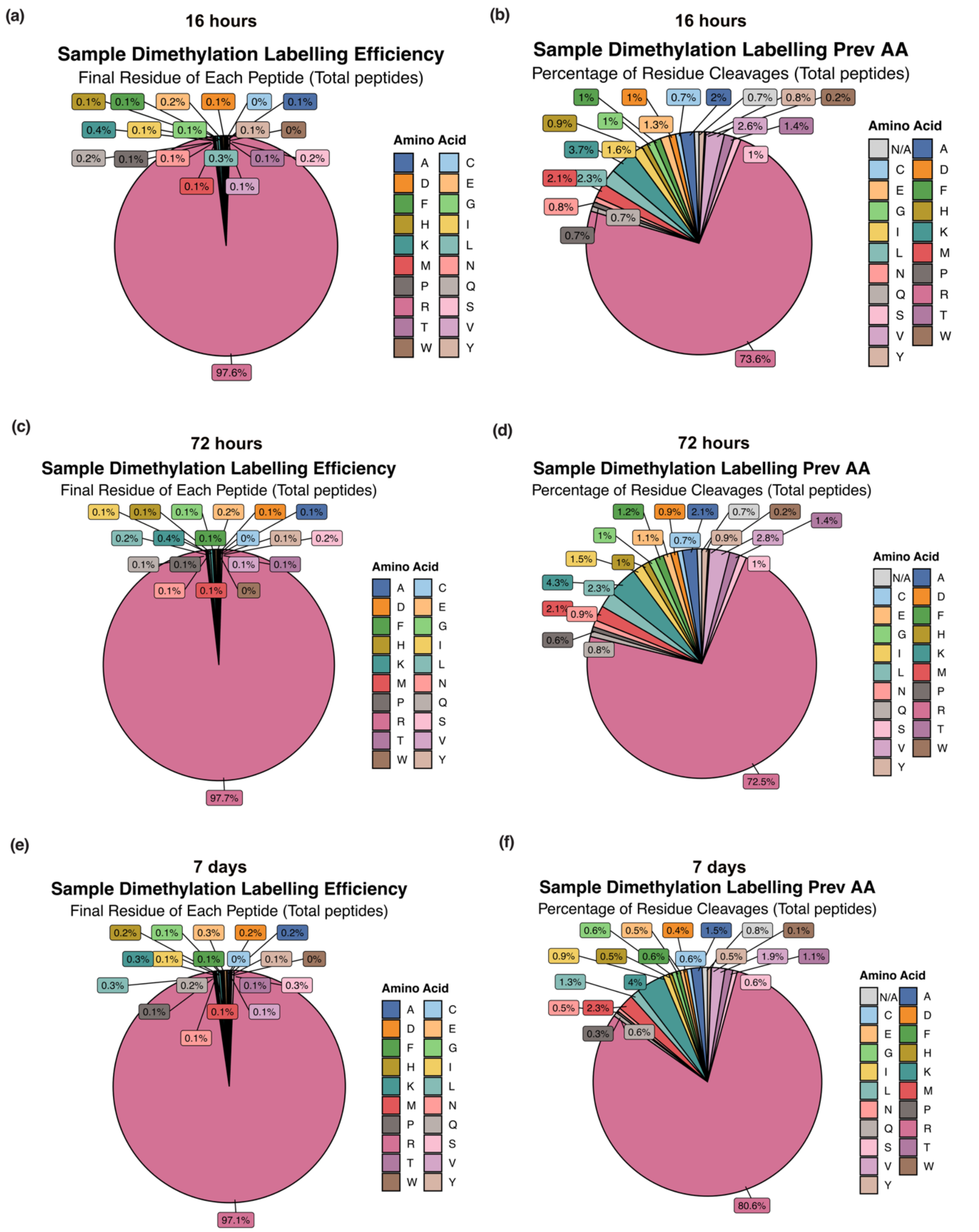


**Supplementary Figure 5. Dimethylation efficiency of HL-60 cells treated with DMSO or brensocatib (20 µM) for 16 h, 72 h, or 7 days.** Samples were reduced and alkylated before dimethylation of the N-termini and lysine side chains and subsequent LC-MS/MS analysis. Amino acid distribution at the end of all identified peptides (left) and amino acid residues prior to the identified N-terminal peptides (right) at 16 h **(a,b)**, 72 h **(c,d)**, and 7 days **(e,f)** were used as measures of dimethylation efficiency.

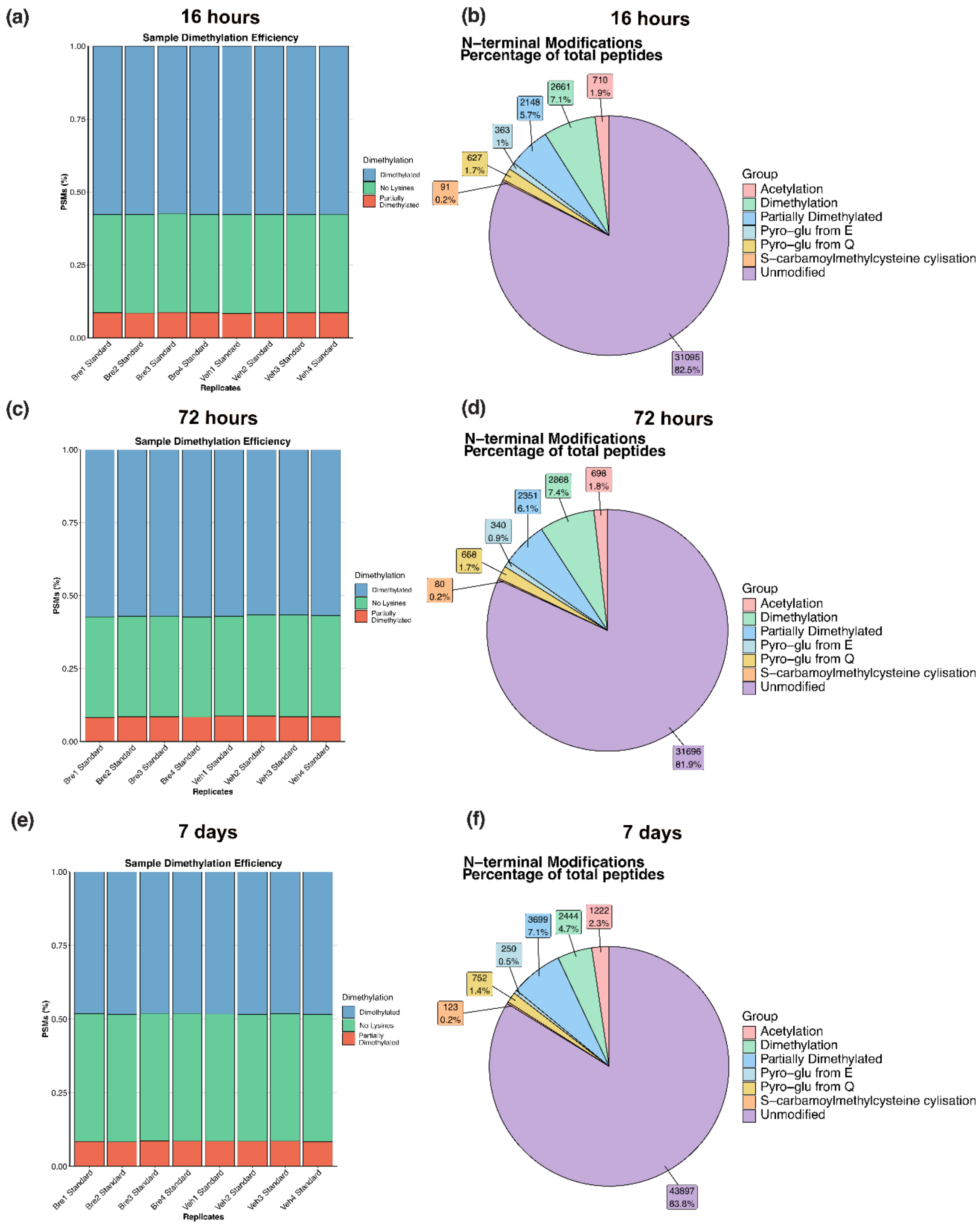


**Supplementary Figure 6. Sample dimethylation efficiency and N-terminal modifications of HL-60 cells treated with DMSO or brensocatib (20 µM) for 16 h, 72 h, or 7 days.** Samples were reduced and alkylated before dimethylation of N-termini and lysine side chains. Dimethylation efficiency of each biological replicate based on lysine labelling status (all dimethylated [blue], partially dimethylated [red], or no lysine [green]) at 16 h **(a),** 72 h **(c),** and 7 days **(e)** using all detected peptides. The pie chart shows the distribution of N-terminal modifications in the normalised and combined precursors across the replicates from each dataset: 16 h **(b),** 72 h **(d)**, and 7 days **(f)** including N-terminal demethylated peptides.

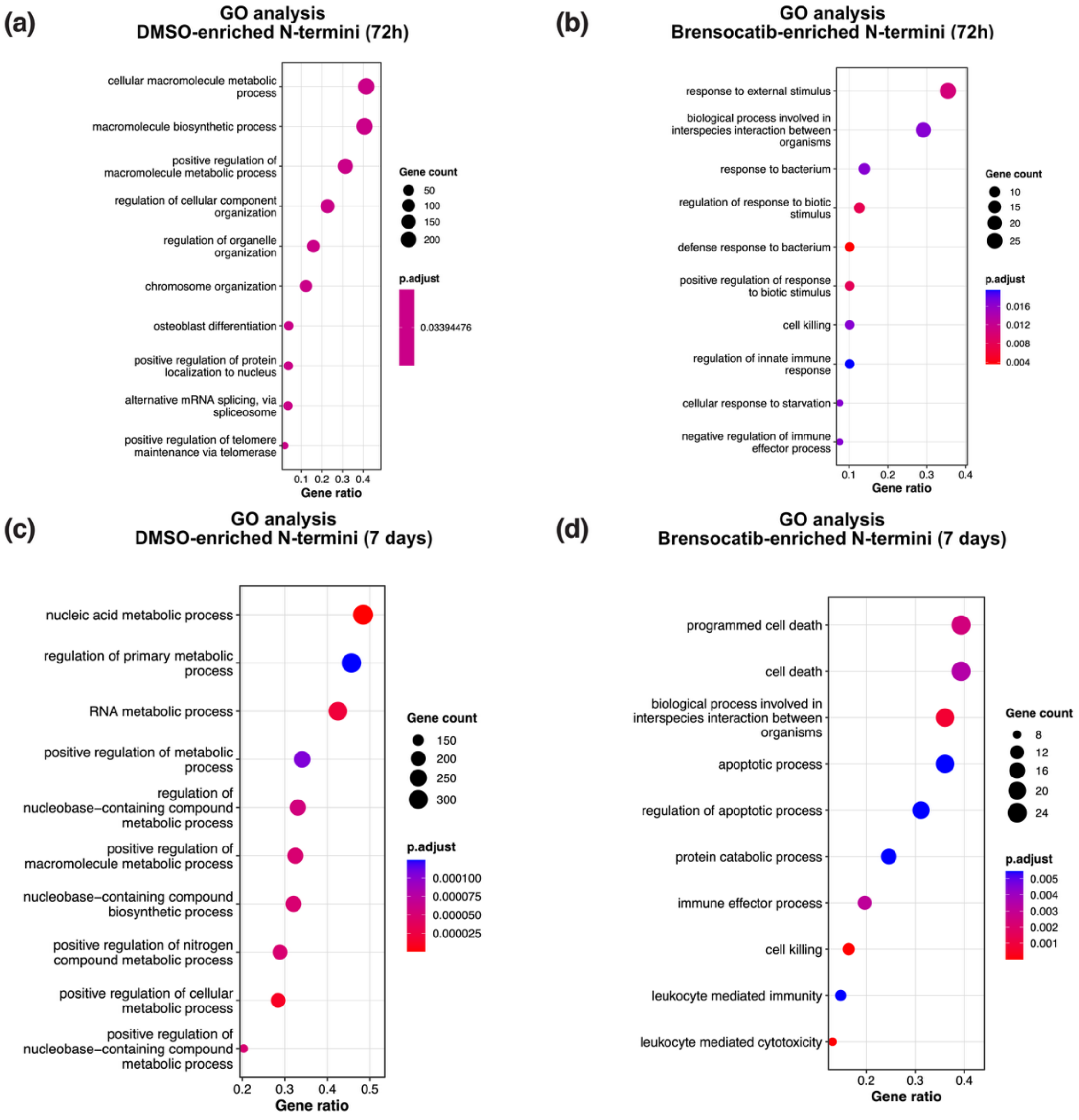


**Supplementary Figure 7. Gene ontology analysis of N-termini enriched in DMSO- or brensocatib (20 µM)-treated HL-60 cells for 72 h and 7 days (n=4/group).** Gene ontology biological processes enrichment dot plot indicates differentially expressed N-termini detected in the DMSO-treated group and brensocatib-treated cells after 72 h **(a-b)**, and 7 days **(c-d).** All peptides identified in the corresponding datasets were used as the background universe for each enrichment analysis.

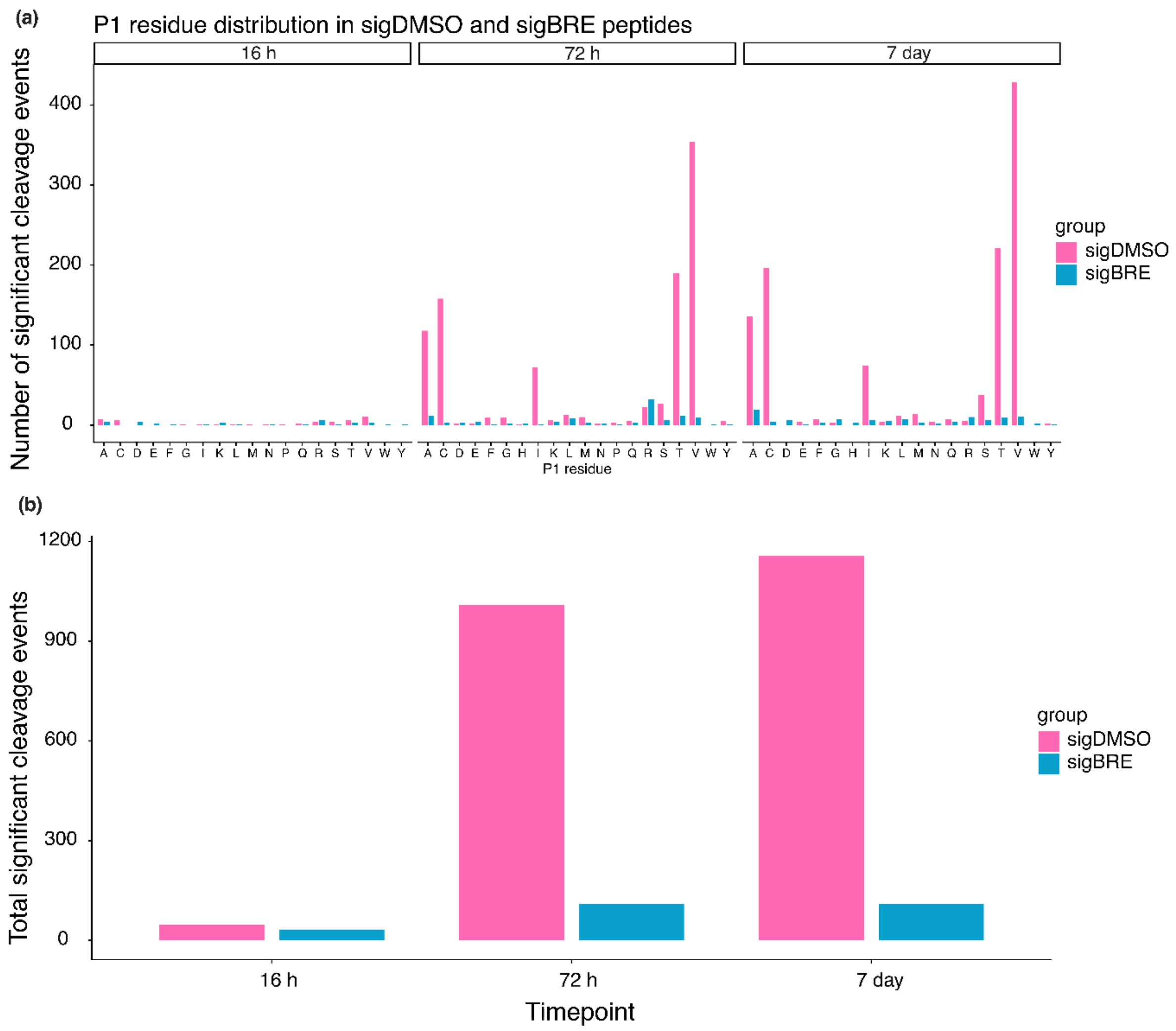


**Supplementary Figure 8. Summary of cleavage distributions.** HL-60 cells were treated with DMSO or brensocatib (20 μM) for 16 h, 72 h and 7 days (n=4/group), neo-N-termini were filtered and identified following filtering. **(a)** Bar plot showing the distribution of P1 residues in the significantly upregulated N-terminal peptides identified at 16 h, 72 h, and 7 days in the DMSO-treated (pink) and brensocatib-treated (blue) cells. **(b)** Total number of differentially expressed N-terminal peptides significantly increased in either DMSO- or brensocatib-treated cells at each timepoint.

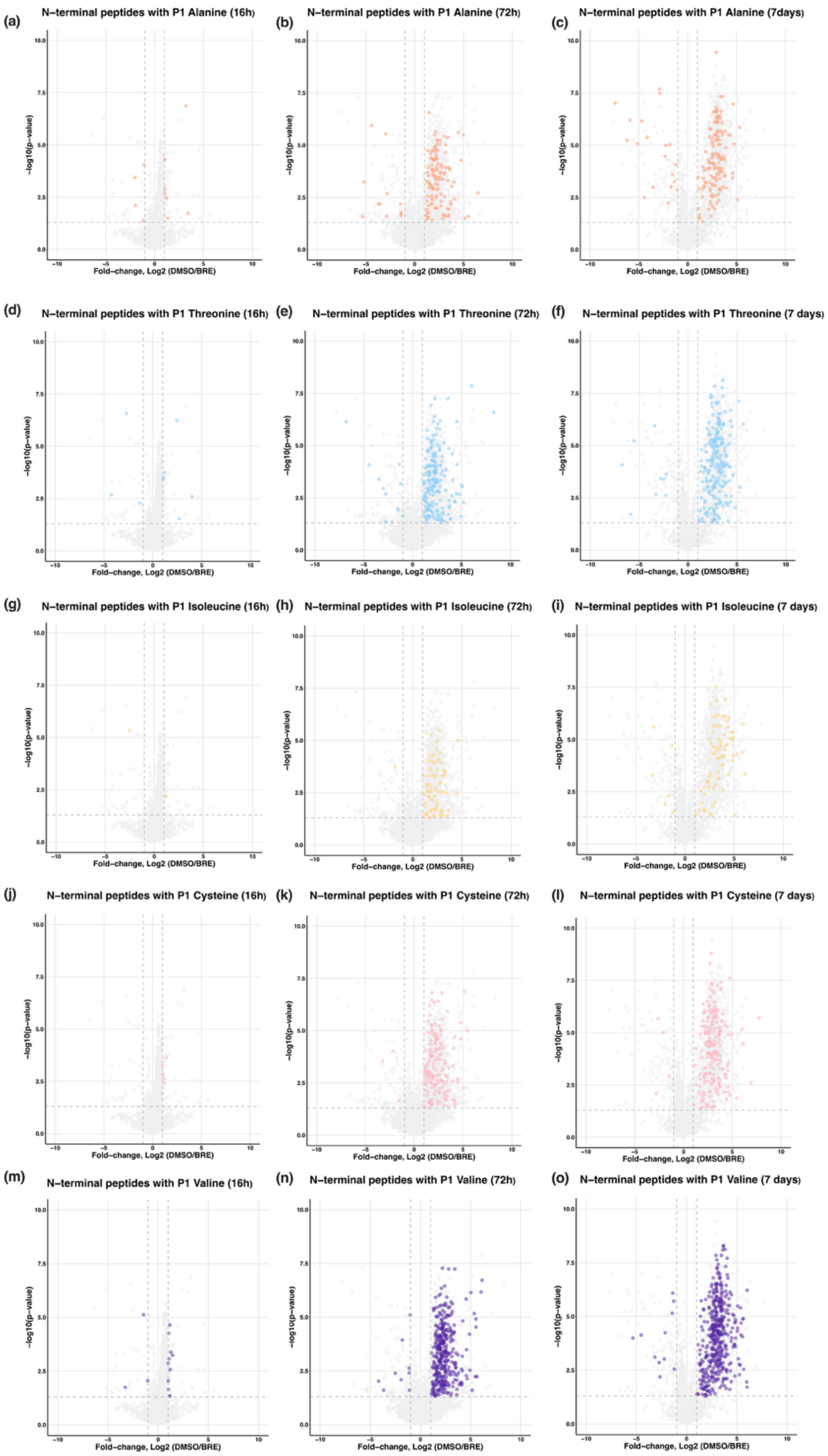


**Supplementary Figure 9. Volcano plots highlighting cleavage events classified by P1 residue in HL-60 cells treated with DMSO or brensocatib (20 µM) for 16 h, 72 h, or 7 days.** Cleavage events are shown for P1 residues: **(a–c)** alanine, **(d–f)** threonine, **(g–i)** isoleucine, **(j–l)** cysteine, and **(m-o)** valine.

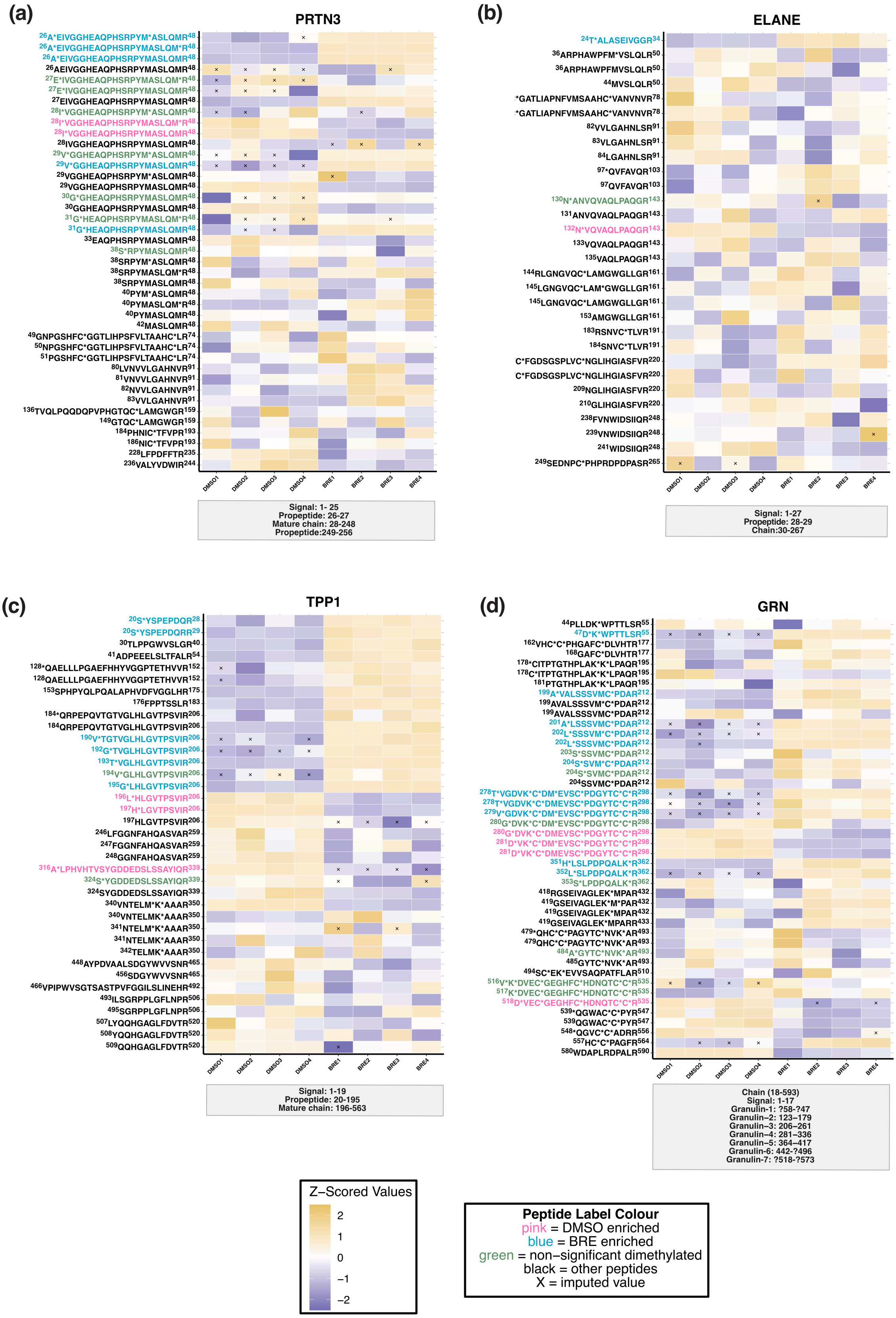
(a)
PRTN3
26A*EIVGGHEAQPHSRPYM*ASLQMR48
26A*EIVGGHEAQPHSRPYMASLQM*R48
26A*EIVGGHEAQPHSRPYMASLQMR48
26AEIVGGHEAQPHSRPYMASLQMR48
27E*IVGGHEAQPHSRPYMASLQM*R48
27E*IVGGHEAQPHSRPYMASLQMR48
27EIVGGHEAQPHSRPYMASLQMR48
28I*VGGHEAQPHSRPYM*ASLQMR48
28I*VGGHEAQPHSRPYMASLQM*R48
28I*VGGHEAQPHSRPYMASLQMR48
28IVGGHEAQPHSRPYMASLQMR48
29V*GGHEAQPHSRPYM*ASLQMR48
29V*GGHEAQPHSRPYMASLQMR48
29VGGHEAQPHSRPYM*ASLQMR48
29VGGHEAQPHSRPYMASLQMR48
30G*GHEAQPHSRPYMASLQMR48
30GGHEAQPHSRPYMASLQMR48
31G*HEAQPHSRPYMASLQM*R48
31G*HEAQPHSRPYMASLQMR48
33EAQPHSRPYMASLQMR48
38S*RPYMASLQMR48
38SRPYM*ASLQMR48
38SRPYMASLQM*R48
38SRPYMASLQMR48
40PYM*ASLQMR48
40PYMASLQM*R48
40PYMASLQMR48
42MASLQMR48
49GNPGSHFC*GGTLIHPSFVLTAAHC*LR74
50NPGSHFC*GGTLIHPSFVLTAAHC*LR74
51PGSHFC*GGTLIHPSFVLTAAHC*LR74
80LVNVVLGAHNVR91
81VNVVLGAHNVR91
82NVVLGAHNVR91
83VVLGAHNVR91
136TVQLPQQDQPVPHGTQC*LAMGWGR159
149GTQC*LAMGWGR159
184PHNIC*TFVPR193
186NIC*TFVPR193
228LFPDFFTR235
236VALYVDWIR244
DMSO1 DMSO2 DMSO3 DMSO4 BRE1 BRE2 BRE3 BRE4
Signal: 1- 25
Propeptide: 26-27
Mature chain: 28-248
Propeptide:249-256
(b)
ELANE
24T*ALASEIVGGR34
36ARPHAWPFM*VSLQLR50
36ARPHAWPFMVSLQLR50
44MVSLQLR50
*GATLIAPNFVMSAAHC*VANVNVR78
*GATLIAPNFVMSAAHC*VANVNVR78
82VVLGAHNLSR91
83VLGAHNLSR91
84LGAHNLSR91
97*QVFAVQR103
97QVFAVQR103
130N*ANVQVAQLPAQGR143
131ANVQVAQLPAQGR143
132N*VQVAQLPAQGR143
133VQVAQLPAQGR143
135VAQLPAQGR143
144RLGNGVQC*LAMGWGLLGR161
145LGNGVQC*LAM*GWGLLGR161
145LGNGVQC*LAMGWGLLGR161
153AMGWGLLGR161
183RSNVC*TLVR191
184SNVC*TLVR191
C*FGDSGSPLVC*NGLIHGIASFVR220
C*FGDSGSPLVC*NGLIHGIASFVR220
209NGLIHGIASFVR220
210GLIHGIASFVR220
238FVNWIDSIIQR248
239VNWIDSIIQR248
241WIDSIIQR248
249SEDNPC*PHPRDPDPASR265
DMSO1 DMSO2 DMSO3 DMSO4 BRE1 BRE2 BRE3 BRE4
Signal: 1-27
Propeptide: 28-29
Chain:30-267
(c)
TPP1
20S*YSPEPDQR28
20S*YSPEPDQRR29
30TLPPGWVSLGR40
41ADPEEELSLTFALR54
128*QAELLLPGAEFHHYVGGPTETHVVR152
128QAELLLPGAEFHHYVGGPTETHVVR152
153SPHPYQLPQALAPHVDFVGGLHR175
176FPPTSSLR183
184*QRPEPQVTGTVGLHLGVTPSVIR206
184QRPEPQVTGTVGLHLGVTPSVIR206
190V*TGTVGLHLGVTPSVIR206
192G*TVGLHLGVTPSVIR206
193T*VGLHLGVTPSVIR206
194V*GLHLGVTPSVIR206
195G*LHLGVTPSVIR206
196L*HLGVTPSVIR206
197H*LGVTPSVIR206
197HLGVTPSVIR206
246LFGGNFAHQASVAR259
247FGGNFAHQASVAR259
248GGNFAHQASVAR259
316A*LPHVHTVSYGDDEDSLSSAYIQR339
324S*YGDDEDSLSSAYIQR339
324SYGDDEDSLSSAYIQR339
340VNTELM*K*AAAR350
340VNTELMK*AAAR350
341NTELM*K*AAAR350
341NTELMK*AAAR350
342TELMK*AAAR350
448AYPDVAALSDGYWVVSNR465
456SDGYWVVSNR465
466VPIPWVSGTSASTPVFGGILSLINEHR492
493ILSGRPPLGFLNPR506
495SGRPPLGFLNPR506
507LYQQHGAGLFDVTR520
508YQQHGAGLFDVTR520
509QQHGAGLFDVTR520
DMSO1 DMSO2 DMSO3 DMSO4 BRE1 BRE2 BRE3 BRE4
Signal: 1-19
Propeptide: 20-195
Mature chain: 196-563
(d)
GRN
44PLLDK*WPTTLSR55
47D*K*WPTTLSR55
162VHC*C*PHGAFC*DLVHTR177
168GAFC*DLVHTR177
178*CITPTGTHPLAK*K*LPAQR195
178C*ITPTGTHPLAK*K*LPAQR195
181PTGTHPLAK*K*LPAQR195
199A*VALSSSVMC*PDAR212
199AVALSSSVM*C*PDAR212
199AVALSSSVMC*PDAR212
201A*LSSSVMC*PDAR212
202L*SSSVM*C*PDAR212
202L*SSSVMC*PDAR212
203S*SSVMC*PDAR212
204S*SVM*C*PDAR212
204S*SVMC*PDAR212
204SSVMC*PDAR212
278T*VGDVK*C*DM*EVSC*PDGYTC*C*R298
278T*VGDVK*C*DMEVSC*PDGYTC*C*R298
279V*GDVK*C*DM*EVSC*PDGYTC*C*R298
280G*DVK*C*DM*EVSC*PDGYTC*C*R298
280G*DVK*C*DMEVSC*PDGYTC*C*R298
281D*VK*C*DM*EVSC*PDGYTC*C*R298
281D*VK*C*DMEVSC*PDGYTC*C*R298
351H*LSLPDPQALK*R362
352L*SLPDPQALK*R362
353S*LPDPQALK*R362
418RGSEIVAGLEK*MPAR432
419GSEIVAGLEK*M*PAR432
419GSEIVAGLEK*MPAR432
419GSEIVAGLEK*MPARR433
479*QHC*C*PAGYTC*NVK*AR493
479QHC*C*PAGYTC*NVK*AR493
484A*GYTC*NVK*AR493
485GYTC*NVK*AR493
494SC*EK*EVSAQPATFLAR510
516V*K*DVEC*GEGHFC*HDNQTC*C*R535
517K*DVEC*GEGHFC*HDNQTC*C*R535
518D*VEC*GEGHFC*HDNQTC*C*R535
539*QGWAC*C*PYR547
539QGWAC*C*PYR547
548*QGVC*C*ADRR556
557HC*C*PAGFR564
580WDAPLRDPALR590
DMSO1 DMSO2 DMSO3 DMSO4 BRE1 BRE2 BRE3 BRE4
Chain (18-593)
Signal: 1-17
Granulin-1: ?58-?47
Granulin–2: 123–179
Granulin–3: 206–261
Granulin–4: 281–336
Granulin–5: 364–417
Granulin-6: 442-?496
Granulin-7: ?518-?573
Z–Scored Values
2
1
0
−1
−2
Peptide Label Colour
pink = DMSO enriched
blue = BRE enriched
green = non–significant dimethylated
black = other peptides
X = imputed value

**Supplementary Figure 10. Heatmap of all peptides identified for the selected proteins at the 7-day timepoint.** Z-scores were calculated from the intensity of each peptide (yellow = increased abundance, purple = decreased abundance). Neo-N-terminal peptides significantly enriched in DMSO- or brensocatib-treated samples are shown in pink and blue, respectively; non-significant neo-N-termini peptides are shown in green. Other detected non-dimethylated peptides are shown in black. Variable modifications were replaced by an asterisk (*) in the peptide sequence. Crosses indicate the sample had imputed intensity values. The proteins shown are **(a)** proteinase 3, **(b)** elastase, **(c)** tripeptidyl peptidase 1 and **(d)** progranulin.

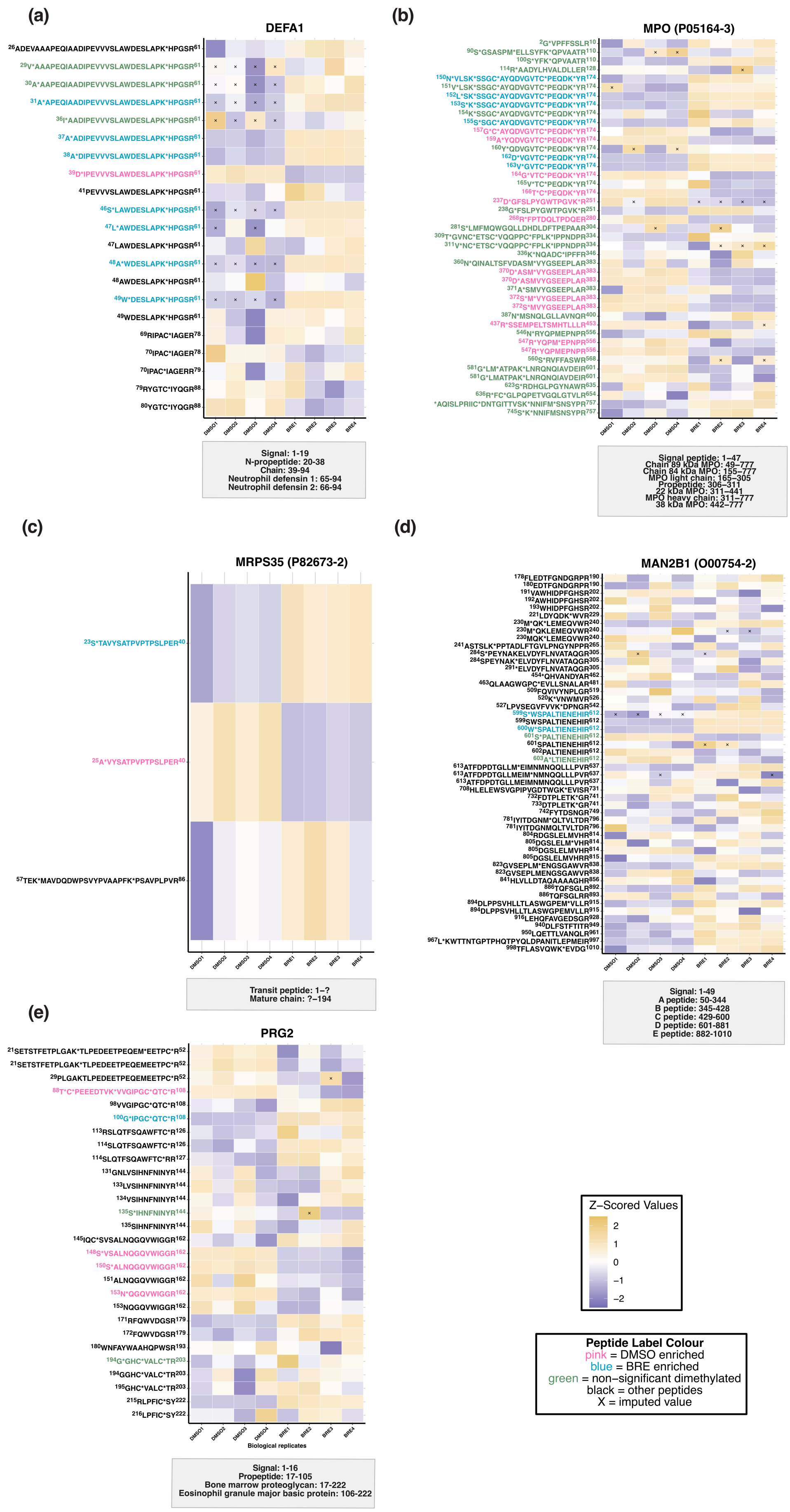
(a)
DEFA1
DMSO1
DMSO2
DMSO3
DMSO4
BRE1
BRE2
BRE3
BRE4
Signal: 1-19
N-propeptide: 20-38
Chain: 39-94
Neutrophil defensin 1: 65-94
Neutrophil defensin 2: 66-94
(b)
MPO (P05164-3)
Signal peptide: 1–47
Chain 89 kDa MPO: 49–777
Chain 84 kDa MPO: 155–777
MPO light chain: 165–305
Propeptide: 306–311
22 kDa MPO: 311–441
MPO heavy chain: 311–777
38 kDa MPO: 442–777
(c)
MRPS35 (P82673-2)
Transit peptide: 1–?
Mature chain: ?–194
(d)
MAN2B1 (O00754-2)
Signal: 1-49
A peptide: 50-344
B peptide: 345-428
C peptide: 429-600
D peptide: 601-881
E peptide: 882-1010
(e)
PRG2
Biological replicates
Signal: 1-16
Propeptide: 17-105
Bone marrow proteoglycan: 17-222
Eosinophil granule major basic protein: 106-222
Z–Scored Values
2
1
0
−1
−2
Peptide Label Colour
pink = DMSO enriched
blue = BRE enriched
green = non–significant dimethylated
black = other peptides
X = imputed value

**Supplementary Figure 11. Heatmap of all peptides identified for selected proteins at 7-day timepoint.** Z-scores were calculated from the intensity of each peptide (yellow = increased abundance, purple = decreased abundance). Neo-N-terminal peptides significantly enriched in DMSO- or brensocatib-treated samples are shown in pink and blue, respectively; non-significant neo-N-termini peptides are shown in green. Other detected non-dimethylated peptides are shown in black. Variable modifications were replaced by an asterisk (*) in the peptide sequence. Crosses indicate the sample had imputed intensity values. The proteins shown are **(a)** defensin 1, **(b)** myeloperoxidase (using only dimethylated peptides), **(c)** mitochondrial small ribosomal subunit protein mS35, **(d)** lysosomal alpha-mannosidase, and **(e)** proteoglycan 2.

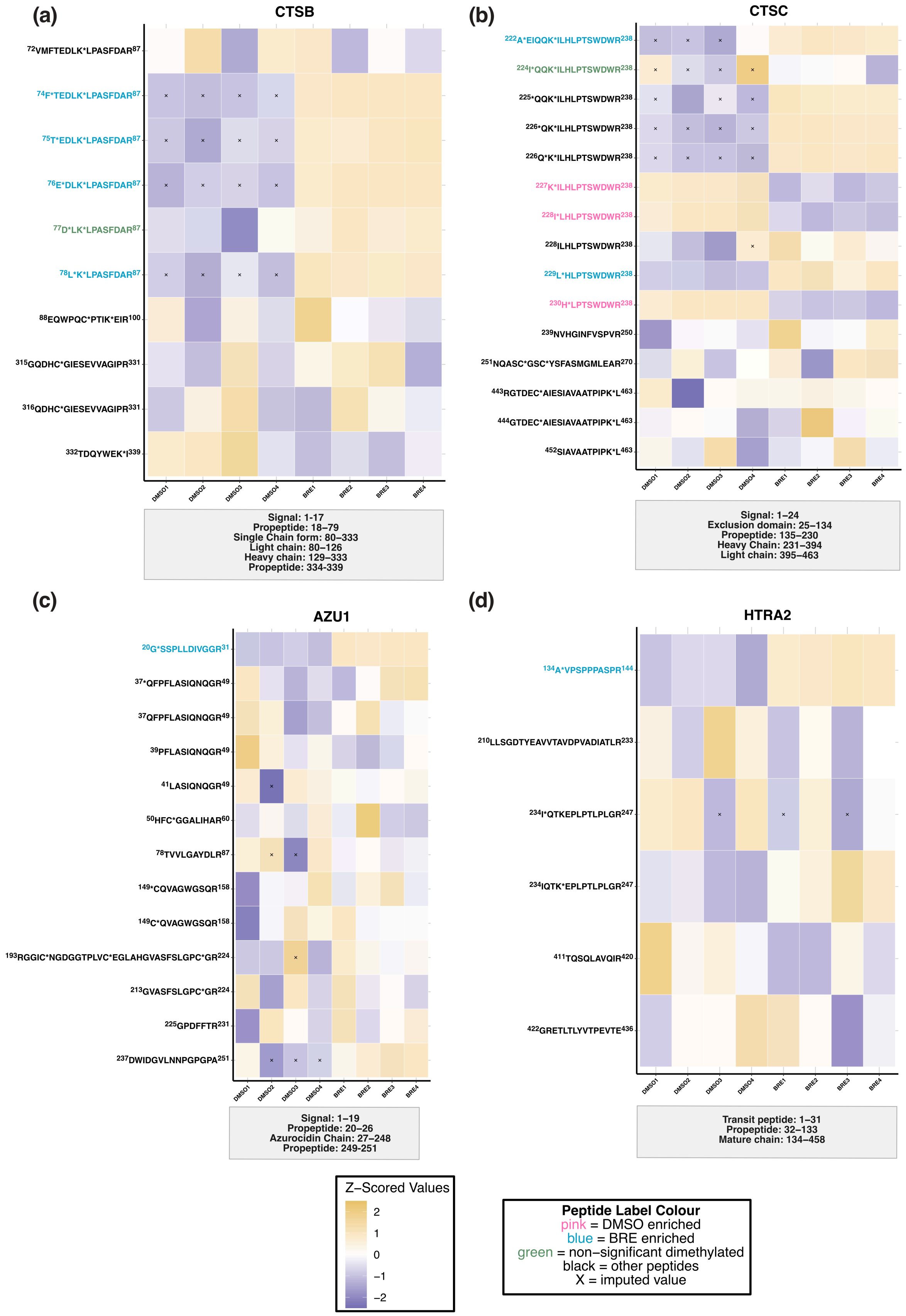

(a)
CTSB
72VMFTEDLK*LPASFDAR87
74F*TEDLK*LPASFDAR87
75T*EDLK*LPASFDAR87
76E*DLK*LPASFDAR87
77D*LK*LPASFDAR87
78L*K*LPASFDAR87
88EQWPQC*PTIK*EIR100
315GQDHC*GIESEVVAGIPR331
316QDHC*GIESEVVAGIPR331
332TDQYWEK*I339
DMSO1 DMSO2 DMSO3 DMSO4 BRE1 BRE2 BRE3 BRE4
Signal: 1-17
Propeptide: 18–79
Single Chain form: 80–333
Light chain: 80–126
Heavy chain: 129–333
Propeptide: 334-339
(b)
CTSC
222A*EIQQK*ILHLPTSWDWR238
224I*QQK*ILHLPTSWDWR238
225*QQK*ILHLPTSWDWR238
226*QK*ILHLPTSWDWR238
226Q*K*ILHLPTSWDWR238
227K*ILHLPTSWDWR238
228I*LHLPTSWDWR238
228ILHLPTSWDWR238
229L*HLPTSWDWR238
230H*LPTSWDWR238
239NVHGINFVSPVR250
251NQASC*GSC*YSFASMGMLEAR270
443RGTDEC*AIESIAVAATPIPK*L463
444GTDEC*AIESIAVAATPIPK*L463
452SIAVAATPIPK*L463
DMSO1 DMSO2 DMSO3 DMSO4 BRE1 BRE2 BRE3 BRE4
Signal: 1–24
Exclusion domain: 25–134
Propeptide: 135–230
Heavy Chain: 231–394
Light chain: 395–463
(c)
AZU1
20G*SSPLLDIVGGR31
37*QFPFLASIQNQGR49
37QFPFLASIQNQGR49
39PFLASIQNQGR49
41LASIQNQGR49
50HFC*GGALIHAR60
78TVVLGAYDLR87
149*CQVAGWGSQR158
149C*QVAGWGSQR158
193RGGIC*NGDGGTPLVC*EGLAHGVASFSLGPC*GR224
213GVASFSLGPC*GR224
225GPDFFTR231
237DWIDGVLNNPGPGPA251
DMSO1 DMSO2 DMSO3 DMSO4 BRE1 BRE2 BRE3 BRE4
Signal: 1–19
Propeptide: 20–26
Azurocidin Chain: 27–248
Propeptide: 249-251
(d)
HTRA2
134A*VPSPPPASPR144
210LLSGDTYEAVVTAVDPVADIATLR233
234I*QTKEPLPTLPLGR247
234IQTK*EPLPTLPLGR247
411TQSQLAVQIR420
422GRETLTLYVTPEVTE436
DMSO1 DMSO2 DMSO3 DMSO4 BRE1 BRE2 BRE3 BRE4
Transit peptide: 1–31
Propeptide: 32–133
Mature chain: 134–458
Z–Scored Values
2
1
0
−1
−2
Peptide Label Colour
pink = DMSO enriched
blue = BRE enriched
green = non–significant dimethylated
black = other peptides
X = imputed value

**Supplementary Figure 12.** Heatmap of all peptides identified for selected proteins at 7-day timepoint. Z-scores were calculated from the intensity of each peptide (yellow = increased abundance, purple = decreased abundance). Neo-N-terminal peptides significantly enriched in DMSO- or brensocatib-treated samples are shown in pink and blue, respectively; non-significant neo-N-terminal peptides are shown in green. Other detected non-dimethylated peptides are shown in black. Variable modifications were replaced by an asterisk (*) in the peptide sequence. Crosses indicate the sample had imputed intensity values. The proteins shown are **(a)** cathepsin B, **(b)** cathepsin C, **(c)** azurocidin-1 and **(d)** HTRA2.
