## Supplementary material for "Temporal N-terminomics analysis reveals proteome remodeling following brensocatib-mediated cathepsin C inhibition in promyeloblast cells": Original gels

**Fig1a**

**FY01 Labelling**

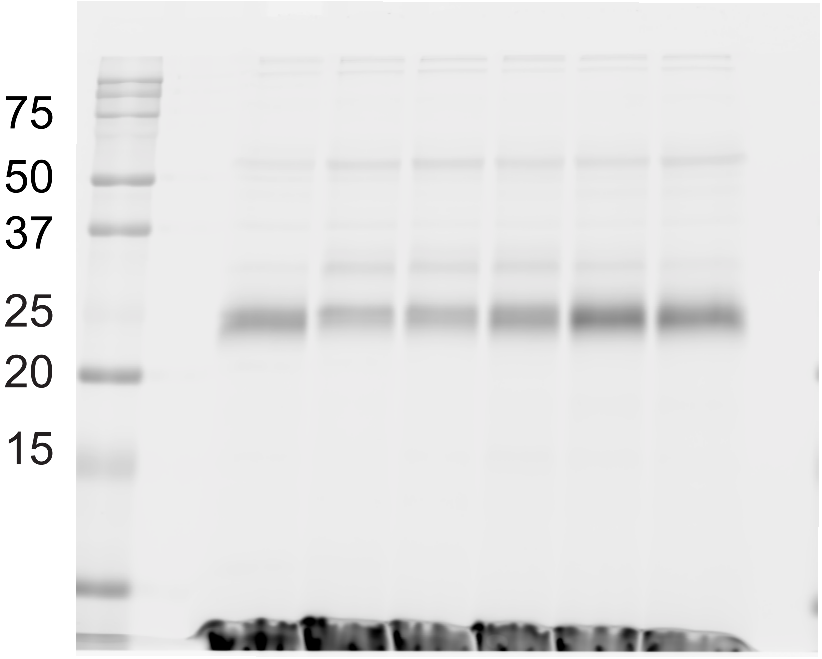


**CatC Immunoblot**

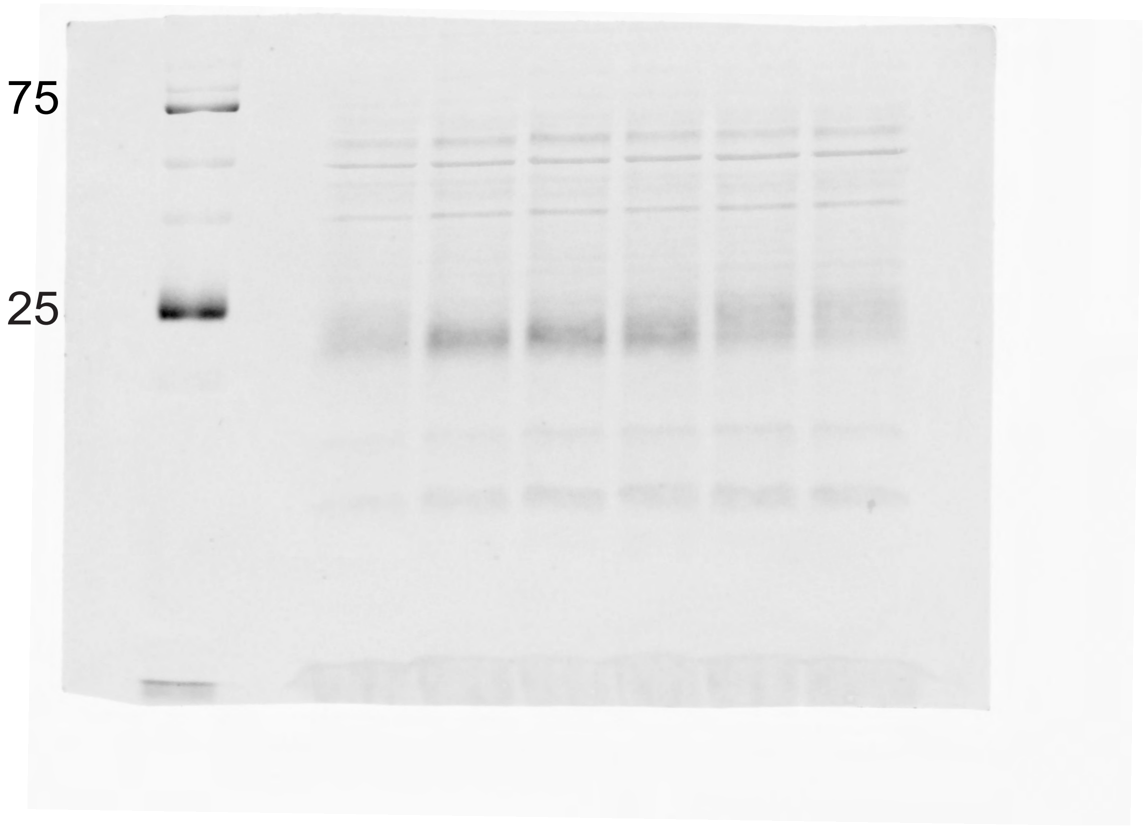

**Actin**

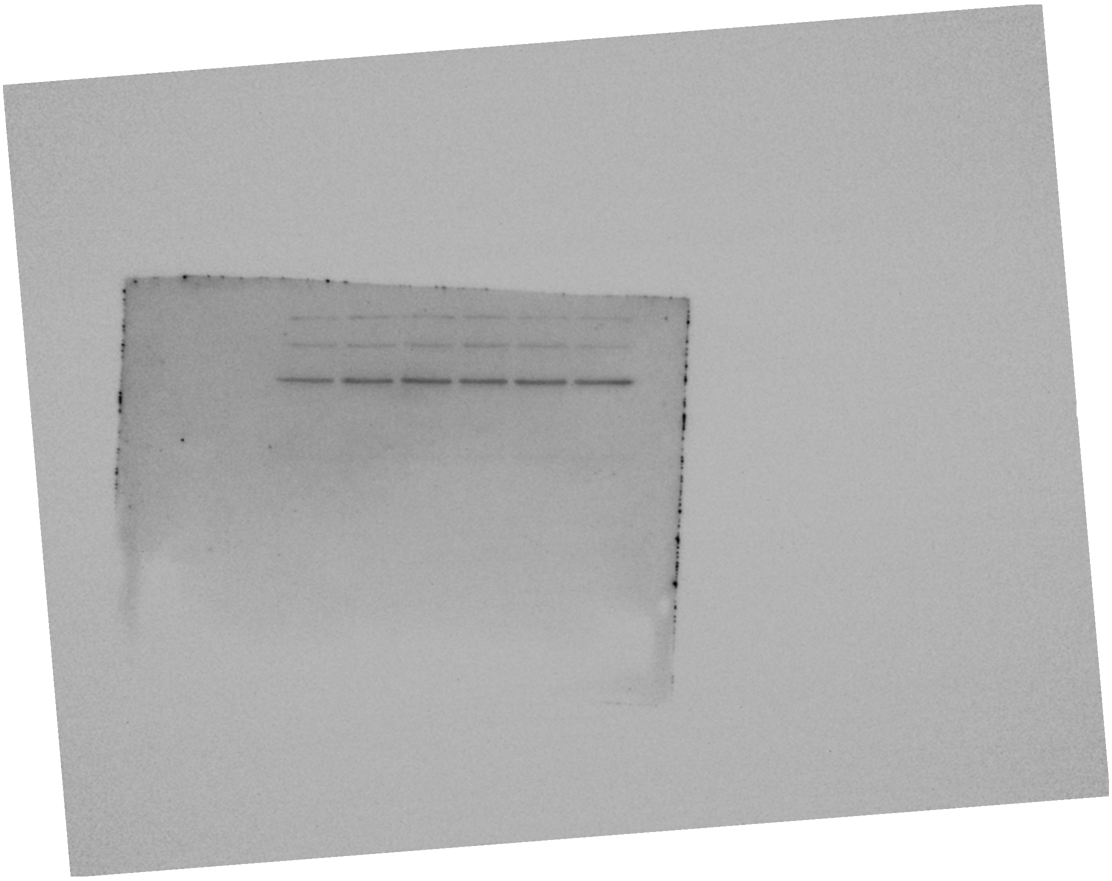

**Ponceau**

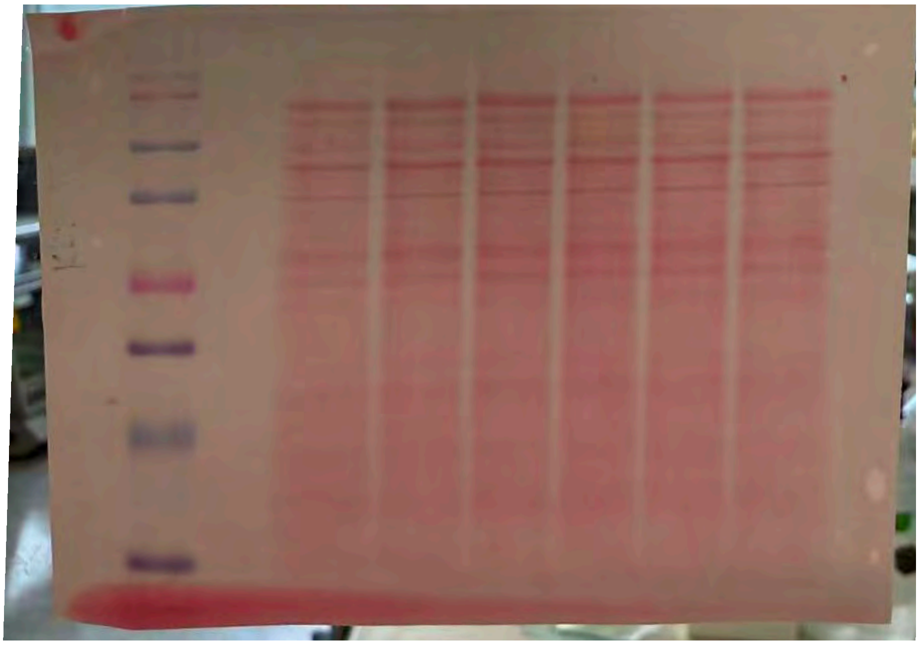

**Fig.1d**

**CatS, CatL, CatC Immunoprecipitation**

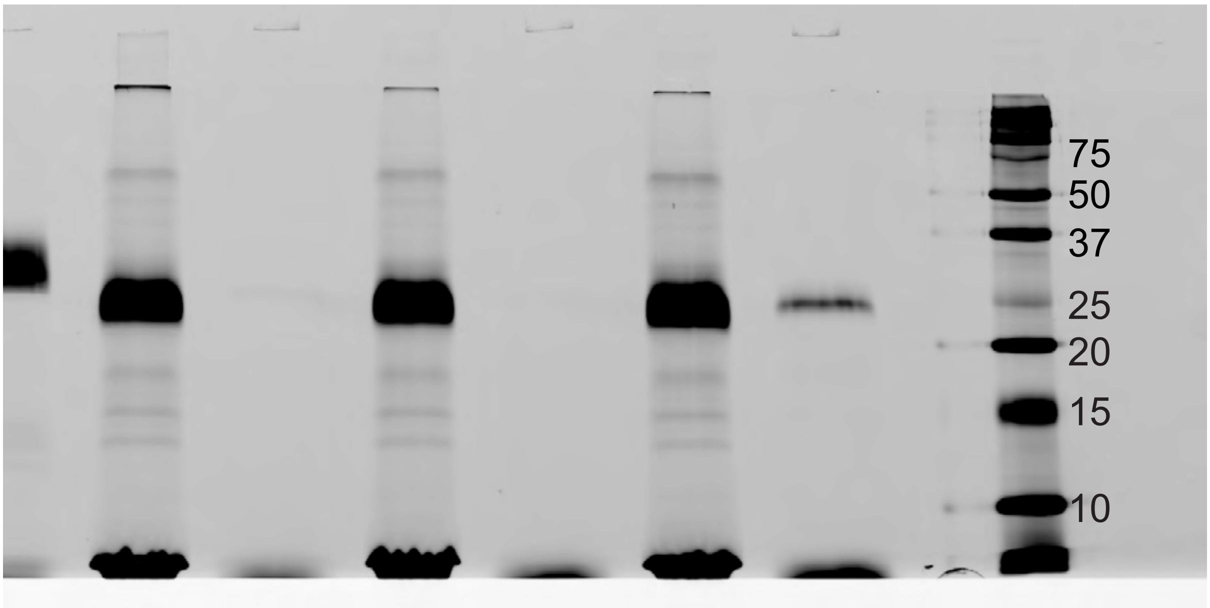

**Fig1e**

**Immunoprecipitation CatC**

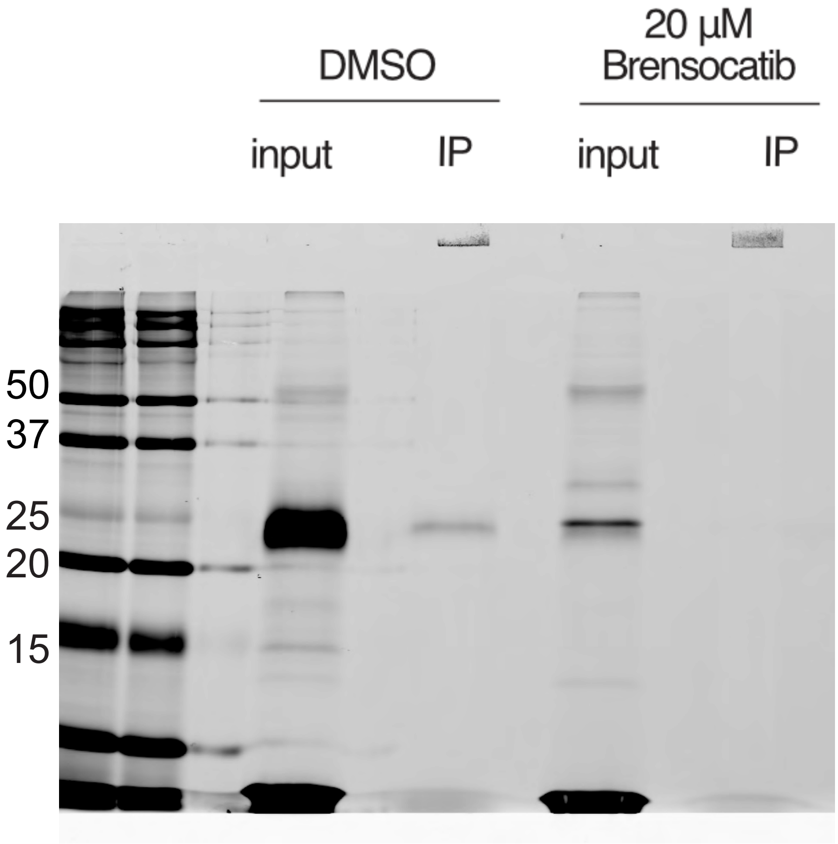

Fig1f

PK105b Labeling

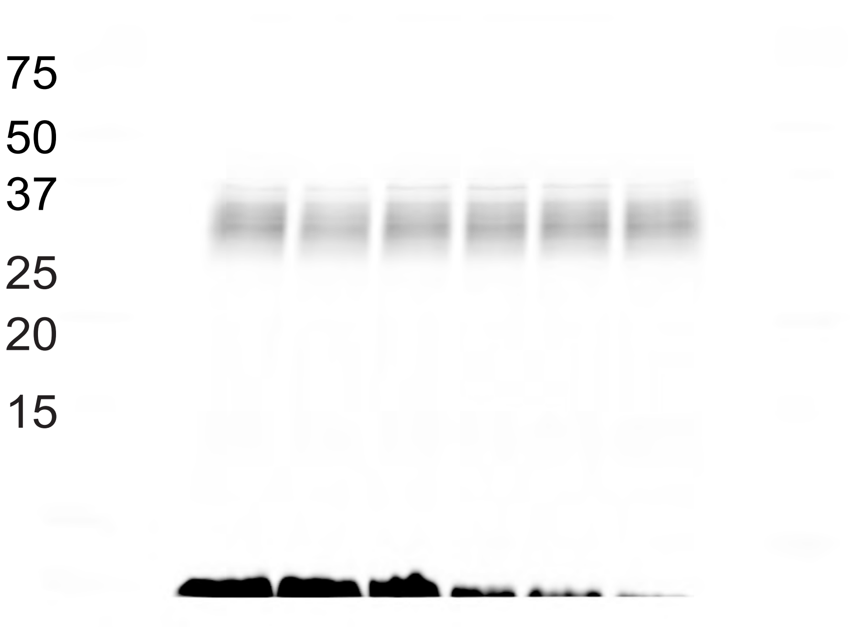


NE Immunoblot

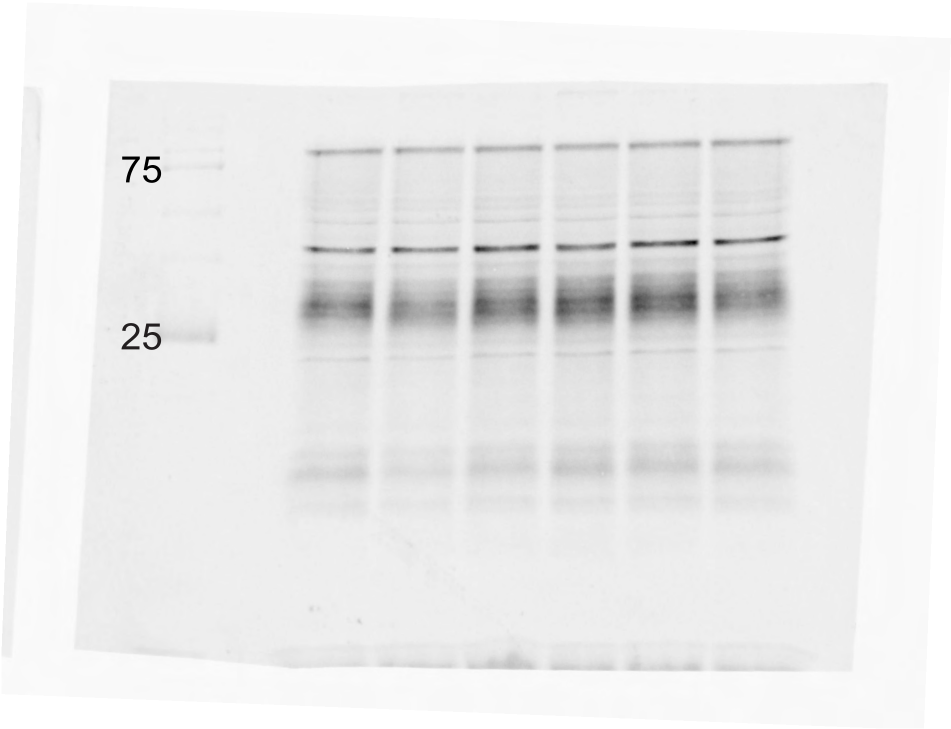

**Actin**

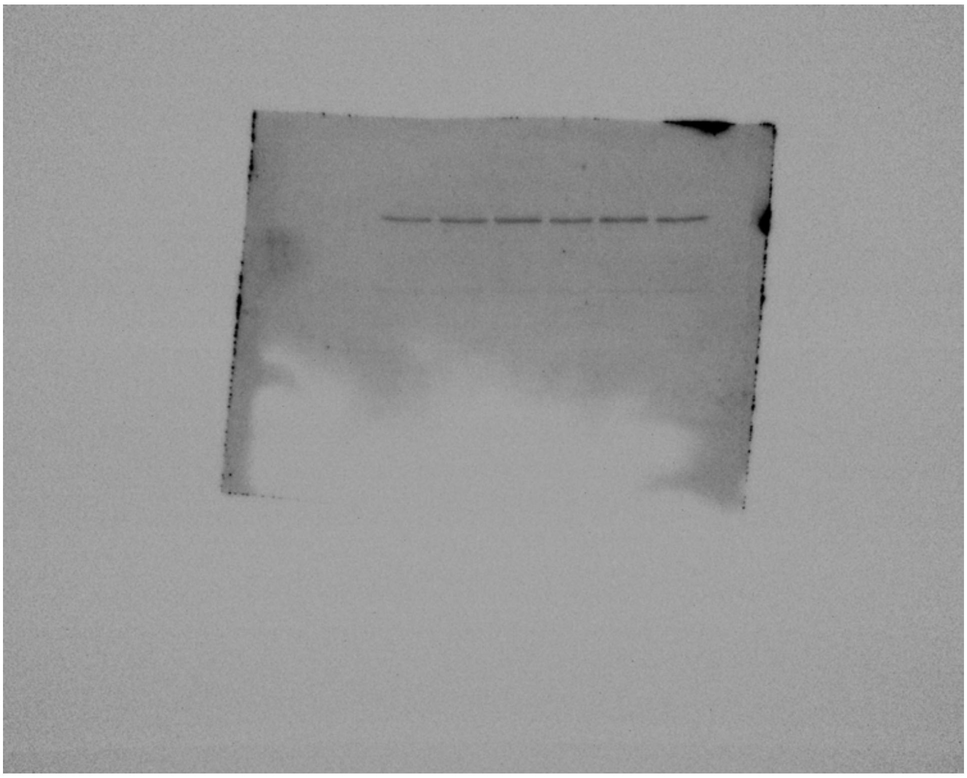

**Ponceau**

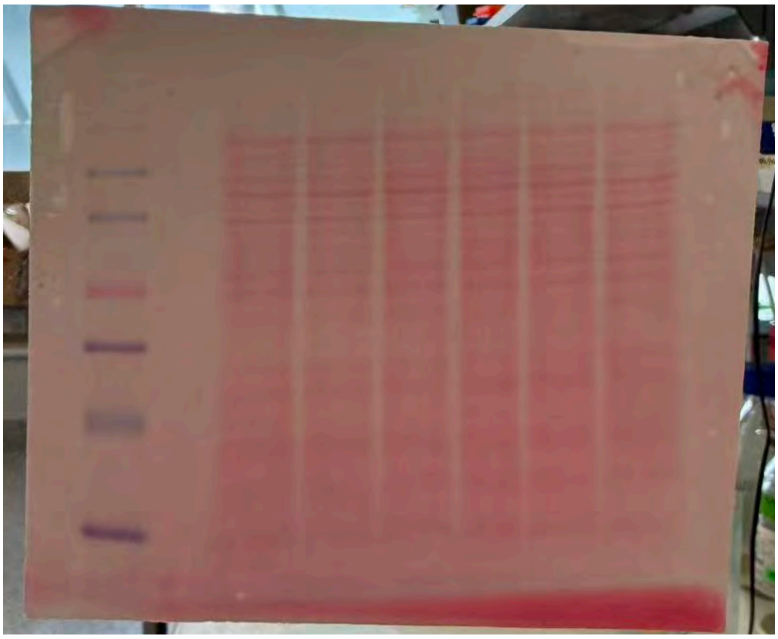

**Fig1i**

**Immunoprecipitation NE**

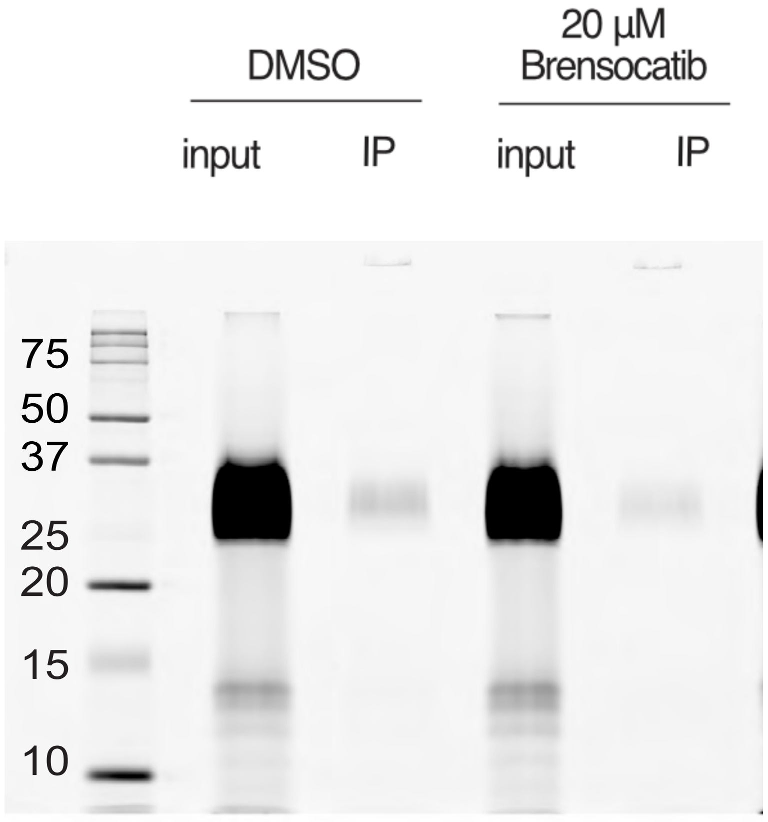

**Fig 1j**

**Immunoprecipitation PR3**

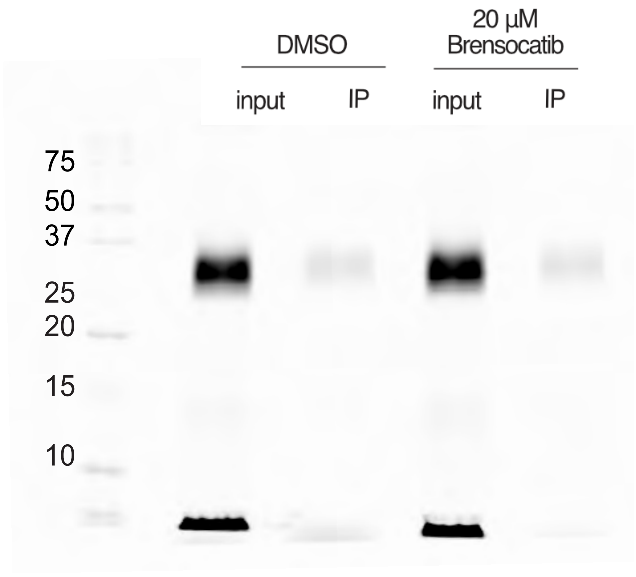

**Figure 2a**

**FY01 Labeling**

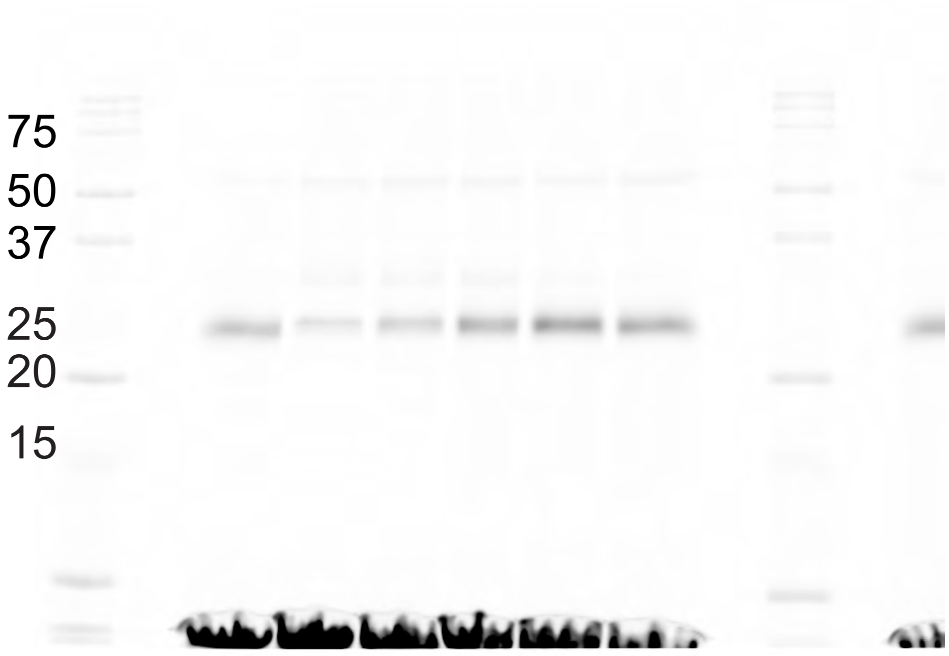


**CatC Immunoblot**

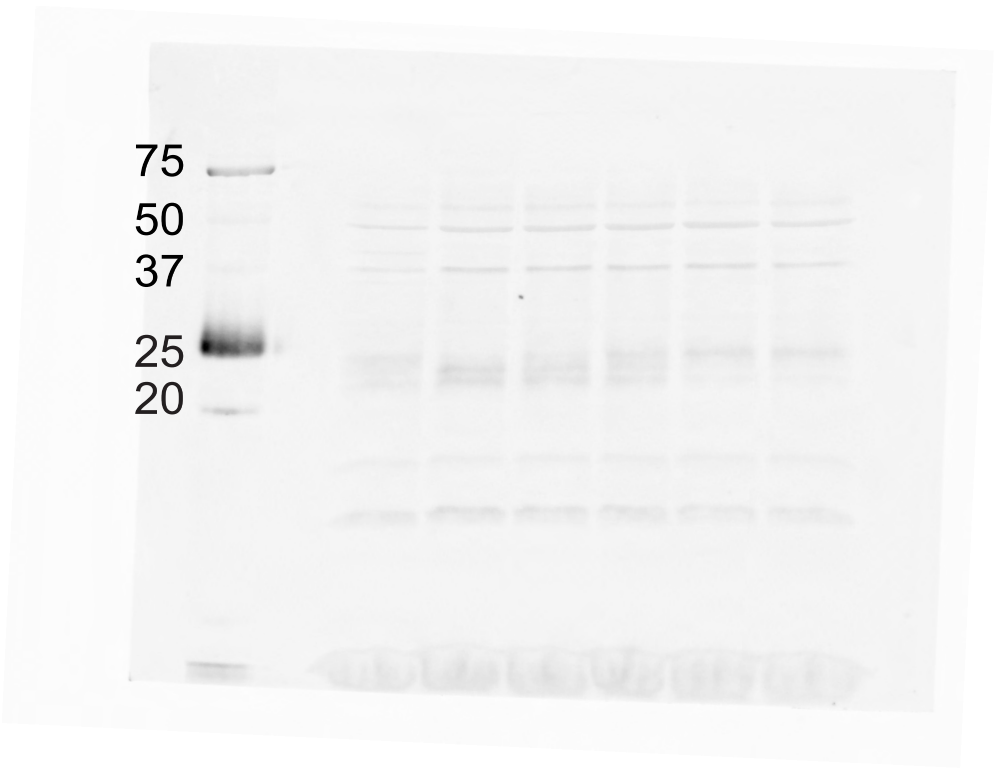

**Actin**

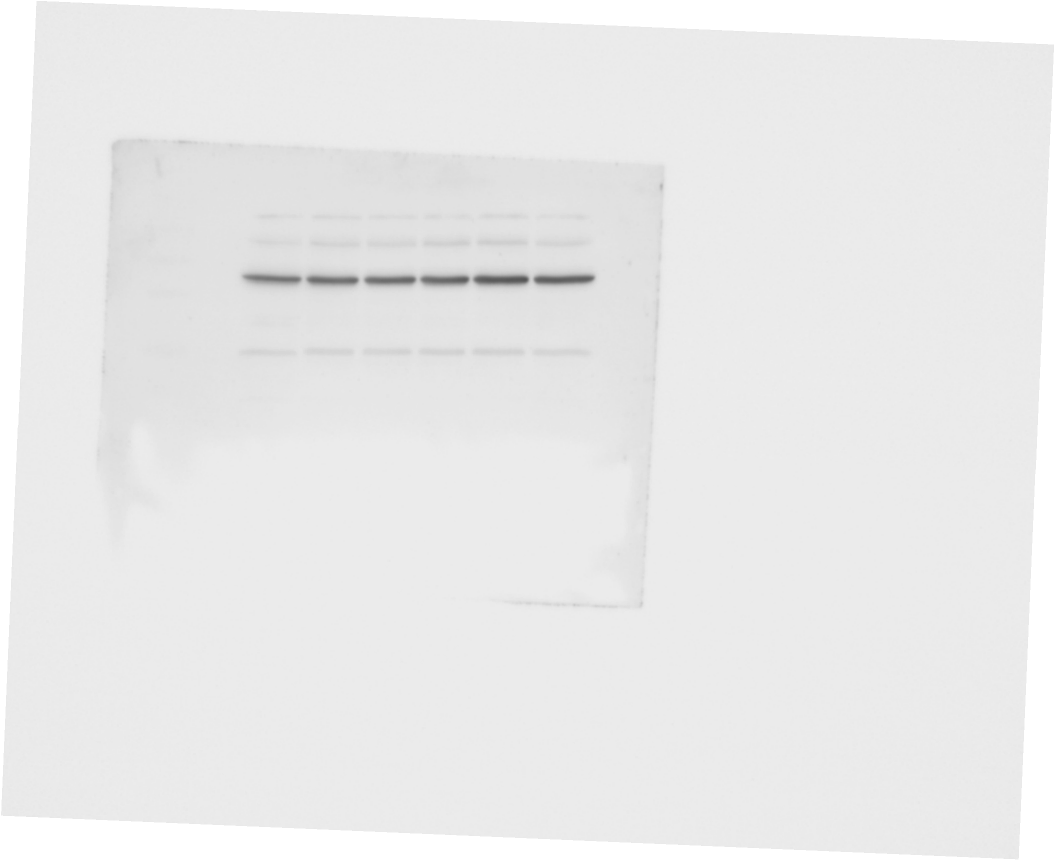

**Ponceau**

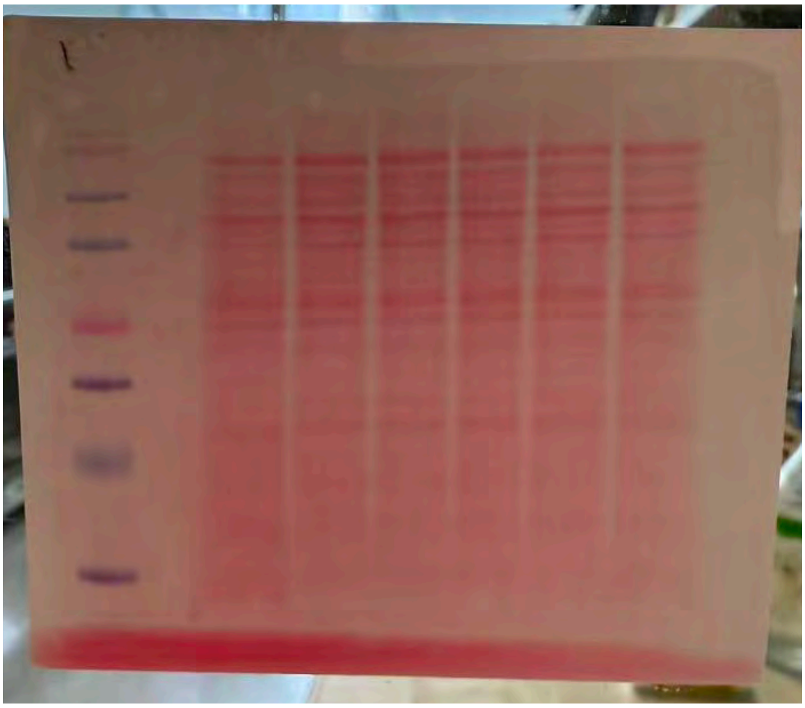

**Figure 2d**

**Immunoprecipitation CatC**

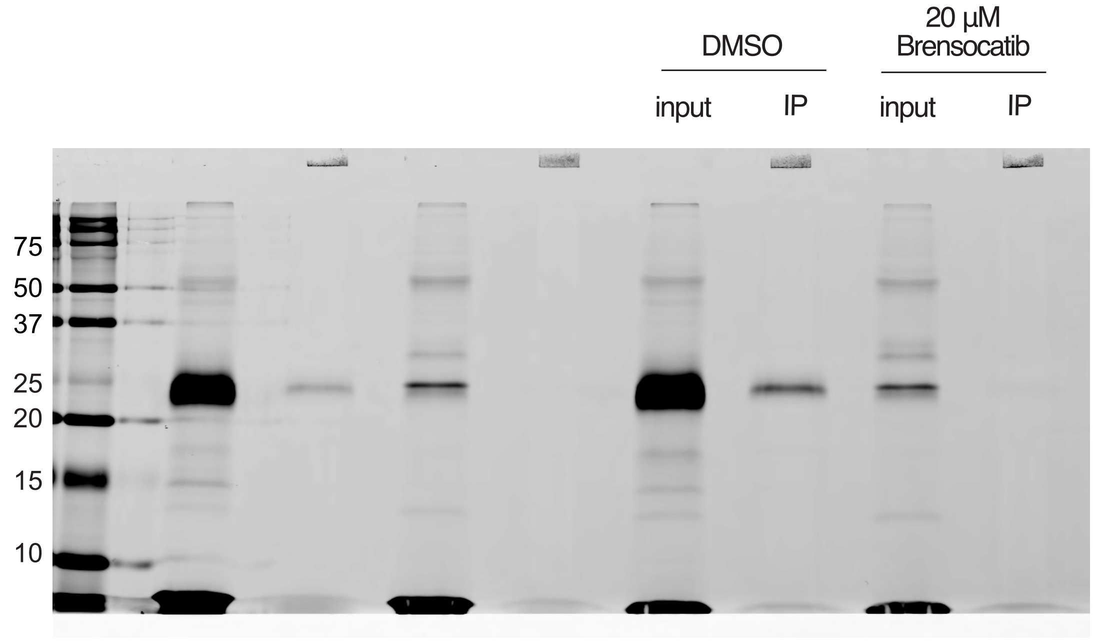

**Fig2e**

**PK105b labeling**

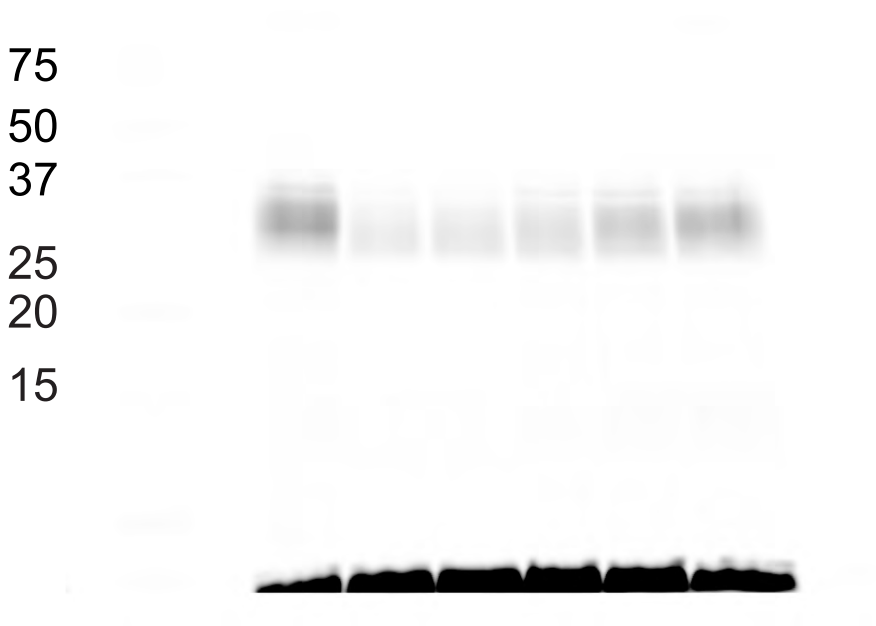


**NE Immunoblot**

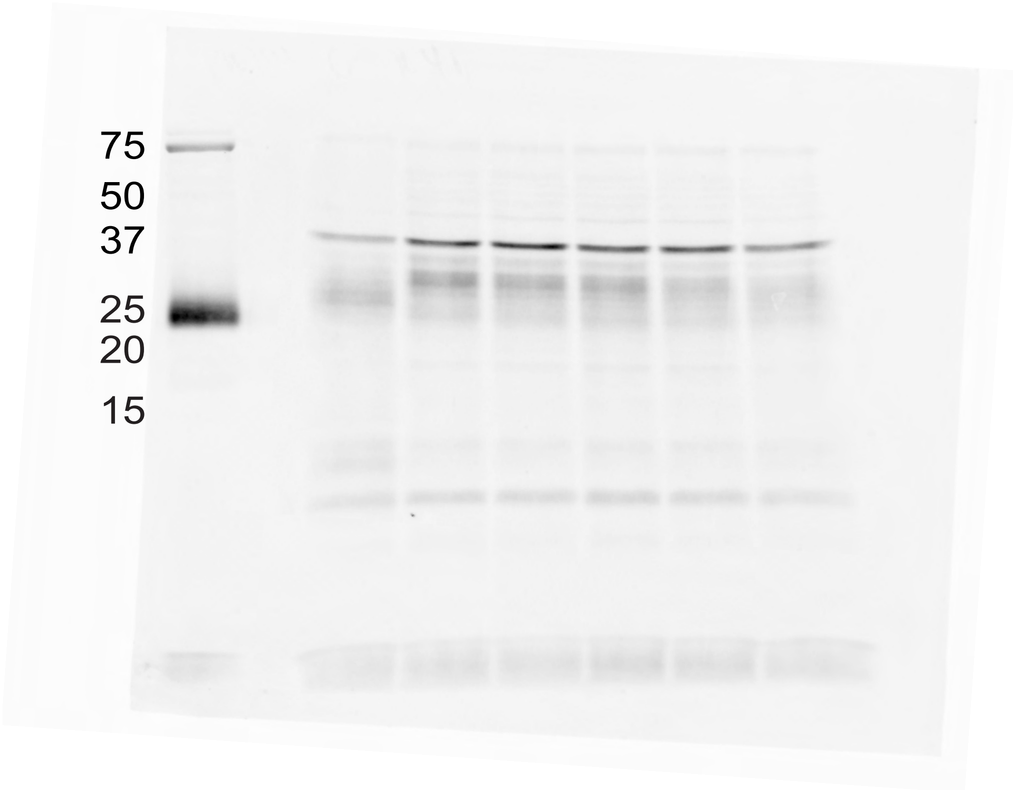

**Actin**

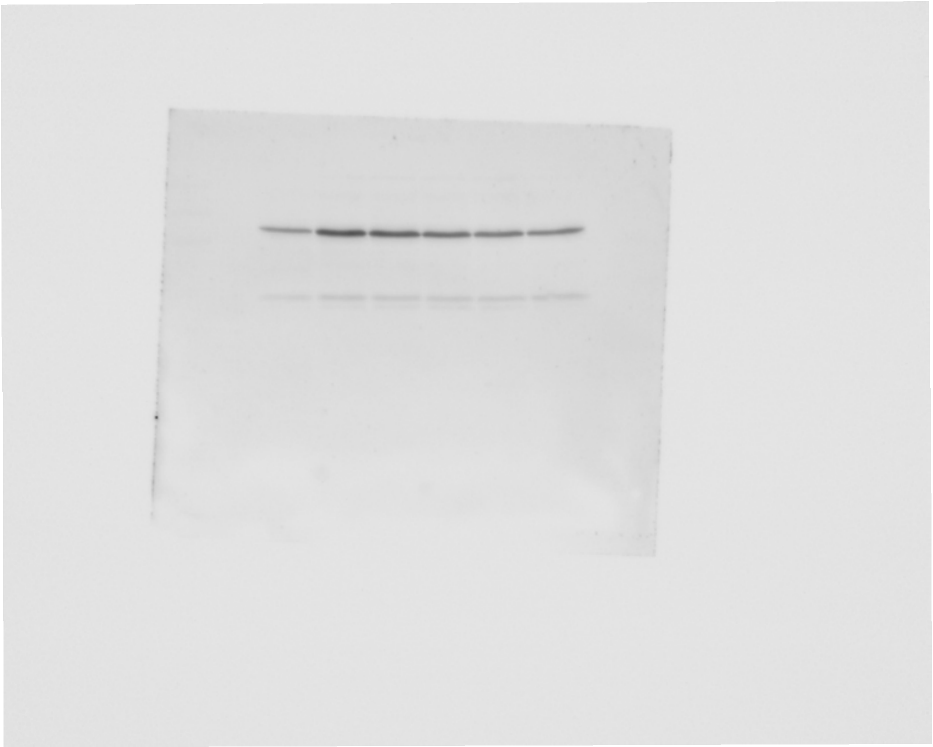

Ponceau

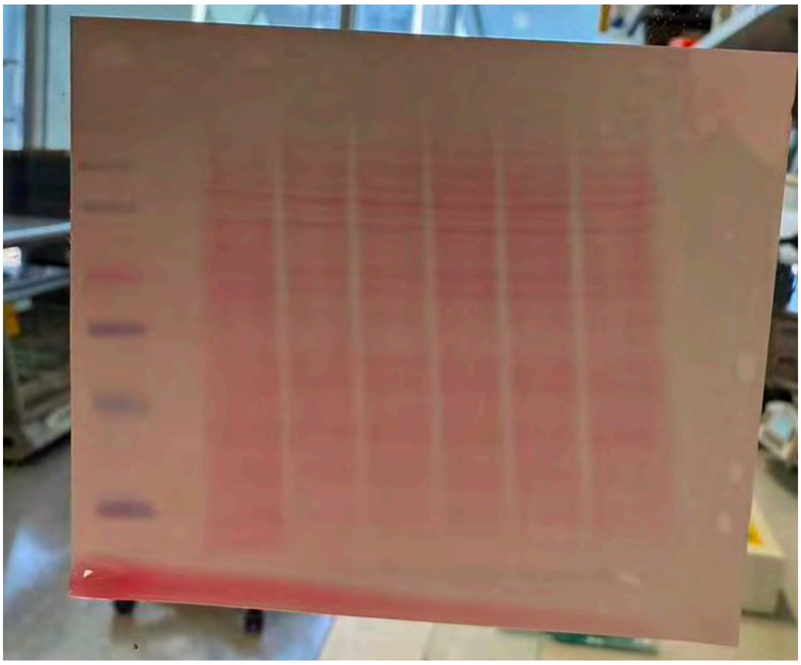

Fig2h

Immunoprecipitation NE

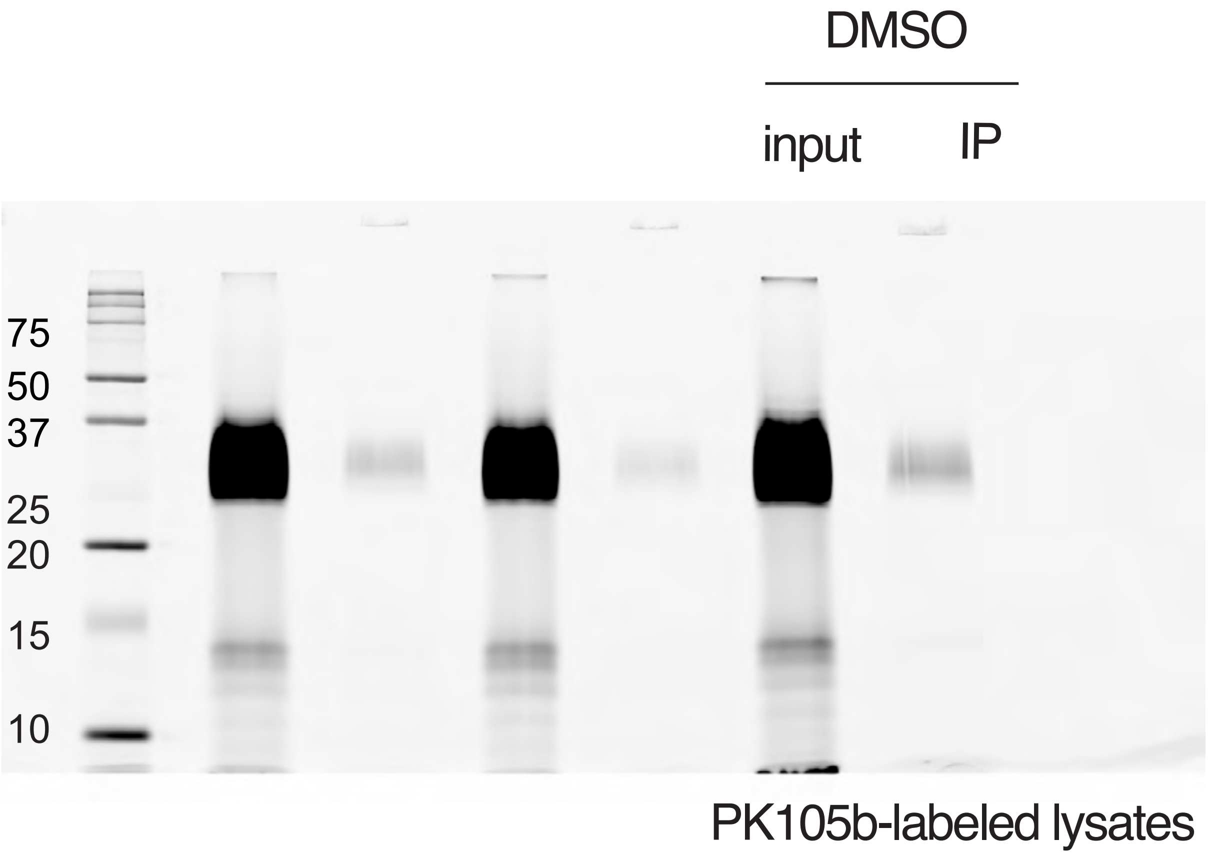

**Immunoprecipitation NE**

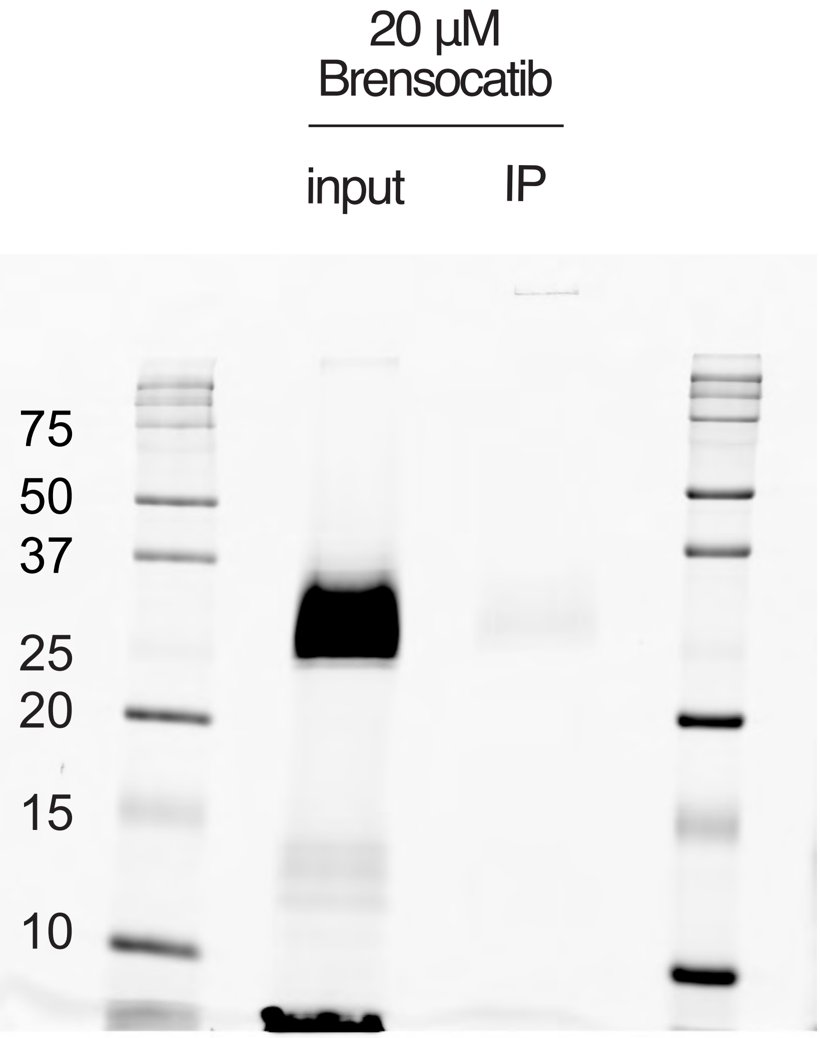

PK105b-labeled lysates

**Fig2i**

**Immunoprecipitation PR3**

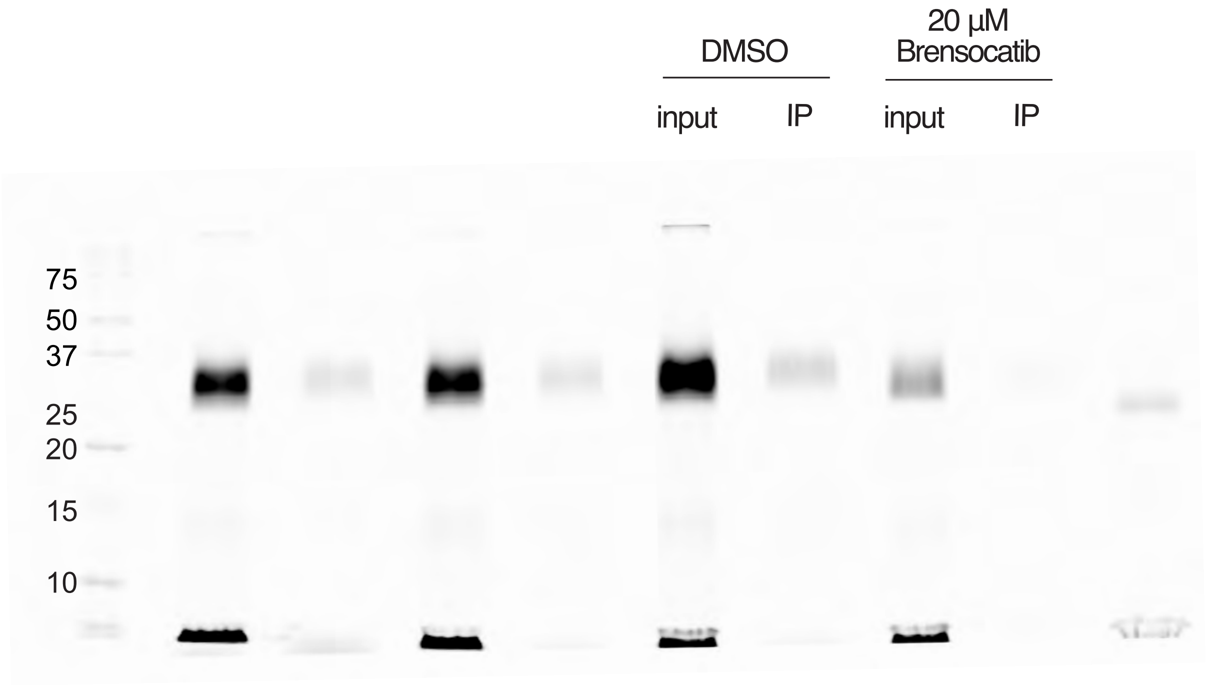

Fig3a

**FY01 labeling**

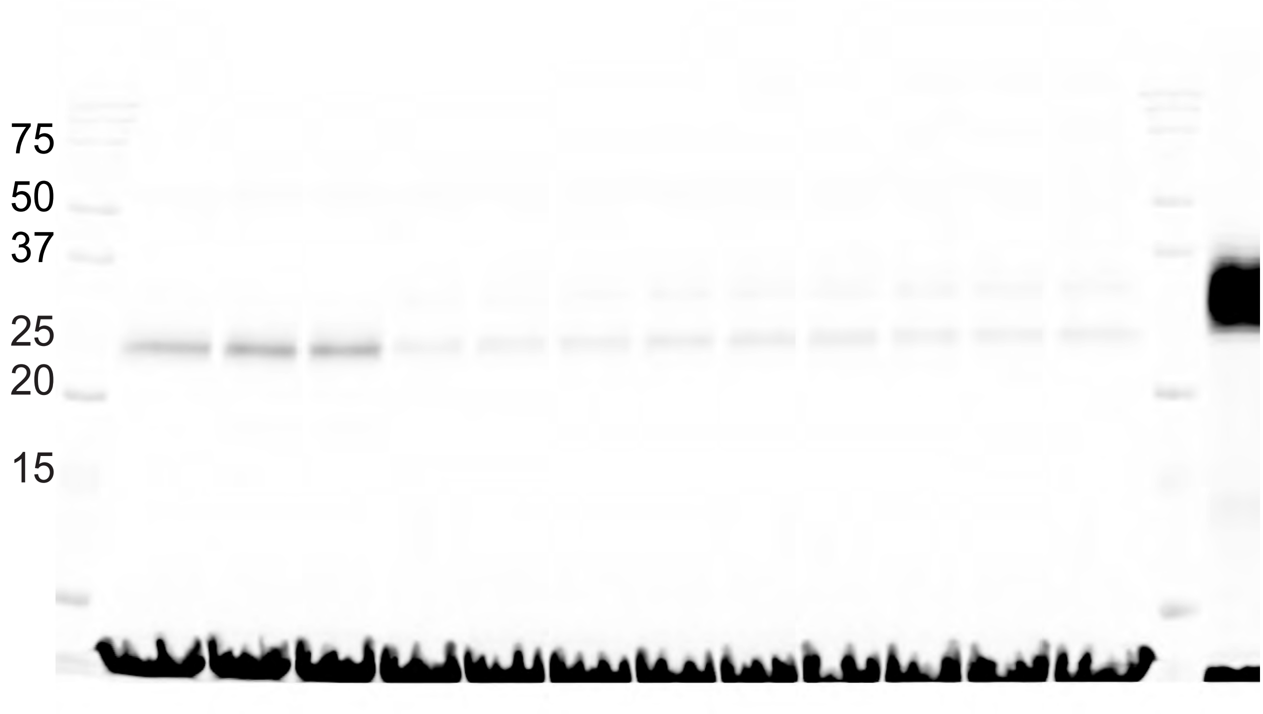


**CatC Immunoblot**

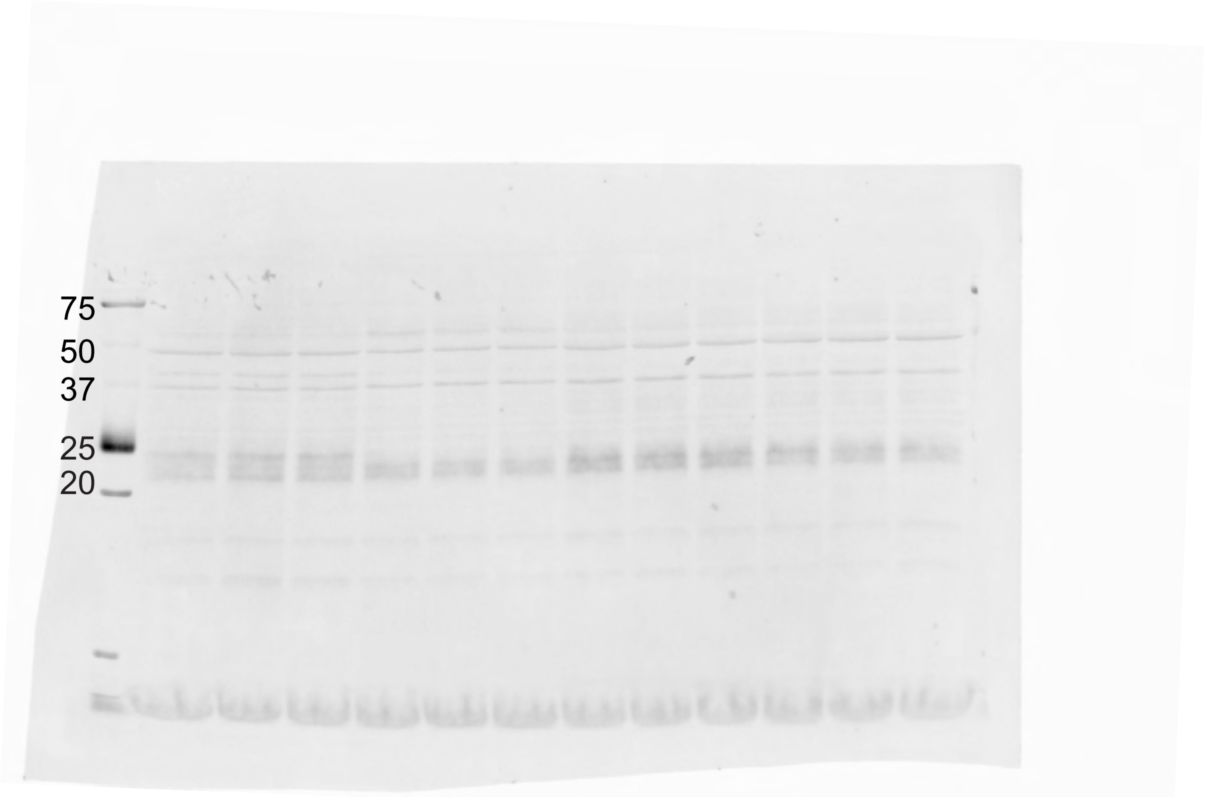

**Actin**

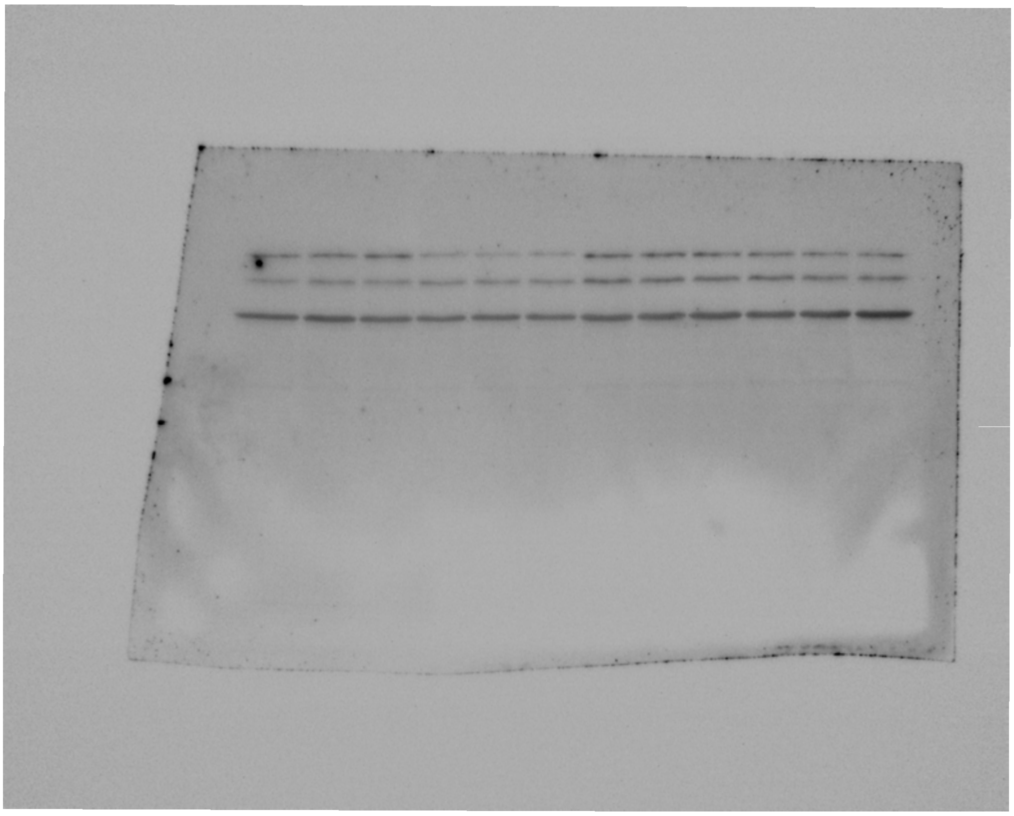

**Ponceau**

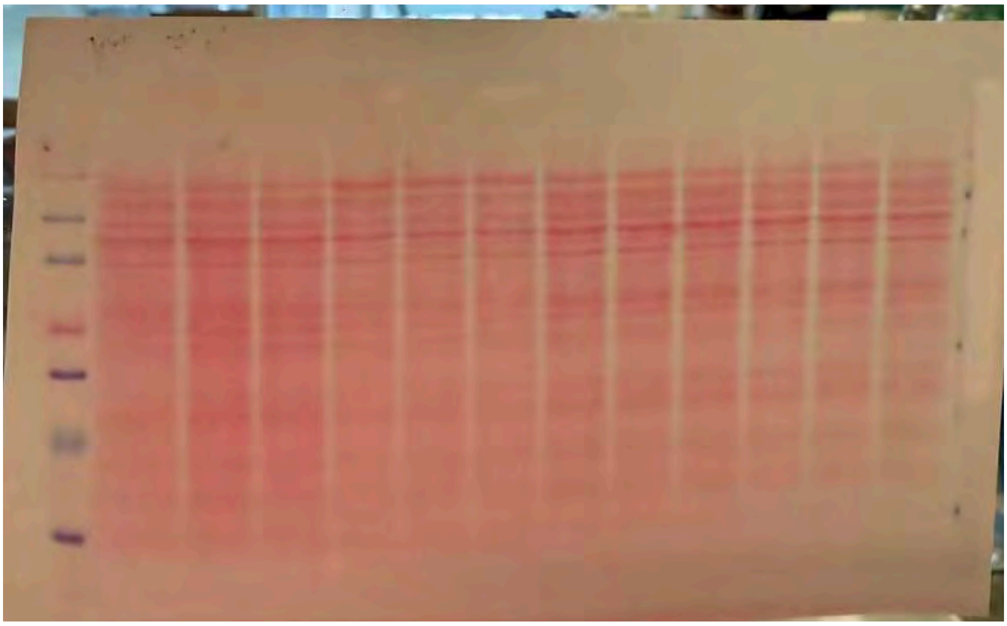

**Fig. 3d**

**PK105b labeling**

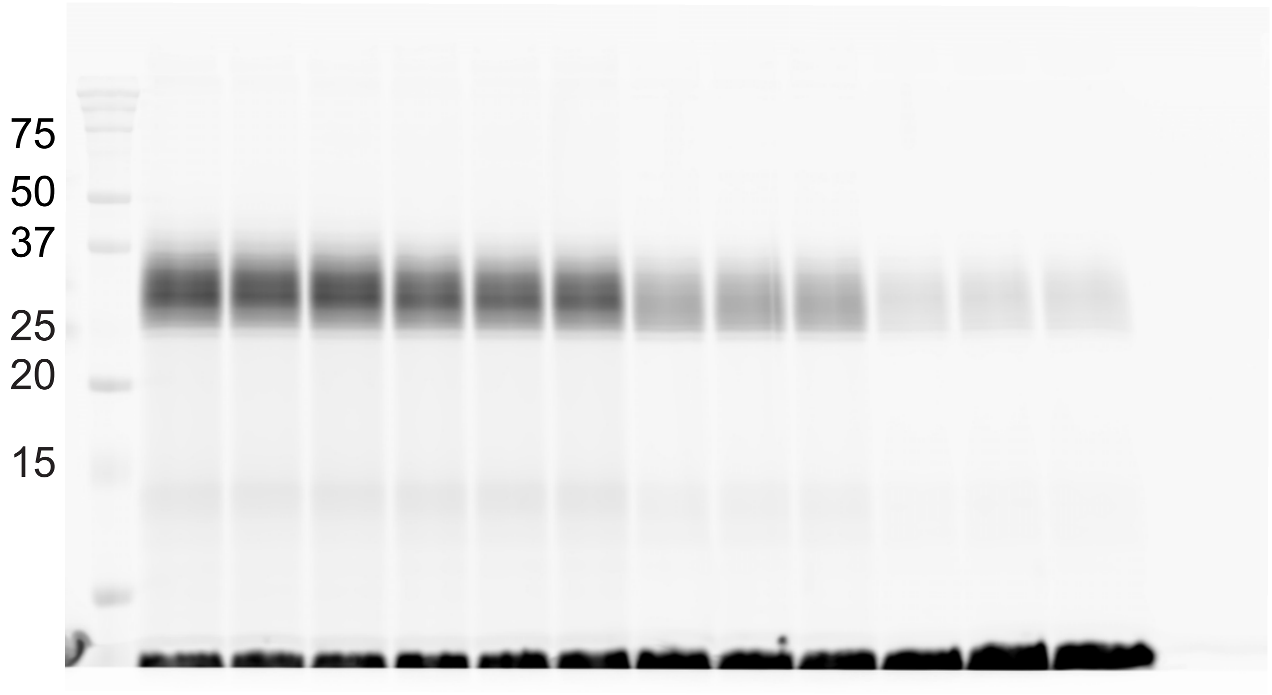


**NE immunoblot**

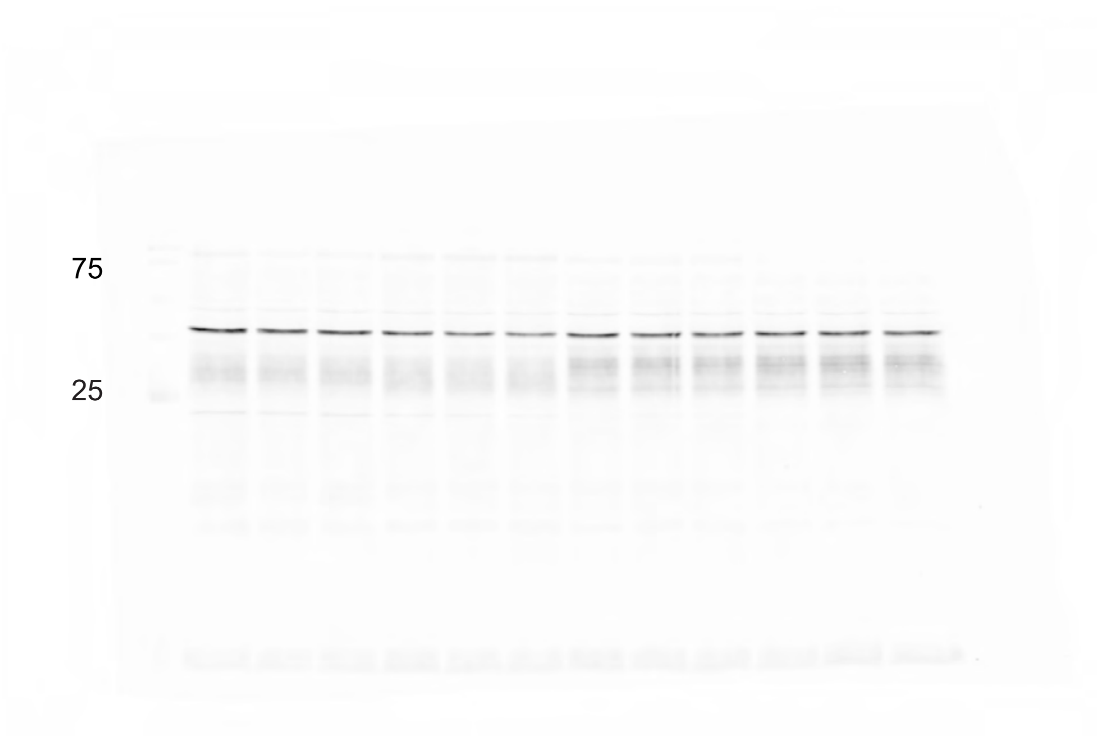

**Actin**

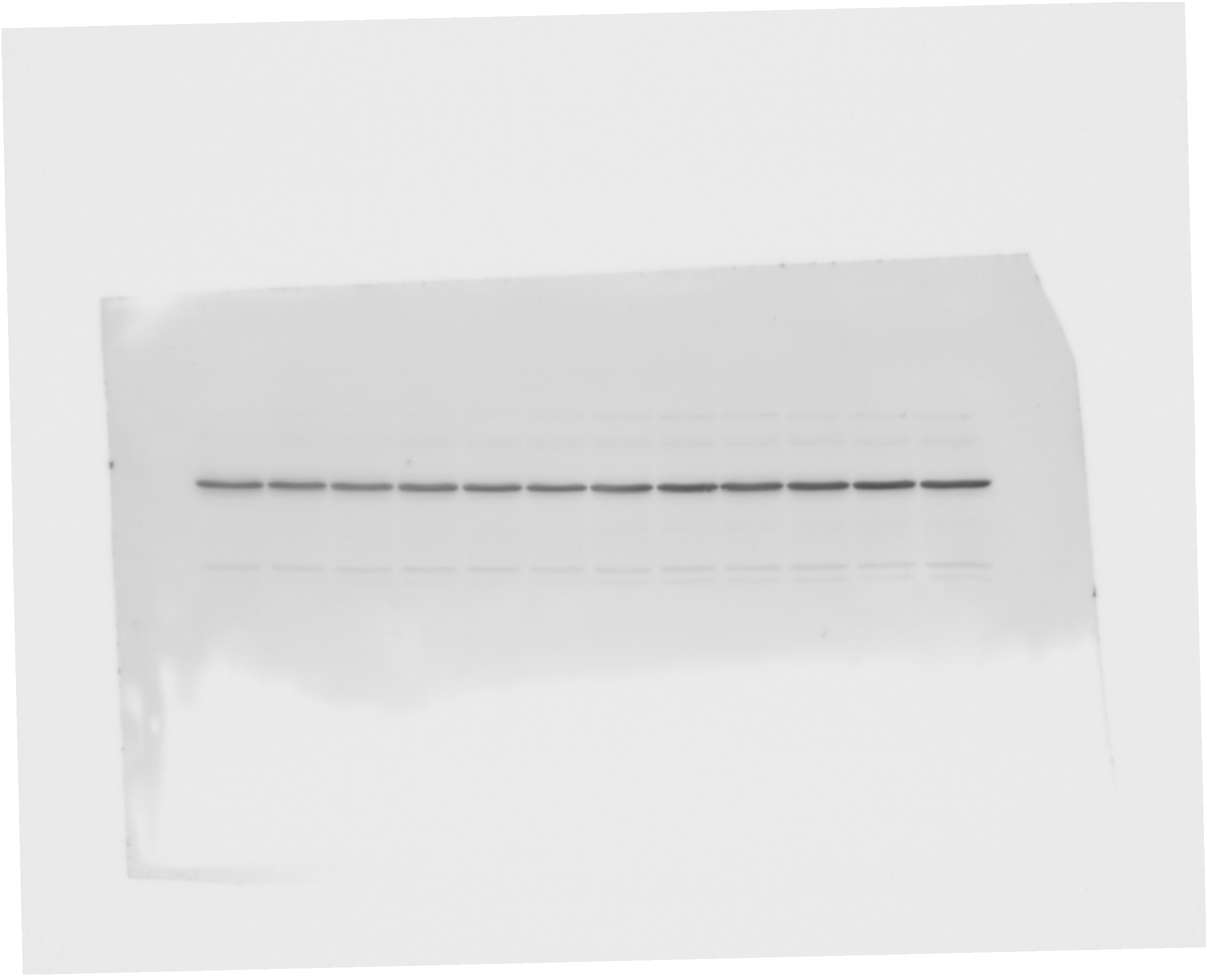

**Ponceau**

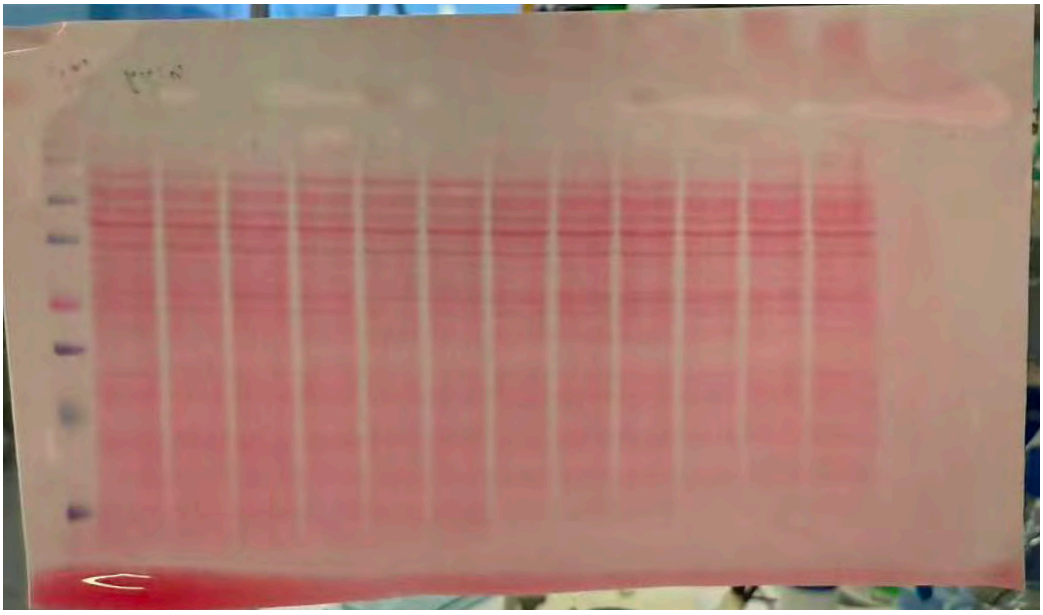

Supplementary Fig2a

PR3 Immunoblot

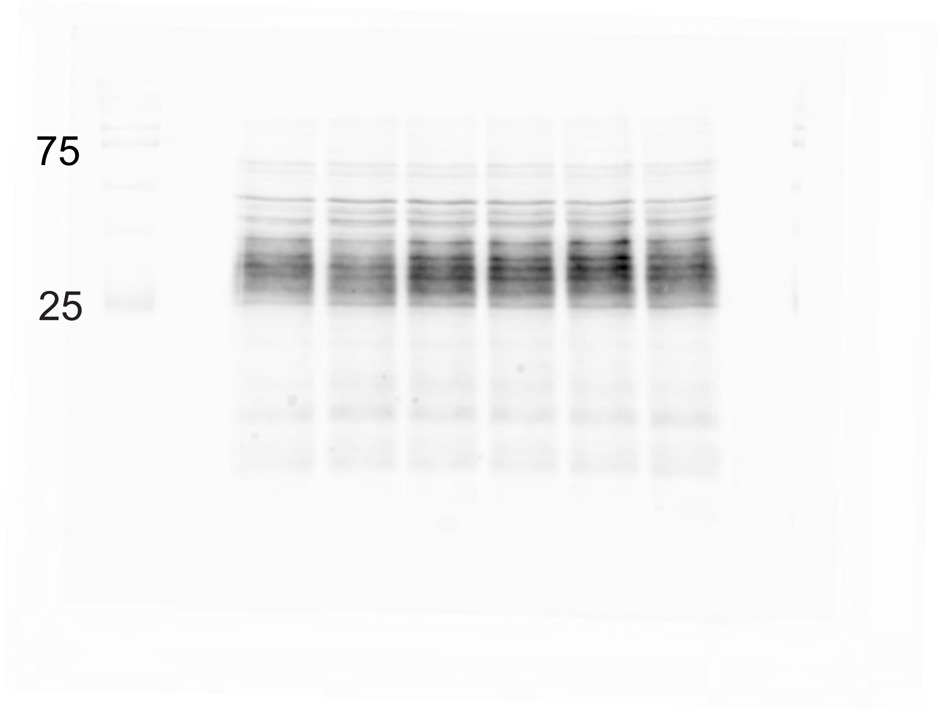


Actin

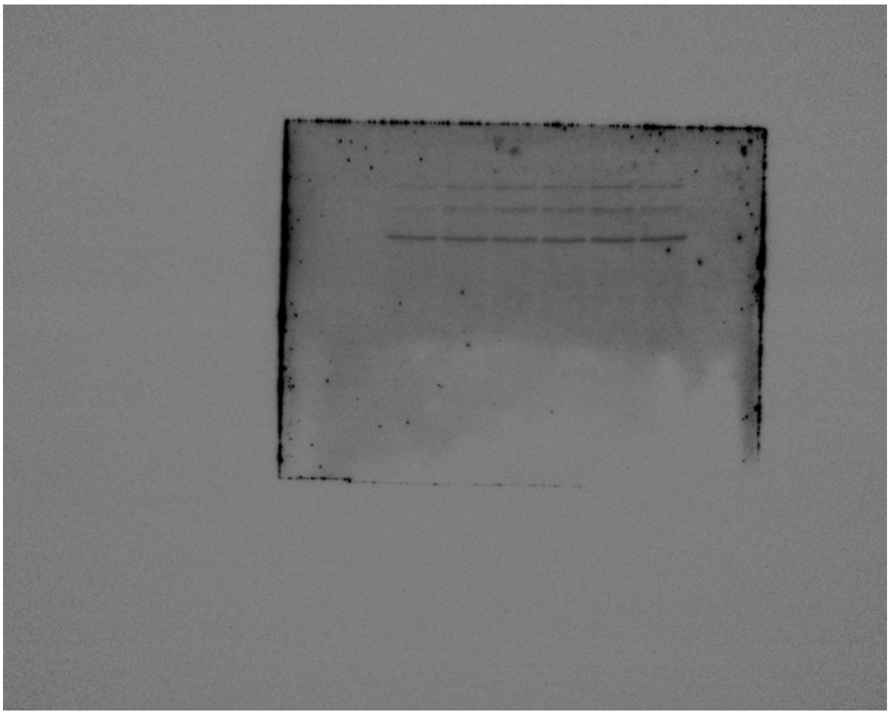

Ponceau

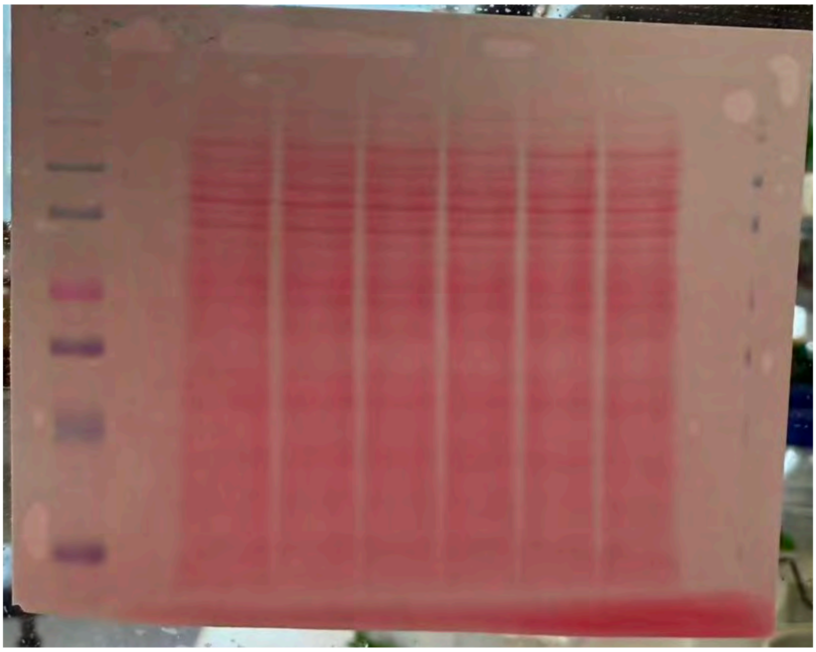

Supplementary Fig 2b

PK105b labeling

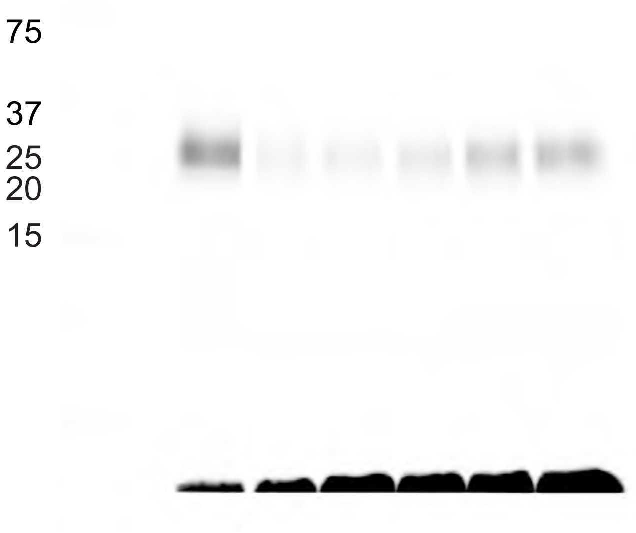


Immunoblot PR3

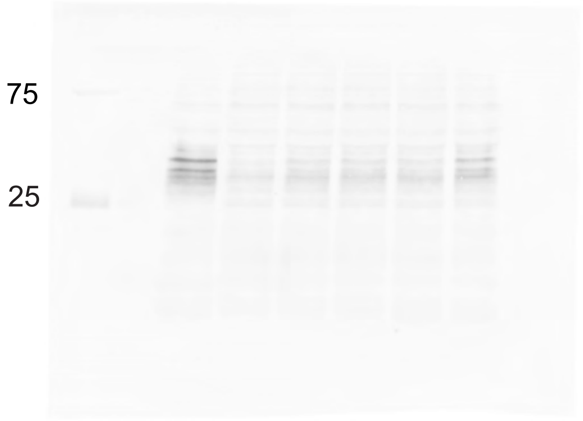

Actin

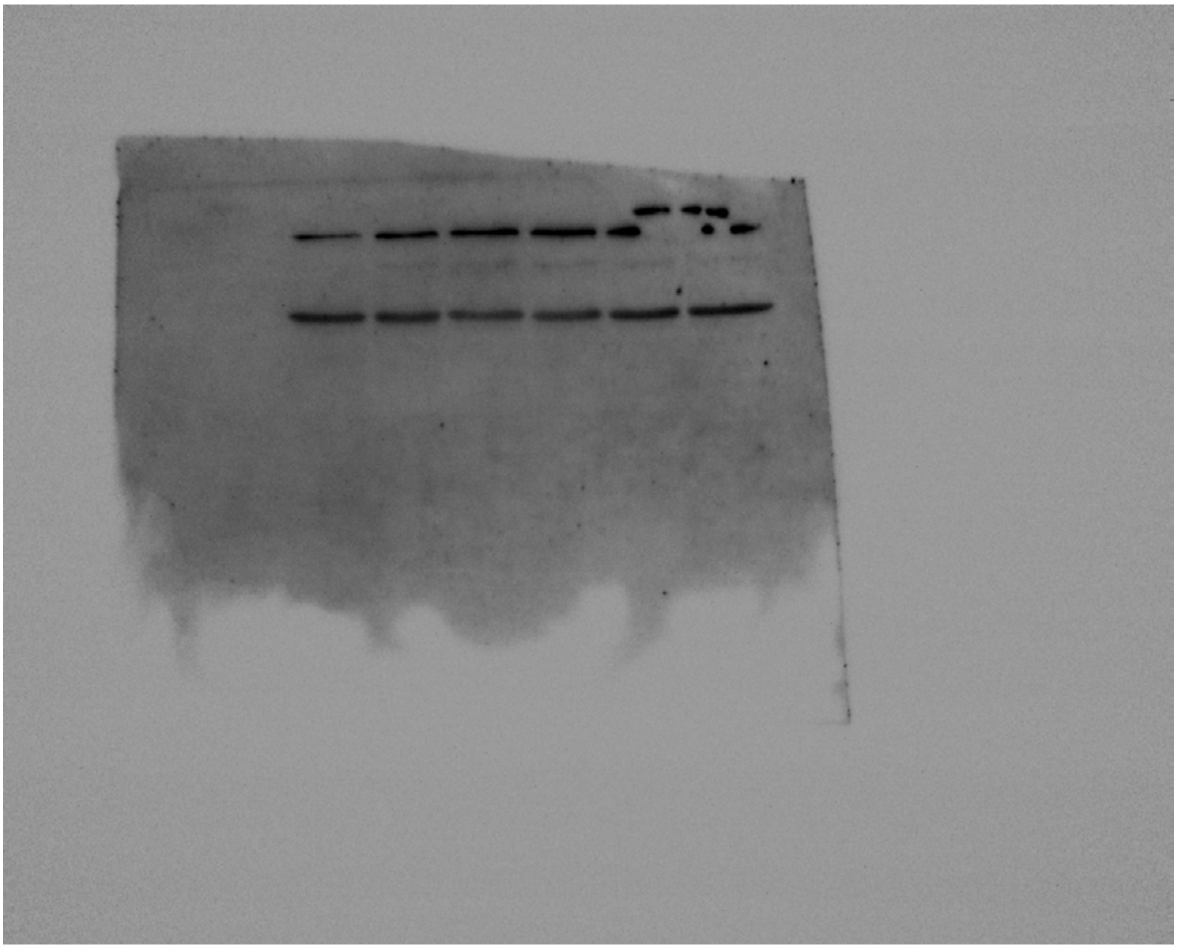

Ponceau

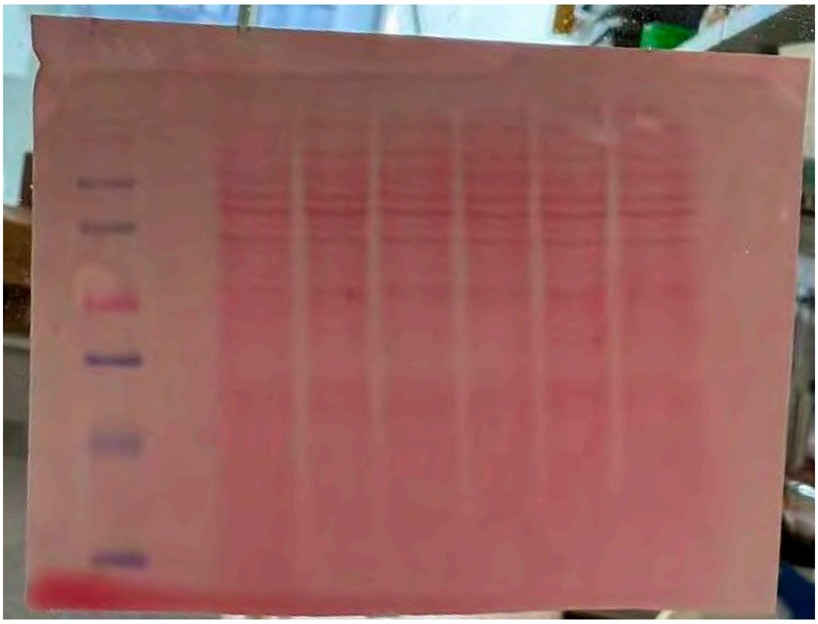

Supplementary Fig2c

PK105b labeling

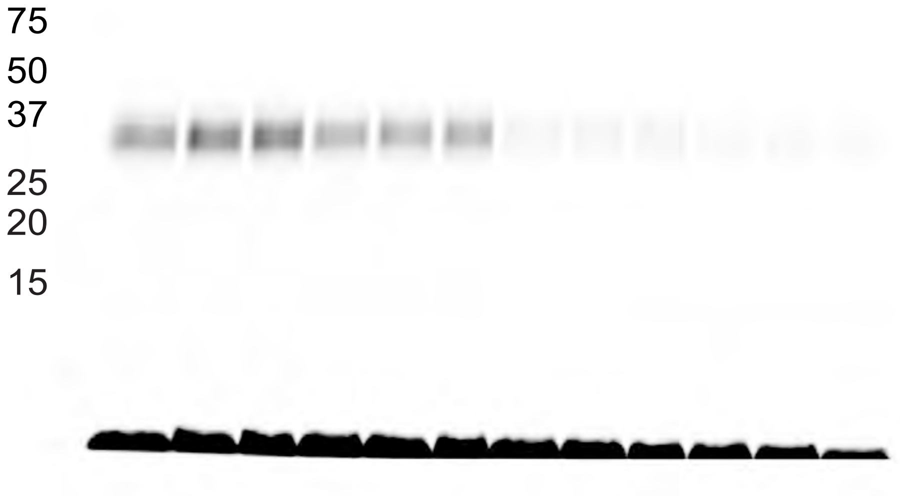


**PR3 Immunoblot**

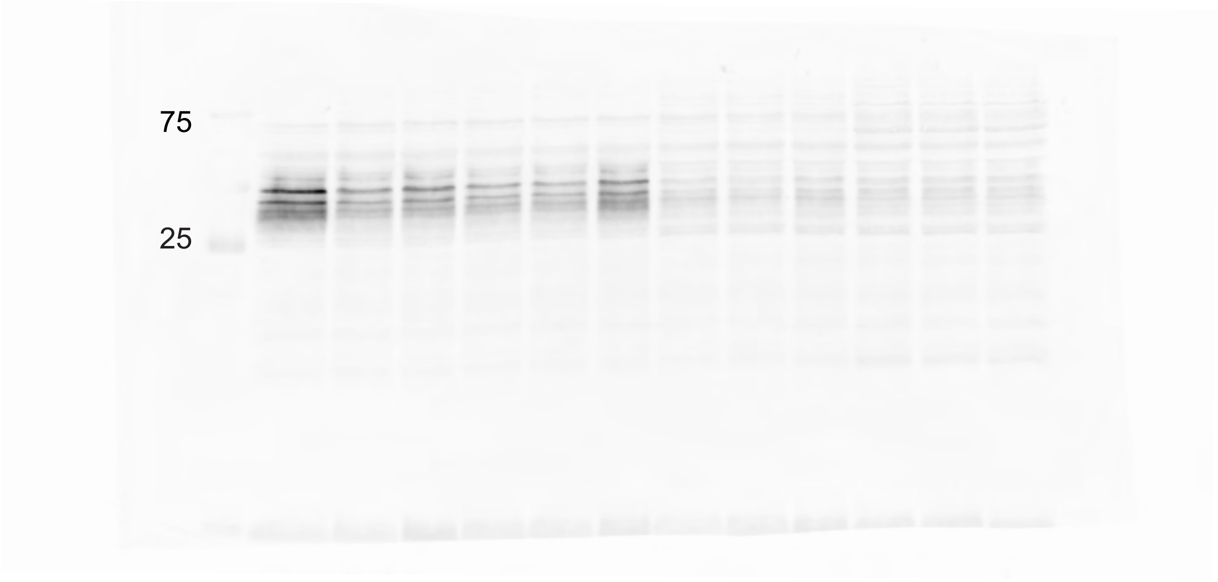

**Actin**

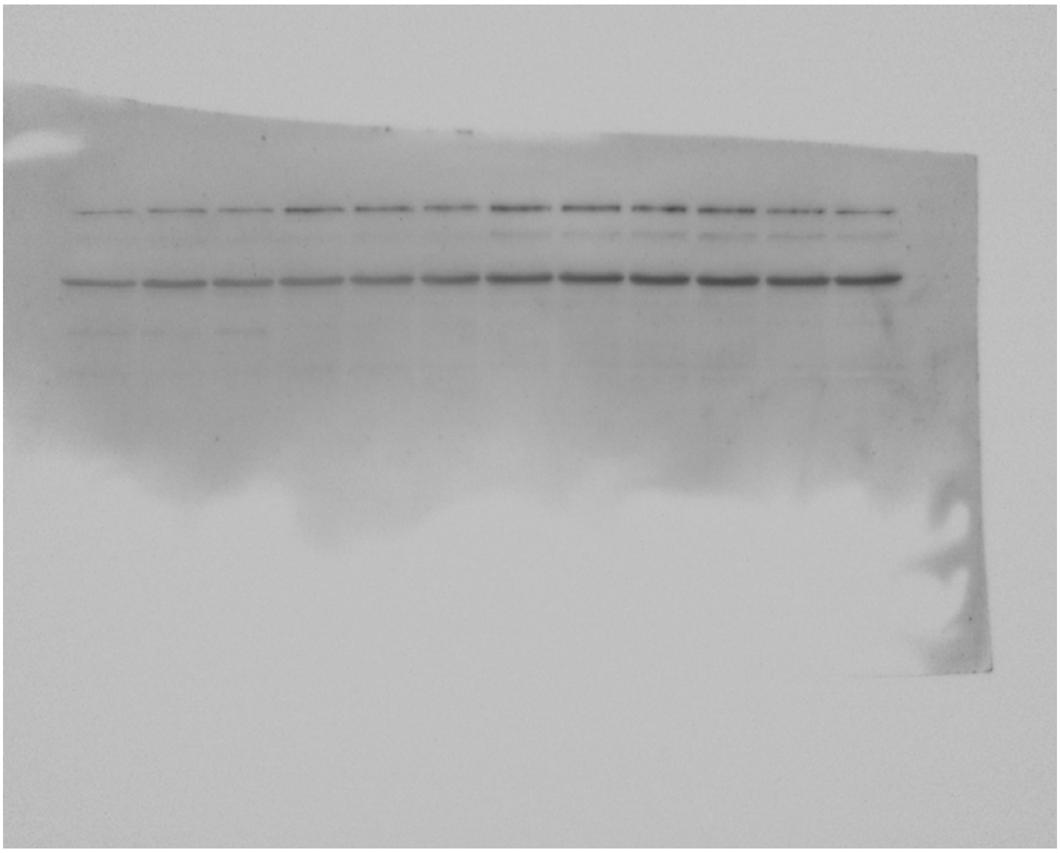

**Ponceau**

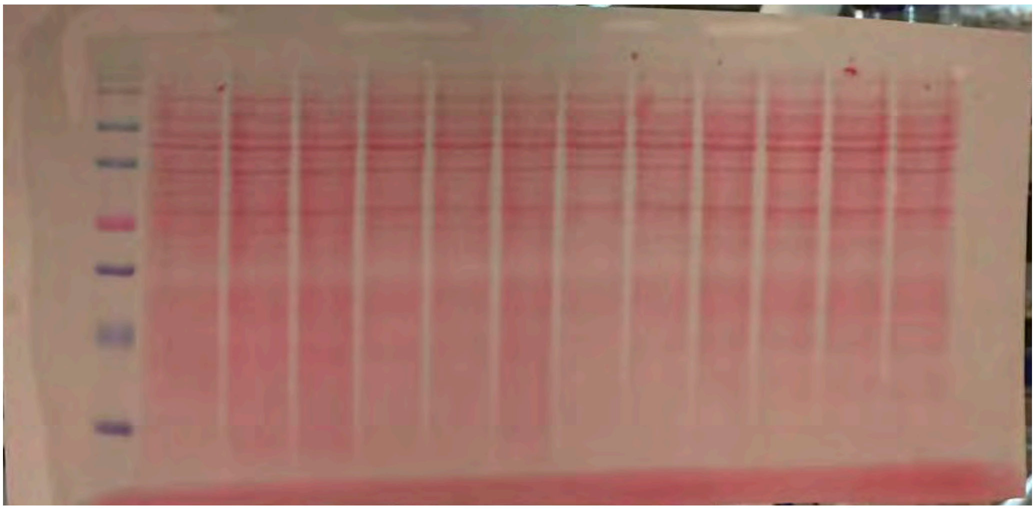

Supplementary Fig 4a

**BMV109 labeling**

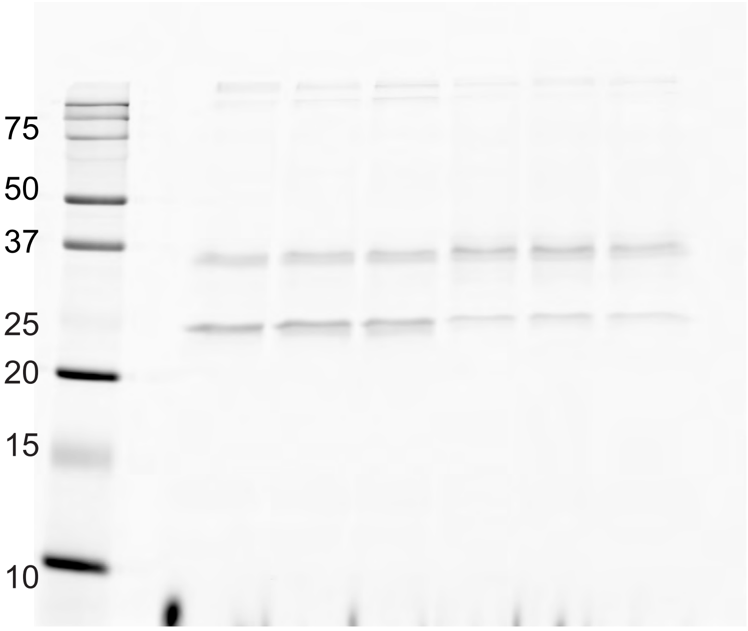


**Immunoblot CatX**

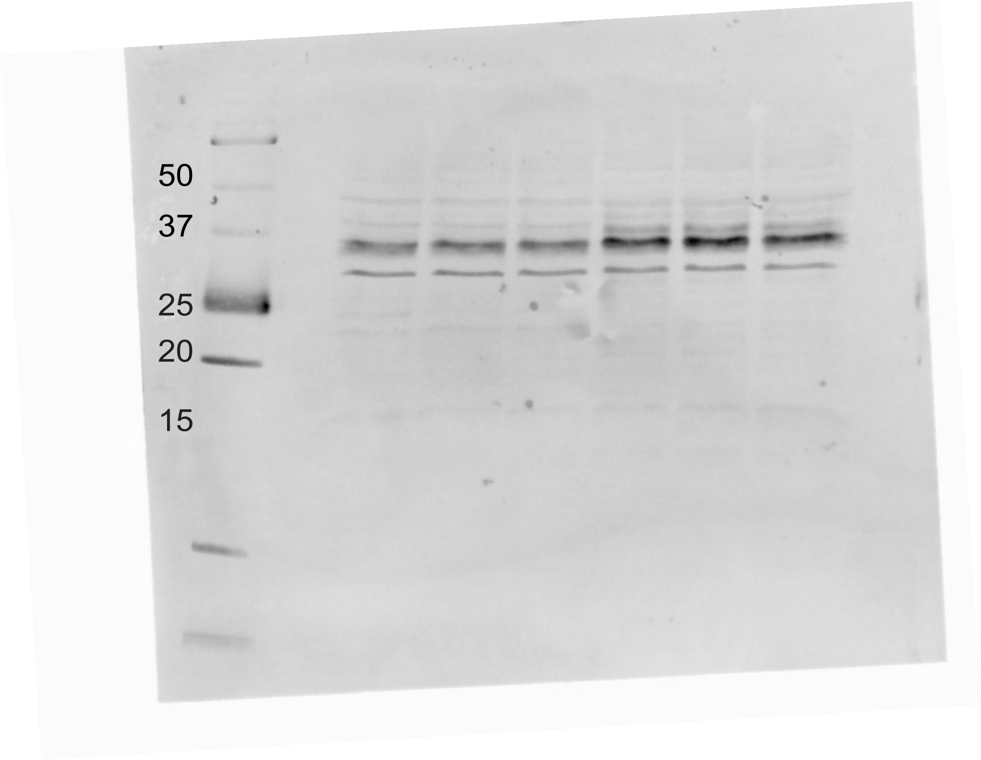

**Actin**

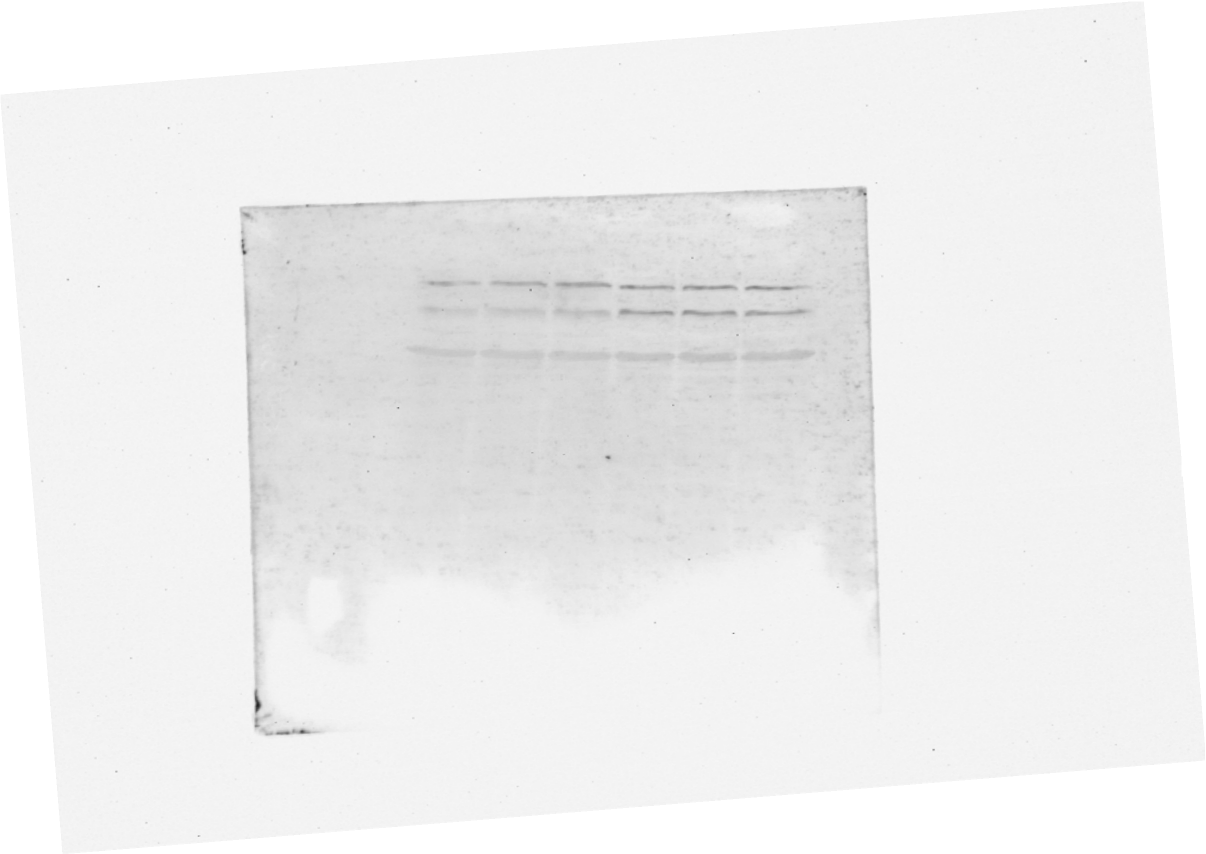

**Ponceau**

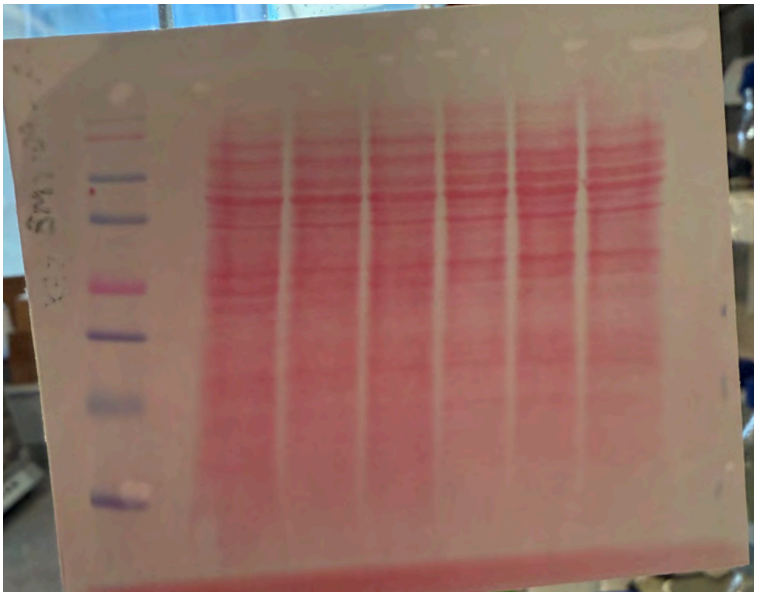

**Supplementary Fig 4e**

**BMV109 labeling**

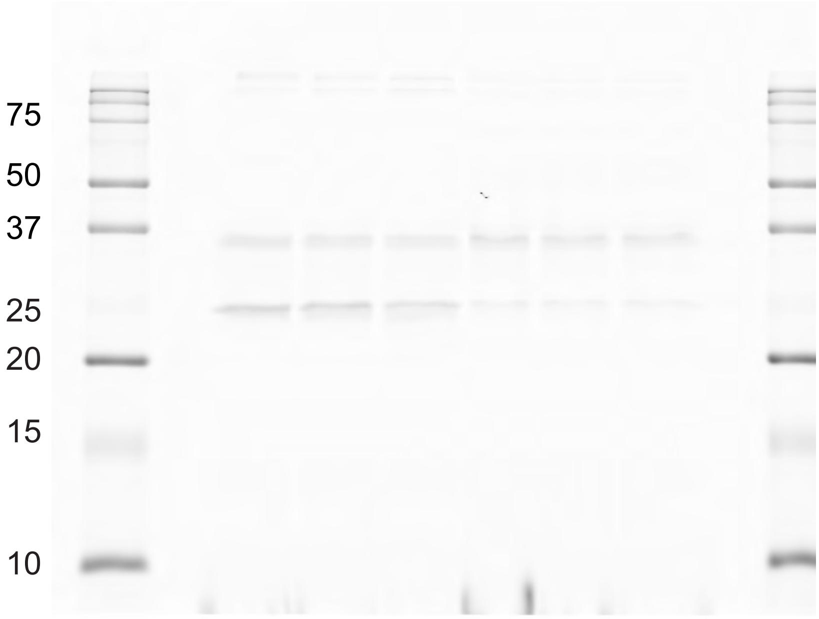


**CatB Immunoblot**

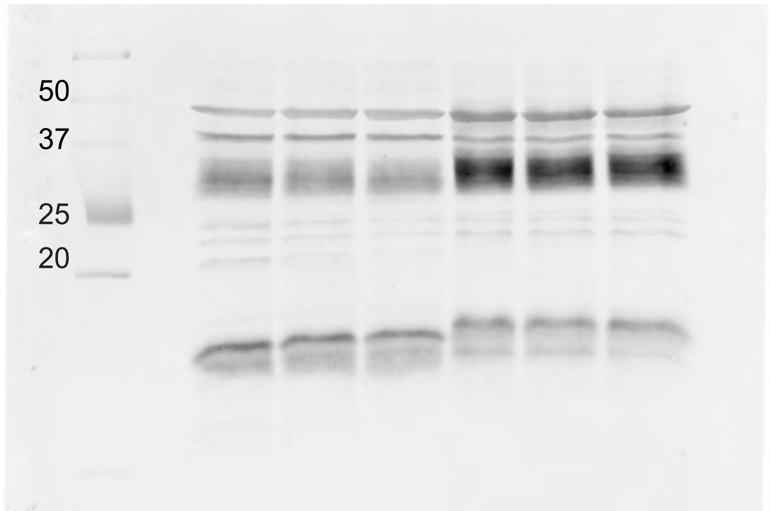

**Actin**

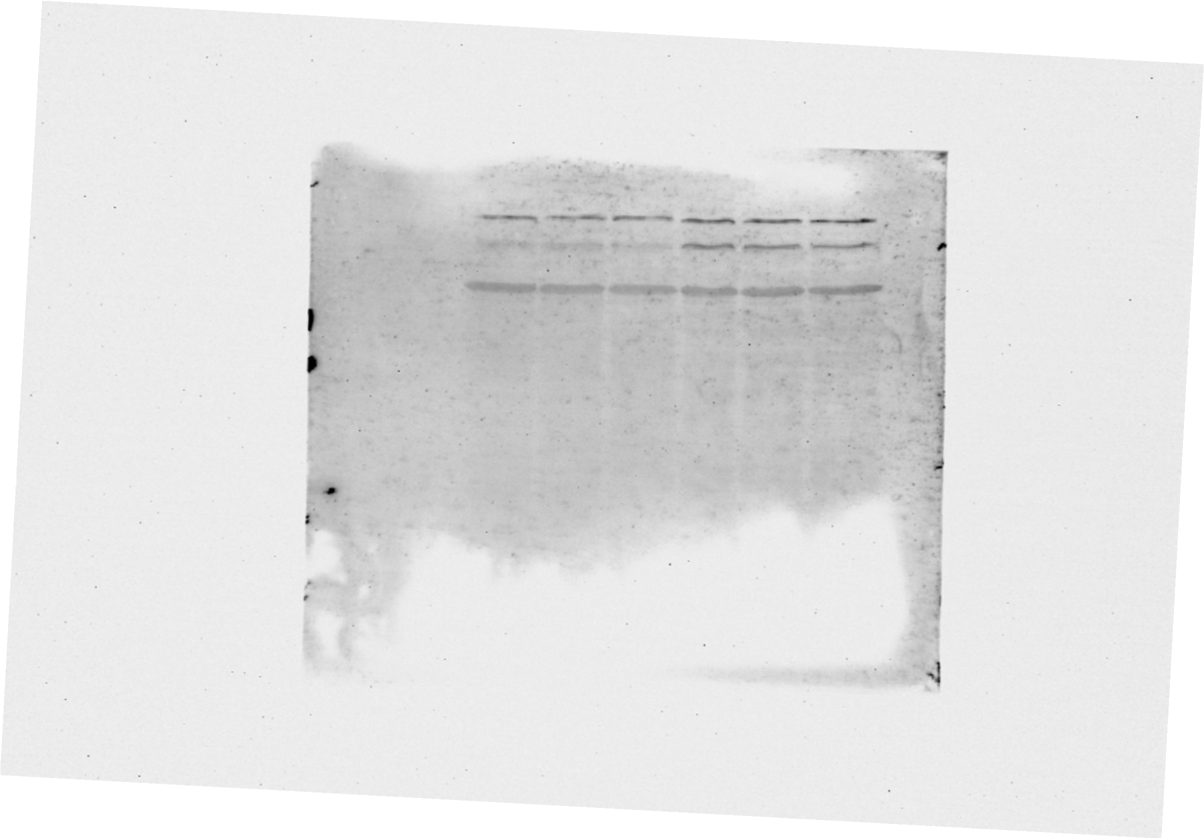

**Ponceau**

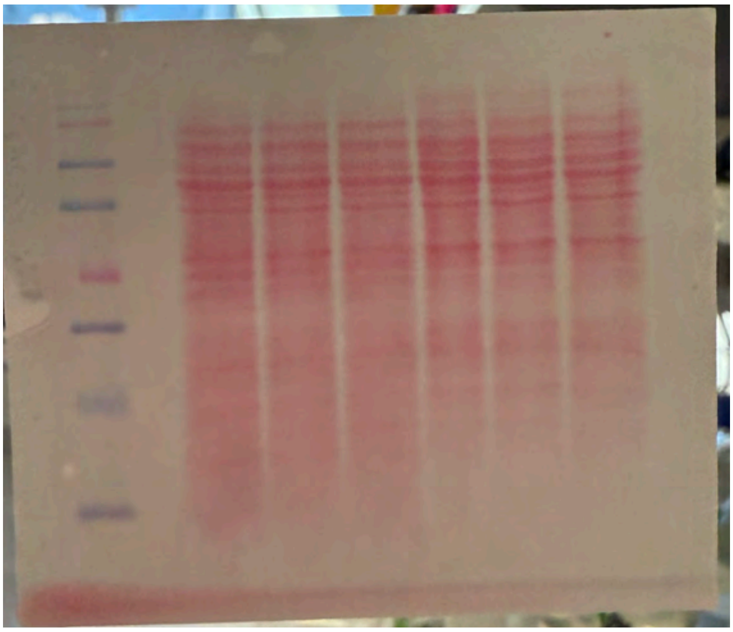
